# Starch granule initiation doesn’t require a starch synthase 4 isoform in *Chlamydomonas reinhardtii*

**DOI:** 10.1101/2024.10.03.616402

**Authors:** Adeline Courseaux, Philippe Deschamps, David Dauvillée

**Affiliations:** University Lille, CNRS, UMR 8576 - UGSF - Unité de Glycobiologie Structurale et Fonctionnelle, Lille, France; University Paris-Saclay, CNRS UMR 8079, AgroParisTech, Laboratoire Ecologie Systématique Evolution, Gif-sur-Yvette, France

**Keywords:** *Chlamydomonas reinhardtii*, Initiation, Starch, Starch phosphorylase, Starch synthase

## Abstract

The initiation of starch granule synthesis remains a relatively poorly understood phenomenon. Recent advances enabled the establishment of a model explaining the synthesis of new starch granules in *Arabidopsis thaliana*. These characterizations revealed the involvement of both a specific starch synthase isoform (SS4) and of several non-enzymatic proteins in this process.

In this work, we investigated whether the initiation of starch synthesis in the green microalga *Chlamydomonas reinhardtii* involves the same machinery as those uncovered in the plant model. Our extensive phylogenetic analysis revealed that most of the key players that were identified in higher plants are not found in microalgae suggesting that a different pathway is taking place. We showed that restoration of ADP-glucose synthesis in a mutant strain devoid of any endogenous primers allowed normal starch synthesis, revealing the existence of an initiation mechanism in Chlamydomonas. Our biochemical characterizations revealed that starch synthase isoform 3 possesses the intrinsic capacity to initiate polysaccharide synthesis *in vitro* and could be one of the functions involved in starch initiation. Our work suggests that the initiation of starch synthesis in Chlamydomonas involves a different pathway to that described in Arabidopsis and that further efforts will be required to identify the proteins involved in this process.

## 1. Introduction

Starch represents, after cellulose, the second most abundant natural polymer on Earth and is the principal source of carbohydrates in human and animal nutrition. The need to secure this food resource, as well as the development and increasing demand for biopolymer-derived renewable products, has prompted important research efforts to decipher the biological mechanisms of starch synthesis and degradation (Zeeman et al. 2010). Although generally considered as the hallmark storage polysaccharide of land plants, starch is actually synthesized by many other photosynthetic eukaryotes. The emergence of starch is one outcome of primary endosymbiosis, namely the definitive association of a cyanobacterium within a heterotrophic eukaryote, which gave rise to the primary plastid and to the Archaeplastida supergroup (*i.e.* red and green algae, glaucophytes and land plants) (Delwiche 1999; Adl et al. 2005). All Archaeplastida species accumulate starch that they synthesize thanks to a partially homologous metabolic pathway which is the result of the merging of cyanobacterial and eukaryotic glycogen metabolisms (Patron and Keeling 2005; Deschamps, Colleoni, et al. 2008). This composite pathway has been transmitted independently via secondary endosymbiosis to complex plastids bearing phyla like Cryptophyceae (Deschamps et al. 2006), Alveolates (Coppin et al. 2005) and Dinoflagellates (Deschamps et al. 2008; Dauvillée et al. 2009). Glycogen and starch share the same primary structure: a homopolymer of glucose residues connected together into linear glucans by α-1,4 bonds and ramified through α-1,6 linkages. However, these two polymers do not present the same physical properties: glycogen forms soluble and amorphous rosettes, while starch is accumulated in the form of highly organized semi-crystalline and insoluble granules. In Viridiplantae those granules are produced within the plastid, and in land plants, alternative mechanisms allow the accumulation of starch in the chloroplasts of photosynthetic tissues or in the amyloplasts of storage organs (Ball and Morell 2003). In source organs, starch is synthesized during periods of illumination as a sink for photosynthetically-fixed carbon, and is alternatively degraded at night to provide energy to the rest of the plant. Starch granules are composed of two subfractions, namely amylopectin and amylose, which greatly differ in their organizations. Amylose is mainly composed of linear glucans with very few ramifications (less than 1% α-1,6 linkages), while amylopectin, the major fraction of the granule (80% of the dry weight), is moderately branched (5% α-1,6 linkages) and exhibits a densely clustered organization responsible for the unique physicochemical properties and semi-crystalline state of starch (Manners 1989; Buléon et al. 1998).

The biosynthesis of glycogen requires few enzymatic steps. In bacteria, for instance, glycogen synthesis requires the action of only three different enzymes: (1) An ADP-glucose pyrophosphorylase (AGPase; EC 2.7.7.27) that activates glucose in the form of ADP-glucose; (2) A glycogen synthase (EC 2.4.1.21) that uses ADP-glucose to elongate linear glucans; and (3) a glycogen branching enzyme (BE, EC 2.4.1.18) that split existing linear chains and transfers the resulting cleaved chain to α-1,6 position. An important property of bacterial glycogen synthase is the ability to self-glucosylate and initiate *de novo* the synthesis of a new glucans, as demonstrated in the model bacteria *Agrobacterium tumefaciens* (Ugalde et al. 2003; Brust et al. 2013). Despite having the same primary structure, starch synthesis in plants occurs through a much more complex metabolic pathway compared to glycogen, both in terms of process and of regulation. Although the first steps are the same, the enzymes involved are more numerous: the AGPase is a heterotetramer containing both a catalytic and a regulatory subunit (Figueroa et al. 2022); there are two to three branching enzymes (Courseaux et al. 2023), and starch synthases have diversified into a complex family of isoenzymes. More precisely, if the synthesis of amylose only requires the action of the Granule Bound Starch Synthase (GBSS), the fine structure of amylopectin, which is composed of clusters of glucans of variable but very controlled lengths, is obtained through the interplay of several soluble starch synthases, each specialized in producing glucans of specific lengths (Szydlowski et al. 2011). The duplication and subfunctionalization of soluble starch synthases mainly predate the diversification of extant Viridiplantae. Indeed, the whole clade shares multiple ancient paralogs pointing to the same ancestral bacterial enzymes (Chang et al. 2023). One of the pivotal differences between glycogen and starch synthesis resides in the mechanism that allows toggling from amorphous to semi-crystalline amylopectin. This change of state, called pre-amylopectin trimming, is processed by a complex of specific isoforms of debranching enzymes called isoamylases (EC 3.2.1.68) (Ball et al. 1996; Dauvillée et al. 2001). These isoamylases remove branches in positions that prevent the formation of amylopectin clusters and consequently allow its insolubilization. Mutants lacking isoamylases substitute starch with an amorphous polysaccharide called phytoglycogen (Mouille et al. 1996). Similarly, to its biosynthesis, bacterial glycogen catabolism is quite a straightforward process involving only 2 enzymes: a glycogen phosphorylase and a debranching enzyme (Dauvillée et al. 2005; Alonso-Casajús et al. 2006). Conversely, starch degradation requires many more steps, including specific enzymes dedicated to the solubilization of amylopectin before amylolysis (Feike et al. 2016; Mahlow et al. 2016). The multi-enzymatic nature of starch metabolism is further complexified by the existence of multi-protein complexes that modulate the activity of the different enzymatic isoforms (Kötting et al. 2010; Ahmed et al. 2015; Mehrpouyan et al. 2021). Finally, the enzymes involved in starch metabolism may support different functions depending on the organism or the organ. For example, it was shown that one plastidial isoform of starch phosphorylase is not involved in building transitory starch but is nonetheless required for proper synthesis of storage starch (Zeeman et al. 2004; Dauvillée et al. 2006; Satoh et al. 2008). Altogether, the metabolism of starch in green algae and plants is a complex machinery involving, for what is known, more than twenty different enzymes and proteins.

The longstanding efforts in deciphering starch metabolism have now offered a relatively precise image of how the molecule is produced and degraded in plants. Conversely, the mechanism by which *de novo* synthesis of a new starch granule occurs has long remained poorly understood. This question encompasses two subaspects which are the initiation of the polymerization of new glucans on the one hand, and the nucleation of a new starch granule on the other hand. These two events, although conceptually different, are necessarily intertwined, but may not occur at the same place nor at the same time. If one refers to the initiation of glycogen particles, three alternative possibilities for starch initiation can be envisioned. First, as mentioned above, the glycogen synthase of bacteria can alone prime *de novo* a new glucan polymer, and ultimately a new glycogen particle. Starch synthases, however, have been so far reported only capable of elongating maltose or longer glucan primers (Brust et al. 2013). Secondly, in non-photosynthetic eukaryotes, the glycogen synthase lacks the priming ability and relies on the action of glycogenins (EC 2.4.1.186). Those enzymes are able to self-glucosylate and to offer a primer to the glycogen synthase to initiate the synthesis of a new glycogen particle. To date, no glycogenin-like enzyme has ever been characterized in plants, but other proteins supporting a similar function might potentially exist. The third scenario, which is the consequence of the first two, proposes that there might be no *de novo* priming at all, and that the presence of pre-existing glucans, or even polysaccharide structures, is required to start a new starch granule. Ral et al. (Ral et al. 2004), for example, described how the single starch granule of the prasinophyte alga *Ostreococcus tauri* is partitioned between daughter cells to ensure the presence of a nucleation structure after mitosis. Recent works, mainly conducted on the model plant *Arabidopsis thaliana*, have partially elucidated the granule initiation process, however only for the specific case of leaf transitory starch synthesis (Mérida and Fettke 2021). It has been demonstrated that the major enzyme responsible for the nucleation of new starch granules is the SS4-type starch synthase. Mutants deficient for the activity of SS4 no longer contain 4 to 7 lenticular starch granules per chloroplast, as usually found in wild-type *A. thaliana* plants, but typically exhibit a single, large, round-shaped granule (Roldán et al. 2007). This starch synthase isoform was also recently shown to be required for normal initiation in amyloplasts but appears to play different roles with respect to A-type and B-type granules (Hawkins et al. 2021). The SS4 belongs to the GT5 family according to the CAZy classification (Lombard et al. 2014) and displays two conserved C-terminal modules found in many other starch synthases: a GT1 glycosyltransferase domain as well as a GT5 domain (Liu et al. 2015). SS4 distinguishes from other synthase by a long N-terminal extension, containing coiled-coil domains and separated from the catalytic site by a small conserved region of about 50 amino acids, that is responsible for its particular intracellular localization and influences the shape of starch granules (Raynaud et al. 2016; Lu et al. 2018). Interestingly, double mutant lines of Arabidopsis defective for both SS4 and SS3 develop chloroplasts that are completely devoid of starch granules (Szydlowski et al. 2009). Moreover, the inactivation of SS5 in Arabidopsis also diminishes the number of starch granules synthesized per chloroplast (Abt et al. 2020). Thus, SS3 and SS5 might also be involved in starch initiation, either as partners for the nucleation, or as providers of glucans primers to the SS4, or could non-specifically substitute SS4 when it is missing.

Apart from starch synthases, several other proteins involved in starch initiation have recently been identified. First, Arabidopsis mutants deficient for PTST2 (Protein Targeting To Starch 2), a protein that directly interacts with starch synthase 4, exhibit a phenotype similar to *ss4* mutants, with chloroplasts containing a reduced number of starch granules (0 or 1 large granule per chloroplast (Seung et al. 2017). PTST2 is related to PTST1, a protein involved in the correct localization of the GBSS into the starch granule (Seung et al. 2015; Sharma et al. 2022). The overexpression of PTST2 leads to an increased number of starch granules per chloroplast, a phenomenon which was shown to be dependent on the presence of a functional SS4 (Seung et al. 2017). PTST2 contains a carbohydrate binding module (CBM48) suggesting that it could promote initiation by facilitating the accessibility of a suitable primer substrate to SS4. Additionally, PTST2 was shown to interact with two other coiled-coil-containing non-enzymatic plastidial proteins to form a putative initiation complex. These partners are MRC/PII1 (Myosin-Resembling Chloroplast protein; Protein Involved in starch Initiation 1) and MFP1 (MAR Binding Filament-like Protein 1) (Seung et al. 2018; Vandromme et al. 2019). Plants lacking MRC/PII1 in Arabidopsis harbor the same phenotype than the *ss4* mutant plants (*i.e.* few starch granules per chloroplasts) with the interesting difference that, unlike observed in *ss4* mutants, the morphology of the remaining granules is not altered. It was suggested that PII1/MRC would either help for the correct folding of SS4 or assist with the recruitment of other protein partners of a potential initiation complex (Vandromme et al. 2019). As mentioned above, in *ss4* mutant plants, a residual granule initiation activity is likely performed by the SS3 with low efficiency. When PII1 and SS4 mutations are combined, this residual SS3 priming activity is almost suppressed (Vandromme et al. 2023). The second coiled-coil protein of the complex, MFP1, was first proposed to be involved in the correct intracellular localization of SS4 (Seung et al. 2018) and later shown to be responsible of the sub-chloroplast localization of the starch granule initiation site (Sharma et al. 2024). A third paralog of the PTST protein family (PTST3) seems to be involved in starch initiation as well, although with only a minor role (Seung et al. 2017).

The complex formed by starch synthase 4 and its partners seems to handle the nucleation of new starch granules and to regulate their localization and morphology. It is, however, less clear how the initiation of new glucans occurs within the complex if no starch synthase has the ability to prime new glucans. Indeed, as mentioned above, starch synthases require at least a maltose primer for elongation (Brust et al. 2013). Additionally, starch phosphorylases, in the presence of an excess of glucose-1-phosphate, can elongate preexisting glucans only starting from maltotetraose or longer (Malinova et al. 2014). Double mutants of Arabidopsis combining a defect for PHS1 (the plastidial starch phosphorylase) together with the inactivation of either MEX1 (the plastidial maltose exporter) (Niittylä et al. 2004) or DPE2 (the cytosolic transglucosidase) (Chia et al. 2004; Fettke et al. 2006) harbor a dramatic decrease in starch granule number (Malinova et al. 2014; Malinova and Fettke 2017). These double mutant phenotypes are, however, only expressed when Arabidopsis plants are grown under a light-cycle that includes a dark phase, which suggests that a perturbation of starch degradation also has an effect on starch initiation. Additionally, it has been observed that, in a *ss4 ss3 amy3* triple mutant plants, most chloroplasts are devoid of starch granules except for a few that exhibit numerous granules, suggesting that a missing amylolytic activity can marginally restores starch initiation. These two observations suggest that malto-oligosaccharides (MOS) released during the starch degradation phase can be reused as a source of primers for the granule nucleation process. The degradation activities responsible for the release of those MOS might include disproportionating enzymes (DPE1 and DPE2), debranching enzymes (ISA) or amylases. Interestingly, in Arabidopsis, the disruption of the α-amylase 3 (AMY3) partially suppresses the phenotype of *ss4* mutants or of *ss3 ss4* double mutants. This indicates that, in wild type plants, AMY3 might also regulate starch initiation by degrading those MOS normally used by SS4/SS3 as primers for the synthesis of new starch granules.

Almost all progress made in understanding the mechanism of initiation of starch granules is restricted to the case of Arabidopsis leaves, and data concerning the initiation of starch in storage organs is much more limited. Nevertheless, SS4 seems to also be required for efficient starch granule initiation in wheat amyloplasts (Hawkins et al. 2021). Moreover, the starch phosphorylase PHS1 was recently shown to participate in the initiation of B-type granules in wheat amyloplasts, but not in that of the A-type granule sub-population (Kamble et al. 2023). Interestingly, the wheat mutant defective for PHS1 accumulates a regular amount and a normal number of starch granules per chloroplast in leaves, as observed in the equivalent Arabidopsis mutant (Malinova et al. 2014). On the contrary, it has been shown that the *ptst2* mutant plants display different phenotypes across the organs or organisms studied (Watson-Lazowski et al. 2022). While the MRC protein was shown to promote starch granule initiation in Arabidopsis leaves, the homologous protein appears to temporarily repress the synthesis of B-type starch granule initiation in the endosperm (Chen et al. 2024). Consequently, it seems reasonable to imagine that different processes could be involved in transitory or storage starch initiation, but also that the initiation machinery described in Arabidopsis leaves might not be universal in Viridiplantae, and that alternative initiation processes could exist in other organisms. In this work, we aimed at testing if starch initiation in *Chlamydomonas reinhardtii* also proceeds through the SS4 initiation complex. Chlamydomonas has been extensively used for more than two decades as a model organism to study starch metabolism in plants (Hicks et al. 2001). It offers many experimental advantages such as a short generation time, a haploid genome as well as a wide toolset for molecular biology and formal genetics. Moreover, the complete annotated sequences of its three genomes are available (Grant and Chiang 1980; Maul et al. 2002; Merchant et al. 2007) and an indexed insertional mutant library has been created, authorizing reverse genetics studies (Li et al. 2016). In Chlamydomonas, starch metabolism proceeds with the same enzymatic complexity compared to land plants (Deschamps et al. 2008), and many crucial elements of the pathway have been historically discovered using this alga (Mouille et al. 1996; Dauvillée et al. 2006). When grown under optimal conditions, Chlamydomonas synthesizes transitory starch around its pyrenoid, while under adverse conditions, the alga switches its metabolism to accumulate storage starch similar to the one found in roots or tubers of higher plants (Maddelein et al. 1994). As Chlamydomonas can mimic both transitory and storage starch metabolisms, we assessed the capacity of this microbial model at initiating starch granule synthesis. We first performed a genomic survey and phylogenetic analysis to test for the existence of proteins homologous to those described in Arabidopsis into the Chlamydomonas genome as well as a wide diversity of Viridiplantae. We show that most of those elements are absent in green algae, suggesting the existence of an alternative initiation process. Additionally, we demonstrate that the functional complementation of an algal mutant defective for the small subunit (catalytic) of the ADP-glucose pyrophosphorylase leads to a complete restoration of the previously depleted glucan metabolism, proving the existence of a *de novo* glucan synthesis process and suggesting that the availability of a pre-existing malto-oligosaccharide primer is not mandatory for starch synthesis. Finally, we tested the intrinsic ability of both starch synthases and starch phosphorylases at initiating polysaccharide synthesis *in vitro*. Our results suggest: (a) that the starch initiation complex described in Arabidopsis leaves is specific to Streptophyta and probably emerged at the time this clade diversified; and (b) that an alternative, probably ancestral, initiation pathway exists in Chlamydomonas and in other Chlorophytes, and that it very likely involves unknown components that still needs to be identified.

## 2. Material and Methods

### 2.1 Chemicals, Strains and culture conditions

All chemicals were purchased from Sigma-Aldrich (St. Louis, MO, USA) unless otherwise stated in the text.

The wild-type cell-wall less reference strain 330 (*mt+ arg7-7 cw15 nit1 nit2*) and its arginine autotrophic derivative starch mutant deficient for the small subunit of ADP-glucose pyrophosphorylase BafJ5 (*mt+ arg7-7 cw15 nit1 nit2 sta6-1::ARG7;* Zabawinski et al. 2001) were used in this study. The Chlamydomonas strains I152 defective for the starch synthase 3 isoform (*Cre04.g215150*) and I73 defective for the PhoB starch phosphorylase isoform (*Cre12.g552200*) were previously described in (Fontaine et al. 1993; Dauvillée et al. 2006). The insertional mutant strains from the CLiP library (Li et al. 2016) used in this study were *LMJ.RY0402.054236* and *LMJ.RY0402.227281* which contain respectively an insertion in the second intron of the *Cre04.g215150* locus encoding the starch synthase 1 isoform and in the first intron of the *Cre07.g336950* locus encoding the PhoA starch phosphorylase isoform.

Solid medium (Sueoka’s High Salts Acetate, HSA) and liquid culture media (Tris Acetate Phosphate, TAP) composition were described in (Sueoka 1960) and (Gorman and Levine 1965) respectively. Nitrogen free media were used to trigger massive starch accumulation both on Petri dishes (HSA-N) and liquid cultures (TAP-N) by replacing the nitrogen source by an equivalent concentration of NaCl.

### 2.2 ADP-glucose pyrophosphorylase assay

Crude extracts were prepared from late-log-phase cells (2.10^6^ cells ml^−1^) grown in TAP under continuous light (60 μE.m^−1^s^−1^). After centrifugation at 3000 g for 5 min, the cell pellets were immediately frozen in liquid nitrogen. The cells were disrupted by grinding in liquid nitrogen with a mortar and pestle in 2 ml of extraction buffer (80 mM Glycylglycine/NaOH pH 7.5, 20 mM MgCl_2_, 5 mM NaF, 5 mM dithiothreitol, 2 mM CaCl_2_, 10% glycerol, 3% PEG 6000). The extract was then centrifuged at 10 000g for 10 min at 4°C to remove the cell debris, and the supernatant was immediately used. The ADP-glucose pyrophosphorylase assay was performed as previously described (Ball et al. 1991). Briefly, ADP-glucose degradation by AGPase was assayed by measuring the amount of glucose-1-phosphate (G1P) formed after a 10 min incubation of crude extracts aliquots at 30°C in the extraction buffer supplemented with 1 mM sodium pyrophosphate, 0.05 mM glucose-1,6-bisphosphate, 0.25 mM NADP and 1 mM ADP-glucose. The AGPase activation factor was determined by following the same procedure except that the incubation buffer contained 1.5 mM 3-PGA. The amount of released G1P was monitored through the increase of absorbance at 340 nm due to the reduction of NADP^+^ after addition of rabbit muscle phosphoglucomutase (1 unit) and *Leuconostoc mesenteroides* glucose-6-phosphate dehydrogenase (1.5 unit).

### 2.3 Functional complementation

A 3.6 kb genomic fragment corresponding to the small subunit of AGPase (*Cre03.g188250*) was amplified from Chlamydomonas genomic DNA with the AGPaseFor (5’-G<u>GAATTC</u>ATGGCCCTGAAGATGCGGGTG-3’) and the AGPaseRev (5’-G<u>GAATTC</u>TTAGATGATGGTGCCGGC-3’) primers allowing the introduction of *Eco*RI restriction sites (underlined in the primer sequences) and cloned in the *Eco*RI site of the pSL18 vector (Dauvillée et al. 2006). The resulting complementation vector pSL-AGPase allows the expression of the AGPase small subunit under the control of both *psaD* 5’ and 3’ UTRs and contains the *aphVIII* gene conferring resistance to paromomycin (Sizova et al. 2001). This plasmid (1µg) was linearized by restriction with *Psi*I and introduced in the BafJ5 mutant strains by transformation using the glass beads method (Kindle 1990). The transformants were selected on TAP solid medium supplemented with 10 µg/mL paromomycin, purified on the same medium before being tested for the presence of starch though iodine staining.

### 2.4 Iodine staining of nitrogen-starved cell patches

Cell patches (20 µL) were incubated on nitrogen free medium (TAP-N) for 5 days under continuous light allowing massive starch accumulation and subsequently exposed to iodine vapor before being photographed.

### 2.5 Starch purification and polysaccharide assays

Liquid cultures were carried out under continuous light (40 µE.m^-2^.s^-1^) in the presence of acetate at 24°C and constant shaking (150 rpm). Late-log phase cultures (+N) were inoculated at 10^5^ cells mL^-1^ and harvested when the cell density reached 2 to 3.10^6^ cells mL^-1^. Nitrogen-starved cultures (-N) were inoculated at 5.10^5^ cells mL^-1^ and were harvested after 5 d at a final density of 1 to 2. 10^6^ cells mL^-1^. Cells were concentrated 100 times by centrifugation at 3000 g 10 min at 4°C and disrupted using a French Press at 10000 psi. The lysate was cleared by centrifugation at 10000 g for 10 min. The supernatant containing water-soluble polysaccharide (WSP) (including malto-oligosaccharides) was immediately boiled for 5 min to inactivate enzymatic activity. The starch contained in the pellet was purified by resuspension in 90% Percoll and centrifugation at 10000 g 10 min. Purified starch pellets were washed twice in ultrapure water to remove residual Percoll. Dilution of starch or boiled supernatant (containing water soluble polysaccharide) were assayed following the Enzytec starch kit instructions as previously described (Delrue et al. 1992).

### 2.6 Gel permeation chromatography

Starch granules (2 mg) were disrupted by incubation at 100°C 10 min in 200 µl DMSO. Amylopectin and amylose were recovered by precipitation with 800 µl ethanol and centrifugation 10 min at 10000g. The pellet was air dried 10 min and resuspended in 10 mM NaOH (300 µl) before being applied to a gel permeation chromatography column. Amylose and amylopectin were eluted from a Sepharose CL-2B column (0.5 cm inner diameter, 65 cm length) pre-equilibrated in 10 mM NaOH at a flow rate of 12 ml.min^-1^ and collected in 300µl fractions. The glucans were detected by mixing 80 µl of each fraction with 20 µl of iodine solution (0.1% I_2_, 1% KI) The absorbance of the polysaccharide/iodine complexes were measured each nm between 500 and 700 nm for each fraction allowing the determination of the wavelength at maximal absorbance (λ_max_).

### 2.7 Zymograms revealing starch synthase and starch phosphorylase activities

Crude extracts were prepared from late-log phase cultures. To detect starch synthase or starch phosphorylase isoforms, the crude extract proteins were separated on a 29:1 (acrylamide:bisacrylamide) 8% (w/v) native polyacrylamide gel containing 0.3% (w/v) rabbit liver glycogen and were separated at 15 V.cm^-1^ for 2h at 4°C in 25 mM Tris/glycine pH 8.3. The gels were incubated overnight in either 20 ml of starch synthase incubation mix (67 mM Glycyl-glycine/NaOH pH 9; 133 mM ammonium sulfate; 80 mM MgCl_2_; 0.6 mg.ml^-1^ bovine serum albumin; 25 mM β-mercaptoethanol; 2.4 mM ADP-glucose) or 20 ml of starch phosphorylase incubation mix (100 mM Sodium citrate/NaOH pH 7; 20 mM glucose-1-phosphate). The activities were detected by staining the gels in an iodine solution (0.1% I_2_, 1% KI). The capacity to initiate polysaccharide synthesis without any primer provided was tested with the same procedure except that the glycogen was omitted in the polyacrylamide gels. Starch synthase isoforms were also detected in denaturing conditions with the same setup except that the crude extract proteins were denatured before electrophoresis (100 °C, 5 min in the presence of 0.2 % SDS and 5 % β-mercaptoethanol), and SDS (0.1 %) was added to both the polyacrylamide gel and the electrophoresis buffer. After electrophoresis, the gels were washed 4 times 30 min in 40 mM Tris before incubation in the starch synthase mix.

### 2.8 Similarity searches, conserved domains and phylogenetic analyses

A custom local protein sequences database was assembled from various online sources and formatted for similarity searches using Blast. The database contains protein sequences predicted from the nuclear genomes and whole transcriptomes of 108 green algal and land plant species, covering the whole diversity of Archaeplastida as well as 708 genomes of prokaryotes. The composition of this database is described in supplementary table S1, Additional file 1. Reference sequences of Arabidopsis proteins known to be involved in starch initiation were used as queries with BlastP to search for similar proteins against our custom database. For each reference protein, hits were extracted into a multiple sequence file and used to produce draft phylogenetic trees as follows: sequence were aligned using mafft v7.205 with default parameters (Katoh and Standley 2013); alignments were trimmed using TrimAl v1.4.rev22 using the option ‘-gt 0.7’; trees were reconstructed using FastTree v2.1.7 (LG + Gamma + CAT20) (Price et al. 2010). These first trees were manually inspected and pruned to remove duplicate sequences as well as sub-groups of similar proteins that were not directly related to the homologous group of interest. Final trees, based on the refined selection of similar proteins, were reconstructed using Iqtree version 2.0.7 with parameters LG+C20+G+I, as well as 1000 ultrafast bootstrap pseudo-replicates (UFBoot) (Minh et al. 2013; Nguyen et al. 2014). Those trees were edited for illustration purposes using figtree version 1.4 (Rambaut 2009) and the vector graphics program Inkscape (https://inkscape.org/). Conserved domains were inferred using the SMART prediction server in normal mode as well as using the CDD server (Letunic et al. 2021; Wang et al. 2023). The resulting domain’s coordinates were downloaded and converted to SVG graphics using a custom python script.

## 3. Results

### 3.1 The starch granule initiation machinery identified in Arabidopsis is missing in Chlamydomonas and generally absent in green algae

As previously stated, functional studies on the initiation of the starch granule have mainly been focusing on transitory starch in Arabidopsis, and additional clues indicating the existence of the same mechanisms in other plants and algae are still scarce. To explore this question, we reconstructed the phylogeny of all proteins reported to be involved in starch granule initiation in Arabidopsis (see methods). Several phylogenetic trees focusing on starch synthase isoforms have been previously published (Liu et al. 2015; Chang et al. 2023). All Viridiplantae possess at least 4 starch synthases isoforms that can be divided in two subfamilies (SS1-2 and SS3-4-5, respectively noted SS-A and SS-B in (Chang et al. 2023)) that diversified from two different bacterial ancestral genes coding for glycogen synthases. Table 1 lists all loci encoding a putative or confirmed starch synthase in the Chlamydomonas genome. Although only two starch synthase activities can be detected on zymogram gels, it appears that the genome of *C. reinhardtii* contains seven isoforms of this enzyme: two SS1, one SS2, two SS3 and two other SS-like enzymes that do not belong to any conventional synthase families (noted SSX1 and SSX2 as in Chang et al. 2023. Figure 1 presents, to date, the most comprehensive maximum likelihood tree of all soluble starch synthases isoforms of the SS-B family. This tree, artificially rooted between starch synthases and the closest group of bacterial glycogen synthases, recovers all isoforms as strongly supported phylogenetic clades. A detailed version of the same tree is available in supplementary figure S2, Additional file 2. Importantly and contrary to what has been stated in previous publications (Mérida and Fettke, 2021), we show here that *C. reinhardtii*, as well as all other green algae, seems to lack both SS4 and SS5 isoforms, indicating that starch initiation in these organisms cannot be performed by these enzymes. The observed lack of SS4 and 5 is not only supported by the topology of the tree, but also by the absence in all chlorophyte genomes of any starch synthase with a domain composition similar to SS4 or SS5 (Figure 1 and supplementary figure S2, Additional file 2). We also, for the first time, highlight the existence of a second SS3 isoform in Chlamydomonas, which seems to be a common feature of all Chlorophyta (including Prasinophyceae). Streptophyta seem to only have one of these two ancient SS3 paralogs, although monocots have a specific, more recent, duplication of SS3. Interestingly, mutants affected at the *Cre06.g282000* locus (affiliated to SS3a in figure 1) show strong modifications in amylopectin structure and quantity, indicating that these two SS3 isoforms do not support fully redundant functions (Fontaine et al. 1993; Ral et al. 2004). Moreover, the strong conservation of those two isoforms across green algae implies an important role for both. Regarding the two additional synthases, SSX1 is shared by Chlorophyta and a subset of Streptophyta, and SSX2 is restricted to Chlamydomonadales. Although they seem to contain all the conserved domains necessary for their proper functioning, the role of these putative starch synthases remains undetermined. On the other hand, the phylogeny of PTST-like proteins (Supplementary figure S4, Additional file 2) shows, again contrary to previous publications (Seung et al. 2017), that PTST1, which is involved in the correct localization of the GBSSI in the starch granule, does in fact exist in Chlorophyta, and is not restricted to Streptophyta. PTST2 and 3 are, however, only found in the genomes of Streptophyta species. Although we could not properly root the PTST tree with an appropriate outgroup, a reasonable hypothesis would be that PTST2 and 3 both evolved from the duplication and sub-functionalization of PTST1. Finally, as previously reported (Seung et al. 2018), we could not find any gene corresponding to MFP1 and PII1 in the genomes of the Chlorophyta species that we explored (Supplementary figures S6 and S5, Additional file 2). Altogether, it seems that the starch initiation complex formed by SS4 and its partners SS5, PTST-2/3, MFP1 and PII1 is totally absent in green algae and is a specific innovation of Streptophytes. Consequently, we can speculate that starch initiation as described in Arabidopsis is probably not ancestral in Viridiplantae and that an alternative mechanism exists and needs to be identified in Chlorophyta.

**Figure 1:**
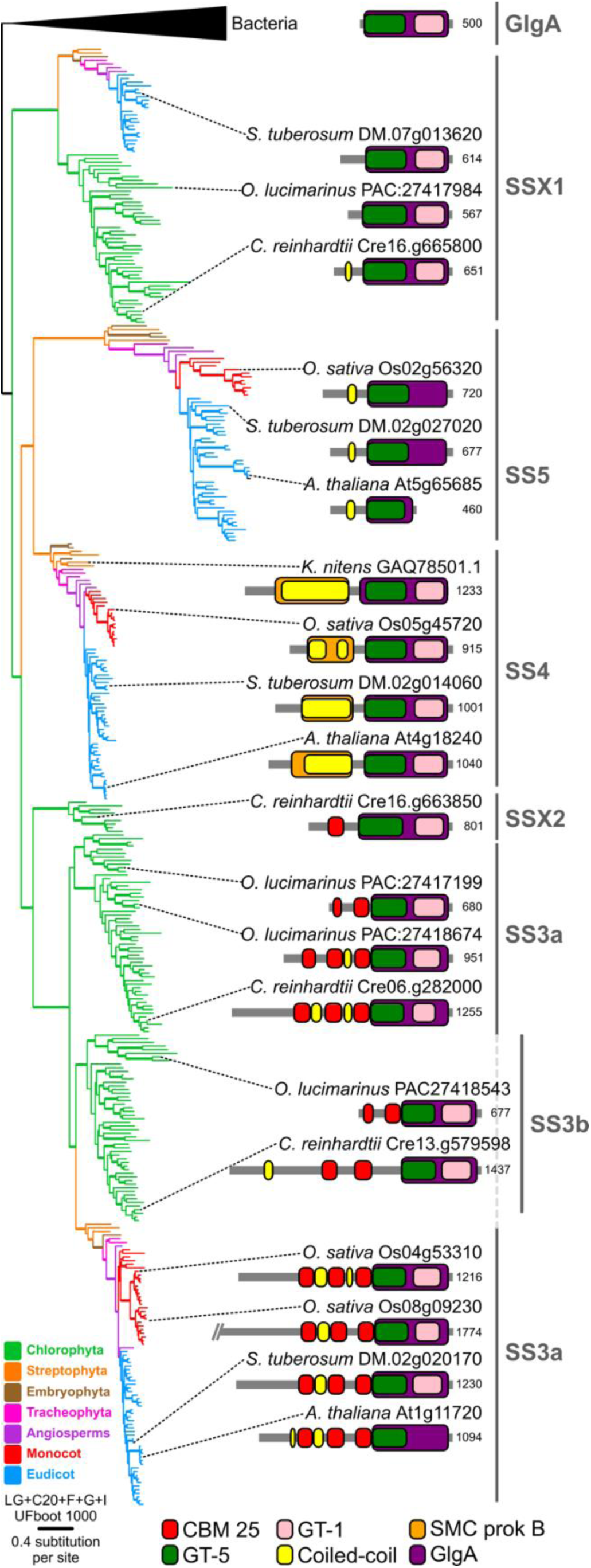
Phylogenetic and conserved domain analysis of starch synthases isoforms belonging to family SS3-4-5. The maximum-likelihood consensus tree on the left was reconstructed using IQ-tree and is artificially rooted between bacteria and viridiplantae. The name of each isoform familly/clade of starch synthase is provided on the right. Branch colors correspond to the taxonomy as described in the legend. Branch supports, estimated using 1000 UFBoot bootstrap pseudo-replicates, are depicted by a thick line if over or equal to 99%. Evolutionary model and branch length scale are also indicated. The corresponding detailed tree with all species names, protein references and support values is available in supplementary figure S1, Additional file 1. Sketches of conserved domain organization are provided for a selection of proteins belonging to each isoform; conserved domain names and colors are detailed in the legend; numbers correspond to the length of each protein.

**Table 1.**
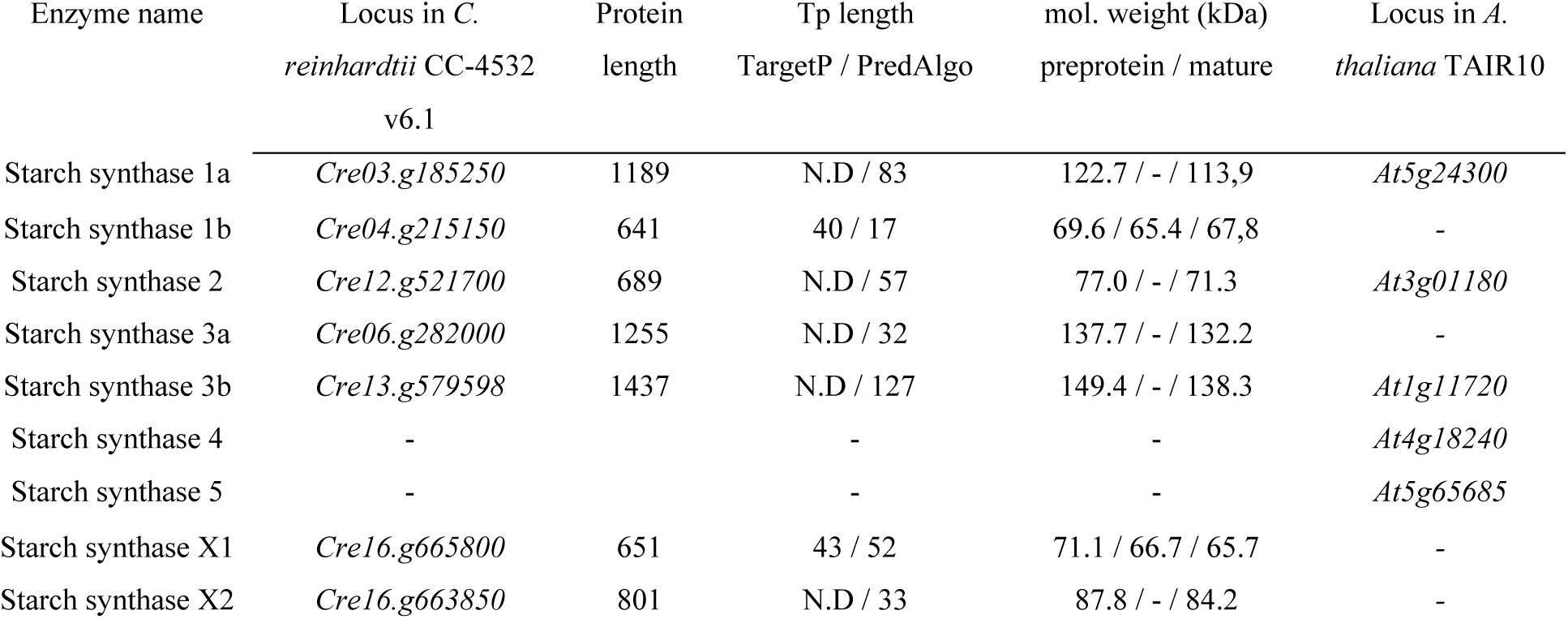
Gene loci encoding Chlamydomonas and Arabidopsis starch synthase isoforms. The genes were identified using the Phytozome (v13) website (Goodstein et al. 2012) based on the the genomes of *Chlamydomonas reinhardtii* CC-4532 (assembly version 6.1) and *Arabidopsis thaliana* (assembly version TAIR10). Loci names, protein lengths in amino-acids, lengths of transit peptides determined by both TargetP (Emanuelsson et al. 2000) and PredAlgo (Tardif et al. 2012) as well as the estimated molecular weight of the Chlamydomonas proteins are indicated.

| Enzyme name | Locus in *C. reinhardtii* CC-4532 v6.1 | Protein length | Tp length TargetP / PredAlgo | mol. weight (kDa) preprotein / mature | Locus in *A. thaliana* TAIR10 |
| --- | --- | --- | --- | --- | --- |
| Starch synthase 1a | *Cre03.g185250* | 1189 | N.D / 83 | 122.7 / - / 113.9 | *At5g24300* |
| Starch synthase 1b | *Cre04.g215150* | 641 | 40 / 17 | 69.6 / 65.4 / 67.8 | - |
| Starch synthase 2 | *Cre12.g521700* | 689 | N.D / 57 | 77.0 / - / 71.3 | *At3g01180* |
| Starch synthase 3a | *Cre06.g282000* | 1255 | N.D / 32 | 137.7 / - / 132.2 | - |
| Starch synthase 3b | *Cre13.g579598* | 1437 | N.D / 127 | 149.4 / - / 138.3 | *At1g11720* |
| Starch synthase 4 | - | - | - | - | *At4g18240* |
| Starch synthase 5 | - | - | - | - | *At5g65685* |
| Starch synthase X1 | *Cre16.g665800* | 651 | 43 / 52 | 71.1 / 66.7 / 65.7 | - |
| Starch synthase X2 | *Cre16.g663850* | 801 | N.D / 33 | 87.8 / - / 84.2 | - |

### 3.2 The complete depletion of MOS does not abolish the ability to initiate starch synthesis in Chlamydomonas

In the absence of a SS4 complex, a preliminary indirect approach to try to determine how starch initiation proceeds in green algae is simply to test if glucan synthesis can be processed *de novo*. Thus, we devised an experimental setup to assess the ability to process glucan synthesis in the absence of a glucan primer. The Chlamydomonas *sta6* mutant (strain BafJ5) was demonstrated to be deficient for the small catalytic subunit of ADP-glucose pyrophosphorylase, the only enzyme capable of synthesizing ADP-glucose in this alga (Zabawinski et al. 2001). We first ensured the total absence of starch as well as of any other soluble polysaccharides such as malto-oligosaccharides in this mutant. Cell patches of the isogenic wild-type strain and of the BafJ5 mutant were incubated on nitrogen-free medium to induce massive starch accumulation for five days. Upon iodine staining, the wild-type strain 330 turned black, revealing the presence of starch, while the BafJ5 mutant showed no interaction with iodine, proving the presence of very low if any amount of polysaccharide (Figure 2A). Iodine staining of total cells is, however, not sufficient to verify that residual quantities of glucans or very short glucans are absent in the mutant cells. Therefore, we assayed the quantity of polysaccharides sensitive to amyloglucosidase in the BafJ5 mutant and the isogenic wild-type strain 330 both in condition of massive starch synthesis (5 days under nitrogen depletion) or of transitory starch synthesis during exponential growth phase. Crude extracts of Chlamydomonas cells harvested in each condition were fractionated by centrifugation, and specific methods to assay α-glucans and glucose were used on both the purified insoluble starch pellet and on the supernatant (water soluble polysaccharide, malto-oligosaccharides, and glucose). Measurements reported in Table 2 show that the BafJ5 mutant does not contain any detectable starch amount in all growth conditions tested (the detection limit calculated in our experiments is 9 ng per million cells). No water-soluble polysaccharide was detected in both the wild-type and the mutant strains. The BafJ5 mutant being completely devoid of alpha-glucans, if Chlamydomonas possesses a glucan priming and a starch initiation system, restoring the synthesis of ADP-glucose should reinstate starch accumulation. Therefore, we cloned the structural gene of the small subunit of ADP-glucose pyrophosphorylase (*Cre03.g188250*), inserted it under the control of the strong *PsaD* promoter and used the construction to transform the BafJ5 mutant’s nuclear genome (see methods). The majority of the paromomycin-resistant strains tested (19 out of 24) displayed a grey/black staining with iodine indicating a restoration of starch synthesis (Figure 2A). Paromomycin resistant control clones obtained by the nuclear transformation of BafJ5 using an empty pSL18 vector did not display the complementation phenotype (6 clones tested, Figure 2A). We then picked 5 independent putatively complemented strains screened by iodine staining and assayed the amount of starch and soluble polysaccharides accumulated upon the two previously tested growth conditions (mixotrophy and after 5 days of nitrogen starvation, Table 2). For the five independent strains tested, we were unable to detect any soluble polysaccharide in both growth conditions. However, all strains accumulate insoluble polysaccharides with amounts ranging from 75 % to 122 % of the wild type reference under mixotrophic conditions and from 82 % to 159 % of the wild-type reference after nitrogen starvation. The ADP-glucose pyrophosphorylase activity in complemented strains was measured using specific assays and is comparable to that of the wild-type strain (Table 3). Additionally, the ADP-glucose pyrophosphorylase in complemented strains is sensitive to 3-phosphoglycerate (3-PGA) activation in a similar fashion to the one measured in the wild-type reference strain, indicating that the multimeric structure of the enzyme was also likely restored. Interestingly, the highest amounts of insoluble polysaccharides were observed in complemented strains for which the measured AGPase activity is also the highest (J5C2 and J5C4 with AGPase activity of respectively 200 and 170 % compared to the wild-type). This observation is in line with previous results obtained in land plants models, for which the amount of starch was shown to be tightly correlated to the size of the available ADP-glucose pool or the level of AGPase activity. As an example, a maize mutant where the AGPase was less sensitive to its negative effector (Pi) accumulates more starch in its endosperm than the wild-type plant (Giroux et al. 1996). Genetic manipulations of AGPase activity in both potato tuber and wheat endosperm led to an increase in starch synthesis as well (Stark et al. 1992; Smidansky et al. 2002). Moreover, we checked that insoluble polysaccharides accumulated in the 5 complemented strains display the typical starch structure and composition than the one found in the isogenic wild-type strain. Starches produced by cells harvested in exponential growth phase or after 5 days of nitrogen starvation were purified from the 5 complemented strains and the wild-type reference and were subjected to gel permeation chromatography (Figure 2B, supplementary figure S7, Additional file 2). Chromatograms obtained by analyzing the insoluble polysaccharide purified from the complemented strains display the same profile as wild-type starch, with the detection of the two starch subfractions amylopectin and amylose (Figure 2B). Moreover, starch granules accumulated by complemented strains display the same characteristics than those of wild-type cells, with an amylopectin λmax value ranging from 571 to 579 nm (average of 574.6 ± 2.9 nm) in mixotrophic growth conditions (WT = 575.2 ± 4.6 nm), and from 552 to 557 nm (554.2 ± 1.6 nm) after nitrogen starvation (WT = 552.4 ± 2.6 nm). The amylose relative content was also similar between the 5 complemented strains and the wild-type reference strain both under optimal growth conditions and after nitrogen starvation with respective values of 6.2 ± 2.9 % and 23 ± 5 % for the complemented strains, and 5.6 ± 2.5 % and 24 ± 6 % for the wild-type. These results demonstrate that reintroducing a functional copy of *Cre03.g188250* in the BafJ5 mutant leads to the restoration of starch synthesis and that *Chlamydomonas reinhardtii* necessarily possesses the capacity to initiate the synthesis of polysaccharides without pre-existing traces of glucans.

**Figure 2:**
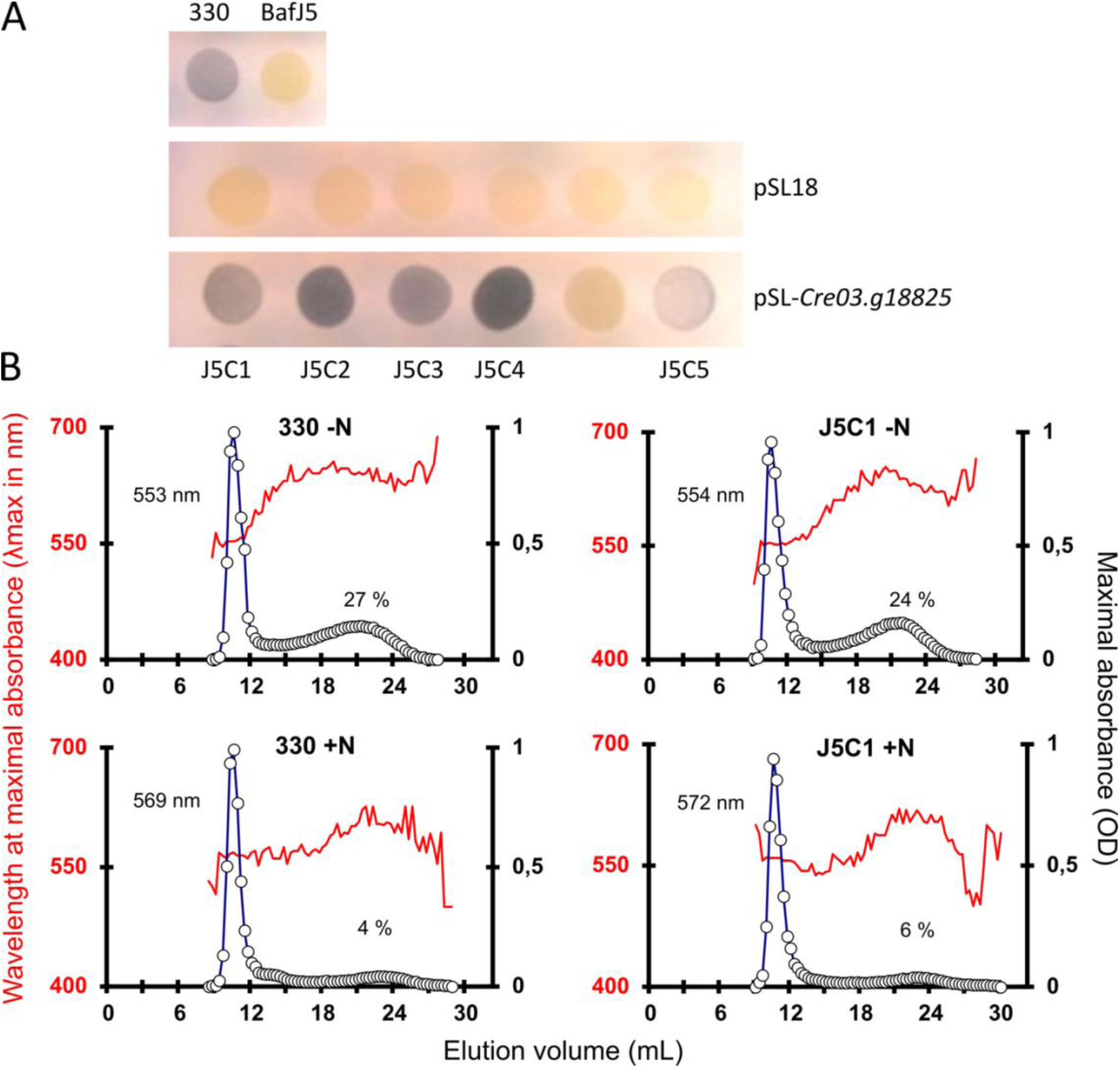
Functional complementation of the ADP-glucose pyrophosphorylase mutant strain. (A) Iodine screening. Chlamydomonas cell patches were incubated on nitrogen-starved medium for 5 days, allowing massive starch accumulation, and then stained with iodine vapor. Upper lane shows the staining of the wild-type strain 330 and the BafJ5 mutant strain defective for AGPase small subunit. Middle lane represents 6 paromomycin resistant strains obtained after transformation of BafJ5 with the pSL18 vector while the staining of 6 strains after transformation with the pSL-*Cre03.g188250* vector is displayed on the lower lane. Complemented strains harbor a grey/black phenotype due to starch accumulation. The brightness of the picture was increased to detect even tiny amount of polysaccharide interacting with iodine. The unmodified original picture is provided in additional file 3. (B) Separation of amylose and amylopectin on CL-2B gel permeation chromatography. The chromatograms obtained for two mg of starches produced by the wild-type (330) and the J5C1 complemented strain are displayed. The optical density (open circles) was measured for each fraction at λmax (red line in nm). Starches were purified from both late-growth phase (+N, lower panel) and nitrogen starved cultures (-N, upper panel). The amylose percentages in starch and the λmax of the pooled amylopectin fractions are indicated on each graph.

**Table 2:**
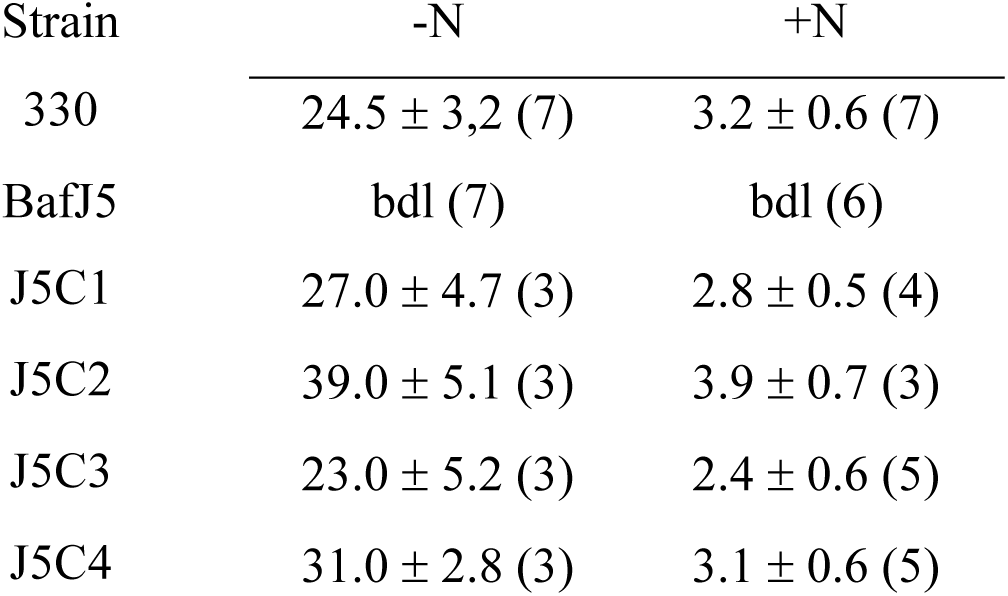
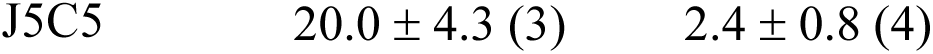
Phenotype of the strains studied during storage (-N) or transitory (+N) starch synthesis. Starch amounts accumulated after 5 days under nitrogen starvation (-N) or under mixotrophic conditions in the wild-type (330), the AGPase mutant (BafJ5) and 5 complemented strains (J5C) are indicated as average ± standard deviation in µg per million cells. The number of biological replicates is indicated between brackets. bdl: below detection limit calculated to be 9 ng per million cells with our experimental procedure.

**Table 3:** ADP-glucose pyrophosphorylase activity in the complemented strains. The assay was performed in the degradation direction by measuring the glucose-1-phosphate formed in the presence of 1 mM ADP-glucose and 1 mM pyrophosphate. The activation factor was calculated with the ratio of activities measured with or without 1.5 mM 3-PGA. The activities are expressed in nanomoles of glucose-1-phosphate formed per milligram of protein per minute. The results are expressed as the average ± SD from 3 biological replicates.

| Strain | -3PGA | +3PGA | Activation factor |
| --- | --- | --- | --- |
| 330 | $3.80 \pm 1.45$ | $12.20 \pm 5.82$ | 3.2X |
| BafJ5 | bdl | bdl | N/A |
| J5C1 | $2.90 \pm 0.87$ | $7.24 \pm 3.34$ | 2.5X |
| J5C2 | $7.80 \pm 3.10$ | $29.92 \pm 7.34$ | 3.8X |
| J5C3 | $3.95 \pm 2.12$ | $12.19 \pm 5.32$ | 3.1X |
| J5C4 | $6.40 \pm 3.23$ | $20.95 \pm 7.15$ | 3.3X |
| J5C5 | $2.30 \pm 2.09$ | $9.80 \pm 0.80$ | 4.2X |

### 3.3 Zymograms allow the visualization and genetic identification of four glucan elongating enzymes of *C. reinhardtii*

Although we demonstrated, using genetic evidence, that Chlamydomonas is able to synthesize glucans *de novo,* the underlying enzymes responsible for this process remain undetermined. Previous *in vitro* assays using recombinant Arabidopsis enzymes excluded the ability of starch synthases and starch phosphorylases to initiate glucan synthesis. We nevertheless tested the actual glucan priming capacity of those same two enzyme classes in Chlamydomonas. Starch synthase and phosphorylase activities can easily be revealed on zymograms, which consist on protein separation by electrophoresis in non-denaturing conditions using particular polyacrylamide gels followed by the incubation of the gel with the required enzyme substrates: ADP-glucose (ADPG) or glucose-1-phosphate (G1P) respectively. When a substrate like glycogen is embedded in the gel before polymerization, the enzymes will alter the polysaccharide during incubation, and those modifications can be revealed by staining the gel with iodine. In Chlamydomonas, zymograms performed on a glycogen-containing gel allow the detection, after incubation, of two distinct ADP-glucose-dependent starch synthase enzymatic activities (Figure 3A). The starch synthase isoform with the highest glycogen affinity (upper band) was previously demonstrated to be the product of the *Cre06.g282000* locus and corresponds to starch synthase 3 (SS3a in Figure 1). This enzyme is disrupted in the Chlamydomonas I152 mutant strain, where starch structure is altered, and which was used to demonstrate the genetic linkage between *Cre06.g282000* and the mutant phenotype (Fontaine et al. 1993; Ral et al. 2006); Figure 3B). The second starch synthase activity revealed on zymogram is probably the SS1 isoform which is encoded by the *Cre04.g215150* locus. There is no null mutant available that could confirm the identity of this enzymatic activity. Nonetheless, the apparent molecular weight of this enzyme around 68 kDa, determined on zymograms in denaturing conditions (Figure 3C), is compatible with the one predicted from the gene model after removal of the transit peptide at cleavage position 40-41 as determined by TargetP/ChloroP (Table 3) (Emanuelsson et al. 1999; Almagro Armenteros et al. 2019). Moreover, the insertional mutant *LMJ.RY0402.054236* available at the CLiP library (Li et al. 2016), and which contains an insertion in the second intron of the *Cre04.g215150* gene, displays a specific decrease in the intensity of the band corresponding to the fast-migrating isoform on starch synthases revealing zymograms (Figure 3D). From these converging observations, we may assume that the second zymogram activity corresponds to the gene product of the *Cre04.g215150* locus which encodes one of the starch synthase 1 isoforms, named SSIb in Table 1. A similar glycogen-containing native zymogram procedure allows the detection of two different starch phosphorylase activities after incubation with G1P. Both PhoA and PhoB show strong affinity to glycogen. The PhoB isoform was previously linked to the *Cre12.g552200* gene thanks to the genetic analysis of the I73 mutant in which the lower activity on zymogram is absent (Dauvillée et al. 2006 ; Figure 3E). The isoform migrating at the top of the gel (PhoA) corresponds to the *Cre07.g336950* gene. This could be confirmed by the analysis of the mutant strain *LMJ.RY0402.227281*, obtained from the CLiP library, which has an insertion in the first intron of the gene that induces the loss of the PhoA activity as observed on the zymogram in Figure 3E.

**Figure 3:**
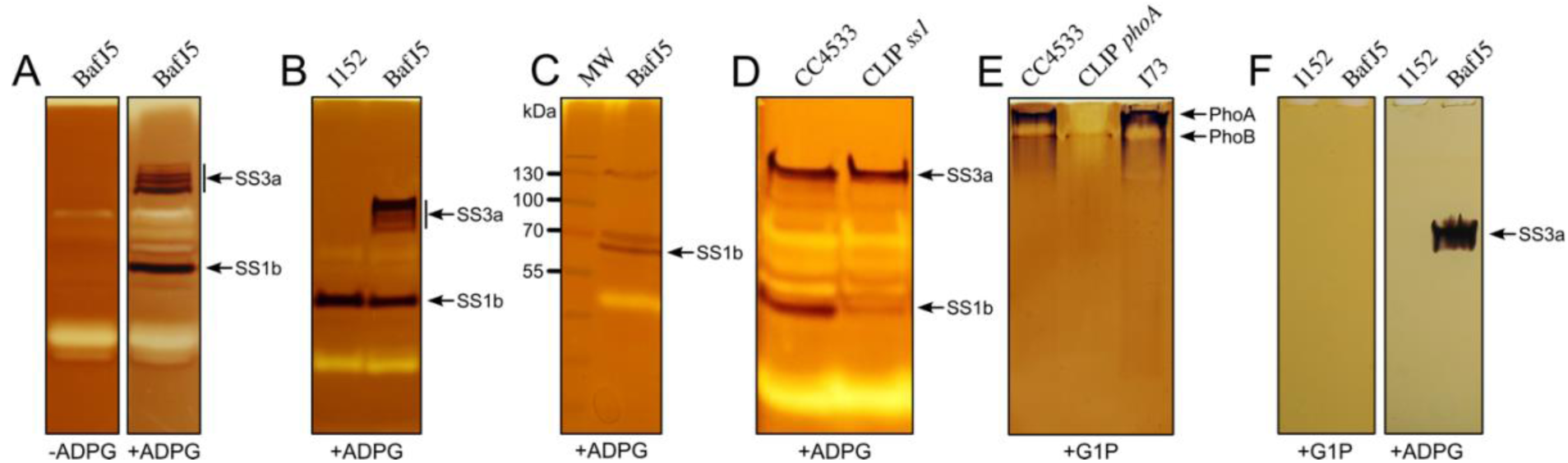
Detection of starch synthase and starch phosphorylase activities on zymograms. (A) Glycogen-containing zymogram revealing the starch synthase isoforms detected in the BafJ5 crude extracts (300µg) performed in denaturing conditions. After migration and renaturation, the gels were incubated in the starch synthase mix devoid of (right) or including (left) ADP-glucose and stained with iodine. Two activities able to elongate the glycogen outer chains and dependent on the presence of ADP-glucose are evidenced (indicated by arrows). (B) Three hundred µg of proteins from the BafJ5 strain and the I152 mutant defective for starch synthase 3 isoform (SS3a) were separated on a glycogen-containing zymogram in denaturing conditions. The gel was incubated overnight in the presence of ADP-glucose after proteins’ renaturation. (C) Three hundred µg of proteins from the BafJ5 strain were separated on a glycogen-containing zymogram in denaturing conditions in order to estimate the molecular weight of each starch synthase isoform. MW: Thermoscientific^TM^ PageRuler^TM^ prestained protein ladder. (D) The same procedure than in (B) was used to evidence the decrease in the activity band corresponding to the SS1 starch synthase isoform through the use of crude extracts of the CLiP mutant strain *LMJ.RY0402.054236* which contains an insertion in the second intron of *Cre04.g215150*. (E) Glycogen-containing zymogram revealing the starch phosphorylase isoforms detected in Chlamydomonas crude extracts. Two hundred µg of crude extract proteins were separated in native conditions at 4°C and the gel was incubated overnight in the presence of glucose-1-phosphate prior to iodine staining. CC4533: wild-type reference strain; CLiP *PhoA* corresponds to the CLiP mutant strain *LMJ.RY0402.227281* containing an insertion in the first intron of *Cre07.g336950*; I73: mutant strain defective for the PhoB isoform (Dauvillée et al., 2006). (F) Initiation of polysaccharide synthesis *in vitro*. Three hundred µg of crude extract proteins from the BafJ5 and the I152 mutant strains were separated by electrophoresis in native conditions on an 8% polyacrylamide gel devoid of any polysaccharide primers. After migration, the gels were incubated overnight in the starch phosphorylase mix containing glucose-1-phosphate (left) or the starch synthase mix containing ADP-glucose. The gels were subsequently immersed in iodine to reveal the synthesis of glucose chains. Only the starch synthase 3 isoform (SS3a) was shown to be able to produce a polysaccharide following this procedure.

### 3.4 Starch synthase 3 is able to synthesize *de novo* polysaccharide *in vitro*

As illustrated above, starch synthases and starch phosphorylases are easily detected on zymograms containing glycogen as a primer. This protocol, however, does not allow discriminating the initiation from the elongation capacity of those enzymes. To try to differentiate these two phenomena, we used zymogram gels containing no glycogen or any glucan substrates and tested the appearance of glucans after incubation with either ADPG or G1P. Figure 3F shows the result of this analysis for the BafJ5 and the I152 mutants. Without glycogen as a primer, neither PhoA, PhoB nor SS1 could produce glucans that could be detected by iodine staining. On the contrary, SS3a seems able to synthesize glucans in the absence of a primer, as observed using the BafJ5 protein extract, while the detected activity disappears when the I152 (SS3a) mutant protein extract is used. To ensure that our results are not falsified by the existence of contaminating glucan primers, we conducted several additional experiments where MOS were systematically eliminated by enzymatic pretreatments. First, we might suspect that the commercial pool of ADP-glucose used in our experiments is contaminated with MOS. Consequently, we incubated it with 10 units of porcine pancreatic amylase for 1h at room temperature before use. Figure 4A confirms that the initiation activity of the SS3 is still visible when using pretreated ADP-glucose, and is very likely not facilitated by the pre-existence of a glucan primer. Secondly, we cannot exclude that an endogenous primer could be already linked to the enzyme, we then tested the initiation capacity of SS3 after a step of denaturation/renaturation which would release this putative primer if present. Proteins from a crude extract of the AGPase mutant strain BafJ5 were separated on a SDS-PAGE in denaturing conditions. After electrophoresis, the gel was extensively washed with 40 mM Tris to allow the enzymes’ renaturation and the gel was incubated in the presence of amylase-pretreated ADP-glucose. The initiation and synthesis of polysaccharide by SS3 were still detected on the gels as evidenced by the interaction with iodine (Figure 4A). We also incubated the Chlamydomonas protein crude extracts in the presence of 0.5 units of amyloglucosidase prior to electrophoresis. Once again, only the starch synthase 3 isoform was able to initiate and elongate a polysaccharide in the gel performed in native conditions (Figure 4B). Up to 10 units of amyloglucosidase were used to treat the protein samples and even for the higher concentrations the polysaccharide synthesized by SS3 was still detected even if the latter was partially digested by amyloglucosidase which migrates at a close position to the starch synthase isoform in the gel (Figure 4C).

**Figure 4:**
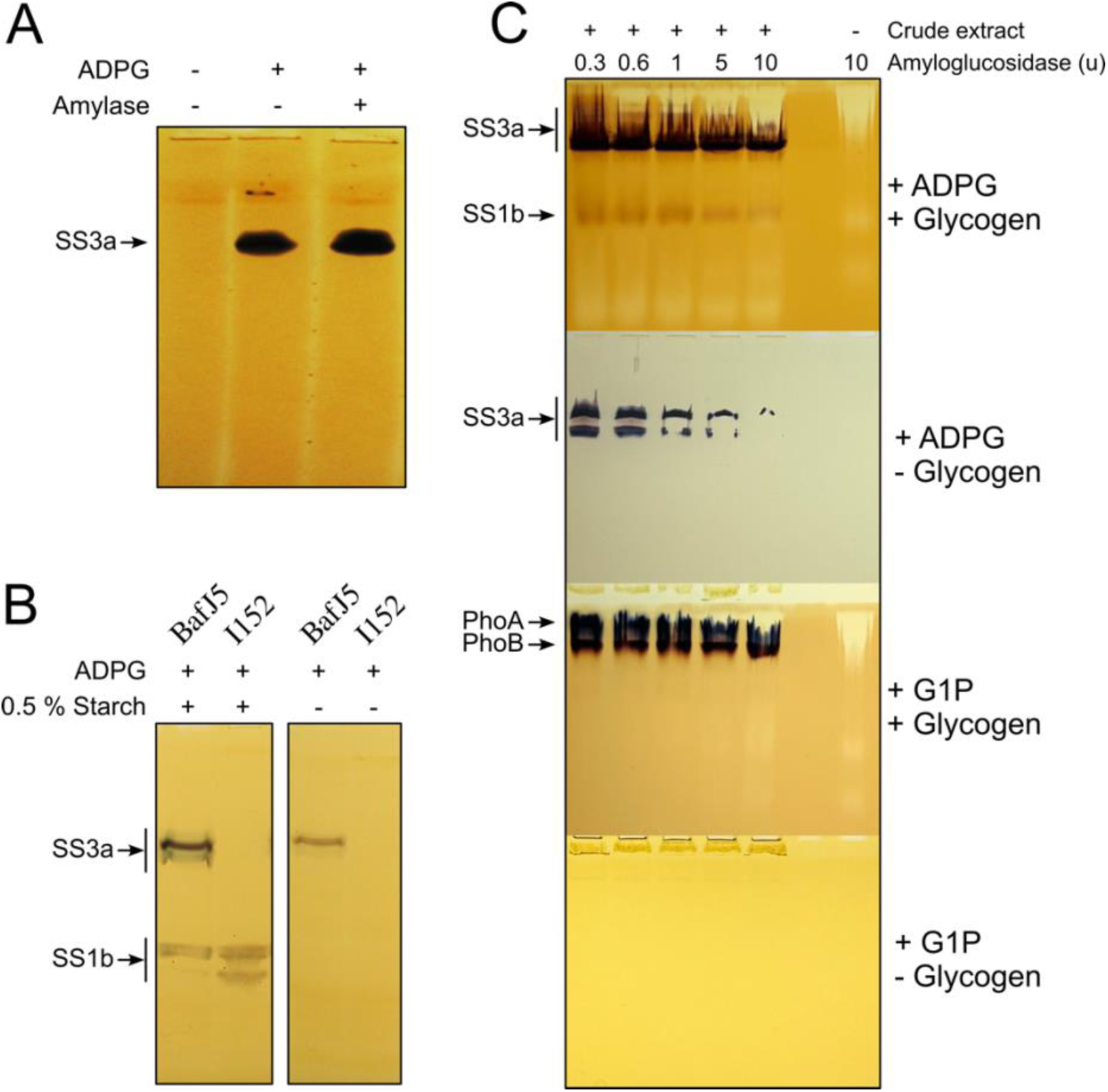
I*n vitro* capacity of starch synthase 3 isoform to initiate polysaccharide synthesis without pre-existing primer. (A) Three hundred µg of crude extract proteins from the BafJ5 mutant strain were separated by electrophoresis in native conditions on an 8% polyacrylamide gel devoid of any polysaccharide primers. Slices of the gel were incubated in a starch synthase mix containing either no ADP-glucose, 2.4 mM ADP-glucose or 2.4 mM amylase-pretreated ADP-glucose. (B) Three hundred µg of crude extract proteins from the BafJ5 and I152 mutant strains were pre-incubated with amyloglucosidase to remove any soluble α-glucans and were separated by electrophoresis in native conditions on an 8% polyacrylamide gel devoid of any polysaccharide primers. The gels were incubated in a starch synthase mix containing 2.4 mM amylase-pretreated ADP-glucose. The gel on the left was obtained with the same procedure except that soluble starch was added to the incubation mix providing a primer to both SS1 and SS3 isoforms. (C) Three hundred µg of crude extract proteins from the BafJ5 mutant strain were pre-incubated with different amounts (indicated at the top of the panel) of amyloglucosidase to remove any soluble α-glucans and were separated by electrophoresis in native conditions on 8% polyacrylamide gels. Gels used were either containing 0.3% rabbit liver glycogen or no substrate and were incubated either in the presence of amylase pre-treated ADP-glucose or glucose-1-phosphate as indicated on the right of the pictures.

## 4. Discussion

The mechanism of starch granule initiation in Arabidopsis has been slowly deciphered during the last 15 years. In Arabidopsis leaves, a protein complex precisely located in the chloroplast is responsible for the initiation of new starch granules. This complex is composed of the type 4 starch synthase combined with several non-enzymatic proteins that ensure the cellular localization of the complex, and may also provide the required glucan primers that the SS4 needs to initiate new glucan synthesis. There is only little experimental evidence for the involvement of the same complex in transitory starch synthesis in other plants, and even less concerning the initiation of granules in storage tissues. In this study, we provide the most taxon-resolved phylogenetic study of the members of this complex. We could observe that the proteins belonging to the SS4 initiation complex are present and conserved throughout the diversity of land plants. This result is an indirect but solid indication that the SS4 initiation complex might represent a canonical starch initiation machinery in Streptophyta. Our results also demonstrate that all green algae lack every component of this initiation complex. Indeed, both SS4 and SS5 are absent from all explored genomes of Chlorophytes, and are only detectable in Streptophytes including Charophyceae. This observation also applies to PTST2 and PTST3 as well as PII1, which all exist in Charophyceae and all other Streptophyte genomes explored, but not in Chlorophytes. Interestingly, the domain organization of the SS4 of Charophyceae comprises the specific coil-coiled domain (Figure 1), denoting that the molecular structures supporting the assembly of the complex already existed in the ancestor of all Streptophytes. The case of MFP1 is slightly different, as we could only detect this protein in Spermatophyta, which suggests that this specific component is a more recent refinement. The fact that the emergence of the SS4 starch initiation complex is somehow linked to the diversification of Streptophytes enables to speculate that the early evolution of multicellularity might have triggered the selection of a mechanism that would let the plant regulate where and when a new starch granule must be nucleated and stored. Although multicellularity arose multiple times in the evolution of Viridiplantae, only Streptophytes have developed cell specialization and complex cell-to-cell exchange and signaling (Umen 2014). In a context where cells begin to specialize, developing a solution to restrict starch storage localization could be advantageous in terms of energy use and cellular organization. For instance, several studies have shown the importance of starch in gravitropism, namely the ability with which plants detect and react to gravity. Indeed, mutants lacking or accumulating an excess of starch are affected in their perception of gravity and in their development (Kiss et al. 1997; Vitha et al. 2007). Gravity sensing is correlated with starch content; which inspired the hypothesis that directional growth in plants is regulated by a gradient of cell mass that depends on differential accumulation of starch (Morita 2010). Thus, the evolution of multicellularity and of directed growth in plants was likely linked to modifications of the starch metabolism that would allow the fine regulation of the number of starch granules per cell as well as their localization in the organism. In this context, the development of the SS4 initiation complex, which was demonstrated to tightly control the cellular localization and number of starch granules might have been central. The modality of glucan priming within the complex remains however not resolved, and no study so far has been able to uncouple glucan initiation from starch nucleation in any Streptophyte model organism.

The absence of the SS4 complex in Chlorophytes demonstrates, however, that it is not absolutely required for the initiation of glucan synthesis or starch granule nucleation, and that these two mechanisms probably originally relied on different enzymes that still need to be identified. Our study provides significant insights into the process of glucan and starch initiation in Chlamydomonas, contributing to a broader understanding of starch biosynthesis in microalgae. Starch synthases in green algae and plants have long been considered unable to initiate glucan synthesis *de novo* and require an extendable primer, as indicated by biochemical assays using recombinant proteins. However, no source of *de novo* oligosaccharides has ever been clearly identified in plants. For instance, even though the direct synthesis of maltose from glucose-1-phosphate has been measured by enzymatic assays on crude extracts, no maltose phosphorylase has ever been detected in a plant genome, and it is quite likely that this molecule might only be produced from the degradation of longer glucans (Schilling and Kandler 1975; Schilling 1982). Previous observation made in microalgae, specifically in the tiny prasinophyte *Ostreococcus tauri*, indicated that the unique starch granule present within the cell divides synchronously with cell division, ensuring that each daughter cell inherits a starch granule (Ral et al. 2004). Starch partitioning has been interpreted as the illustration that physical transmission of a glucan primer to daughter cells was a necessary mechanism that Viridiplantae developed to compensate for their inability to prime glucans. If this dead-end configuration was real, any event that would lead to the loss of glucans would be irrecoverable, which seems evolutionary very unlikely. We show here that, even after complete depletion of glucans, Chlamydomonas is still able to synthesize starch, meaning that there exists a mechanism for *de novo* glucan initiation. We also bring evidence that one specific isoform of starch synthase (SS3a) of Chlamydomonas is able, in our experimental setup, to generate new glucans in the absence of any pre-existing primer, suggesting that this enzyme might be involved in starch initiation. This contradicts previous studies, and leaves questions as to why previous assays using SS3 from other organisms concluded otherwise. One possibility is that, in our experiments, even if we ensured as much as possible the degradation of any contaminant, traces of glucans might have triggered elongation by the SS3. Alternatively, we can also speculate that the conformation of the enzyme is crucial for this function, and that a recombinant SS3 is not properly folded, thus failing to initiate in an assay. It’s however interesting to note that even after a cycle of denaturation-renaturation, the SS3a of Chlamydomonas is still able to initiate glucan synthesis. Finally, there is also the possibility that SS3 of green algae does not have the exact same properties as those of land plants.

Contrary to what has been previously published, our phylogenetic analyses highlight that all green algae share two isoforms of SS3, that we named SS3a and SS3b, while Streptophytes only have one. The SS3a is the only one of the two that is detectable on zymograms and for which we could evaluate the ability to synthesize glucans *de novo*. The mutant strains defective for this particular starch synthase 3 isoform are strongly affected in starch content and structure, but still accumulate starch (Fontaine et al. 1993; Ral et al. 2006). This indicates that SS3 isoforms are not strictly redundant enzymes, but also that SS3a is maybe not the only enzyme capable of glucan/starch initiation. This latest point remains not totally demonstrated however. Indeed, as it becomes clear that starch metabolism is a process of ever recycling MOS, it is imaginable that SS3a mutants never really stopped synthesizing glucans through the degradation of starch or the action of the other starch synthases using available primers. Thus, only after depleting SS3a mutants from any trace of glucans would one observe if the absence of SS3a only can prevent the restoration of starch synthesis. Experiments on SS3a / AGPase double mutants could offer more information on that point. On the other hand, evaluating the characteristics of SS3b and the effect of its absence in a mutant would be of particular interest. Although many campaigns of mutagenesis and screening for strains with altered starch phenotype have been conducted, no mutant affecting the SS3b gene has ever been isolated. Apart from a simple lack of chance, reasons might be that absence of SS3b in a strain still containing SS3a leads to a mild phenotype, or that, because SS3b is not detectable on zymograms, a potential mutant wouldn’t have passed in-depth screening. Insertional mutants potentially affecting this gene seem to be available in the CLiP library, and might provide an interesting material to push those investigations further. Our results, however, probably only reveal a portion of the real mechanism that allows starch initiation in Chlorophyta. Indeed, even if both SS3 enzymes are able to promote starch initiation by starting the synthesis of new glucans, it remains very likely that other enzymes or structural proteins should exist to regulate the initiation of starch granules and avoid random nucleation by the action of the SS3. Consequently, future work should focus on unraveling the molecular mechanisms underlying starch granule initiation in Chlamydomonas. Recently, we isolated the Chlamydomonas mutant *bsg1* that lacks a protein without apparent catalytic activity. This mutant exhibited a profound inability to initiate new starch granules when cells were transferred from mixotrophic growth conditions to nitrogen starvation. In the *bsg1* mutant, new stromal starch granules were hardly detected, and instead, pyrenoidal starch was utilized as a primer to sustain starch deposition. Interestingly, the Bsg1 protein contains both a coiled-coil domain and a CBM20 module which is closely related to the CBM48 family found in the PTST proteins (Findinier et al. 2019). These observations suggest that Bsg1 could play a crucial role in the initiation of starch granule formation. Identifying other elements that interact with this protein could be a route to uncover the components of the starch initiation machinery in *Chlamydomonas reinhardtii*.

## 5. Conclusion

In conclusion, by combining phylogenetics and experimental approaches, we were able to determine that Chlamydomonas and all other Chlorophytes do not possess an SS4 initiation complex and that they likely nucleate new starch granules by an alternate mechanism. This unknown pathway used by Chlorophytes might represent the ancestral starch initiation process as it existed before the diversification of Viridiplantae. The clear partitioning between Chlorophytes and Streptophytes with respect to the presence/absence of some isoforms of starch synthase as well as of the other components of the SS4-complex probably reflects a link between the evolution of new characters in Streptophytes that have benefited from developing a new way of regulating starch initiation, as we discussed above. Did the SS4 complex just replace the ancestral mechanism or do both still co-exist in Streptophytes, but with specialized actions in time or space? This question can only be addressed if the modalities of starch initiation in Chlorophytes are further investigated. More generally, a deeper understanding of the different steps leading to starch granule synthesis in microalgae is crucial, notably because these organisms are increasingly utilized as bioresources. For example, the production of bioplastics from algal starch presents a valuable alternative to conventional plastics (Six et al. 2024). This approach offers a sustainable solution that does not compete with plants in terms of arable land use or food supply, making it a promising avenue for reducing our reliance on fossil fuels and addressing environmental challenges.

## Credit authorship contribution statement

**Adeline Courseaux** : Investigation, validation. **Philippe Deschamps** : Conceptualization, Investigation, validation, Writing-original draft. **David Dauvillée** : Conceptualization, Investigation, validation, Writing-original draft.

## Data availability

The datasets used and/or analyzed during the current study are available from the corresponding author on reasonable request.

## Acknowledgments

This work has been financially supported by Centre National de la Recherche Scientifique France.

## Supplementary Data

Supplementary table S1 is available in Additional file 1.

Supplementary figures S1 to S7 are available in Additional file 2.

Original pictures used for figures are available in Additional file 3.

