## Supplemental table 1 for "Starch granule initiation doesn’t require a starch synthase 4 isoform in *Chlamydomonas reinhardtii*"

### Supplementary Table S1: Composition of the genomes database used for similarity searches.

| Species | Domain | Taxonomy | Assembly type | N. proteins | Assembly source | Source link |
|---|---|---|---|---|---|---|
| Acaryochloris marina MBIC11017 | Bacteria | Cyanobacteria\|Group E | genome | 8383 | NCBI | https://www.ncbi.nlm.nih.gov/genbank/ |
| Acaryochloris marina S15 | Bacteria | Cyanobacteria\|Group E | genome | 5833 | NCBI | https://www.ncbi.nlm.nih.gov/genbank/ |
| Acaryochloris sp. CCMEE 5410 | Bacteria | Cyanobacteria\|Group E | genome | 6776 | NCBI | https://www.ncbi.nlm.nih.gov/genbank/ |
| Acholeplasma laidlawii PG_8A | Bacteria | Firmicutes | genome | 1380 | NCBI | https://www.ncbi.nlm.nih.gov/genbank/ |
| Acidianus hospitalis W1 | Archaea | Crenarchaeota | genome | 2368 | NCBI | https://www.ncbi.nlm.nih.gov/genbank/ |
| Acidilobus saccharovorans 345-15 | Archaea | Crenarchaeota | genome | 1499 | NCBI | https://www.ncbi.nlm.nih.gov/genbank/ |
| Acidiphilium cryptum JF_5 | Bacteria | Proteobacteria\|Alphaproteobacteria\|Rhodospirillales | genome | 3559 | NCBI | https://www.ncbi.nlm.nih.gov/genbank/ |
| Acidithiobacillus ferrooxidans ATCC 2327 | Bacteria | Proteobacteria | genome | 3147 | NCBI | https://www.ncbi.nlm.nih.gov/genbank/ |
| Acidobacteria bacterium Ellin345 | Bacteria | Acidobacteria | genome | 4777 | NCBI | https://www.ncbi.nlm.nih.gov/genbank/ |
| Acidothermus cellulolyticus 11B | Bacteria | Actinobacteria | genome | 2157 | NCBI | https://www.ncbi.nlm.nih.gov/genbank/ |
| Aciduliprofundum boonei T469 | Archaea | Euryarchaeota | genome | 2949 | NCBI | https://www.ncbi.nlm.nih.gov/genbank/ |
| Acinetobacter baumannii AB0057 | Bacteria | Proteobacteria | genome | 3801 | NCBI | https://www.ncbi.nlm.nih.gov/genbank/ |
| Actinobacillus pleuropneumoniae L20 | Bacteria | Actinobacteria | genome | 2012 | NCBI | https://www.ncbi.nlm.nih.gov/genbank/ |
| Aeromonas hydrophila ATCC_7966 | Bacteria | Proteobacteria | genome | 4122 | NCBI | https://www.ncbi.nlm.nih.gov/genbank/ |
| Aeropyrum pernix K1 | Archaea | Crenarchaeota | genome | 1700 | NCBI | https://www.ncbi.nlm.nih.gov/genbank/ |
| Agrobacterium tumefaciens C58 | Bacteria | Proteobacteria\|Alphaproteobacteria\|Hyphomicrobiales | genome | 5360 | NCBI | https://www.ncbi.nlm.nih.gov/genbank/ |
| Akkermansia muciniphila ATCC BAA-835 | Bacteria | PVC\|Verrucomicrobia | genome | 2138 | NCBI | https://www.ncbi.nlm.nih.gov/genbank/ |
| Alcanivorax borkumensis SK2 | Bacteria | Proteobacteria | genome | 2755 | NCBI | https://www.ncbi.nlm.nih.gov/genbank/ |
| Aliivibrio salmonicida LFI1238 | Bacteria | Proteobacteria | genome | 3911 | NCBI | https://www.ncbi.nlm.nih.gov/genbank/ |
| Alkalilimnicola ehrlichei MLHE_1 | Bacteria | Proteobacteria | genome | 2865 | NCBI | https://www.ncbi.nlm.nih.gov/genbank/ |
| Alkaliphilus metalliredigens QYMF | Bacteria | Firmicutes | genome | 4625 | NCBI | https://www.ncbi.nlm.nih.gov/genbank/ |
| Alphaproteobacteria bacterium 33-17 | Bacteria | Proteobacteria\|Alphaproteobacteria\|Rickettsiales\|Rickettsiaceae | genome | 1687 | GTDB | https://gtdb.ecogenomic.org/ |
| Alphaproteobacteria bacterium CG11_GCA_002787615 | Bacteria | Proteobacteria\|Alphaproteobacteria\|Rickettsiales | genome | 1625 | GTDB | https://gtdb.ecogenomic.org/ |
| Alphaproteobacteria bacterium CG11_GCA_002787635 | Bacteria | Proteobacteria\|Alphaproteobacteria\|Rickettsiales | genome | 2227 | GTDB | https://gtdb.ecogenomic.org/ |
| Alphaproteobacteria bacterium sp002791805 | Bacteria | Proteobacteria\|Alphaproteobacteria\|MarineProteo1 | genome | 1391 | GTDB | https://gtdb.ecogenomic.org/ |
| Alphaproteobacteria bacterium sp003450915 | Bacteria | Proteobacteria\|Alphaproteobacteria\|MarineProteo1 | genome | 1535 | GTDB | https://gtdb.ecogenomic.org/ |
| Alphaproteobacteria bacterium sp003531345 | Bacteria | Proteobacteria\|Alphaproteobacteria\|Rickettsiales | genome | 1564 | GTDB | https://gtdb.ecogenomic.org/ |
| Alphaproteobacteria bacterium sp004293935 | Bacteria | Proteobacteria\|Alphaproteobacteria\|Rickettsiales | genome | 1660 | GTDB | https://gtdb.ecogenomic.org/ |
| Alphaproteobacteria bacterium UBA6149 | Bacteria | Proteobacteria\|Alphaproteobacteria\|Rickettsiales | genome | 2126 | GTDB | https://gtdb.ecogenomic.org/ |
| Alphaproteobacteria bacterium UBA6187 | Bacteria | Proteobacteria\|Alphaproteobacteria\|Rickettsiales | genome | 2299 | GTDB | https://gtdb.ecogenomic.org/ |
| Alteromonas macleodii Deep ecotype | Bacteria | Proteobacteria | genome | 4072 | NCBI | https://www.ncbi.nlm.nih.gov/genbank/ |
| Amborella trichopoda 291 v10 | Eukaryota | Viridiplantae | genome | 26846 | Phytozome | https://phytozome-next.jgi.doe.gov/ |
| Amoebophilus asiaticus 5a2 | Bacteria | Chlorobi | genome | 1283 | NCBI | https://www.ncbi.nlm.nih.gov/genbank/ |
| Anabaena azollae 0708 | Bacteria | Cyanobacteria\|Group B | genome | 3646 | NCBI | https://www.ncbi.nlm.nih.gov/genbank/ |
| Anabaena cylindrica PCC 7122 | Bacteria | Cyanobacteria\|Group B | genome | 6129 | JGI | https://genome.jgi.doe.gov/portal/ |
| Anabaena sp. 90 | Bacteria | Cyanobacteria\|Group B | genome | 4556 | NCBI | https://www.ncbi.nlm.nih.gov/genbank/ |
| Anabaena sp. PCC 7108 | Bacteria | Cyanobacteria\|Group B | genome | 5148 | JGI | https://genome.jgi.doe.gov/portal/ |
| Anabaena sp. WA102 | Bacteria | Cyanobacteria\|Group B | genome | 4890 | NCBI | https://www.ncbi.nlm.nih.gov/genbank/ |
| Anabaena sp. YBS01 | Bacteria | Cyanobacteria\|Group B | genome | 5632 | NCBI | https://www.ncbi.nlm.nih.gov/genbank/ |
| Anabaenopsis circularis NIES-21 | Bacteria | Cyanobacteria\|Group B | genome | 5642 | NCBI | https://www.ncbi.nlm.nih.gov/genbank/ |
| Anabaenopsis elenkinii CCIBt3563 | Bacteria | Cyanobacteria\|Group B | genome | 3542 | NCBI | https://www.ncbi.nlm.nih.gov/genbank/ |
| Anaerocellum thermophilum DSM 6725 | Bacteria | Firmicutes | genome | 2666 | NCBI | https://www.ncbi.nlm.nih.gov/genbank/ |
| Anaeromyxobacter sp. Fw109_5 | Bacteria | Myxobacteria | genome | 4466 | NCBI | https://www.ncbi.nlm.nih.gov/genbank/ |
| Ananas comosus 321 v3 | Eukaryota | Viridiplantae | genome | 27024 | Phytozome | https://phytozome-next.jgi.doe.gov/ |
| Anaplasma marginale StMaries | Bacteria | Proteobacteria\|Alphaproteobacteria\|Rickettsiales\|Anaplasmatace | genome | 948 | NCBI | https://www.ncbi.nlm.nih.gov/genbank/ |
| Anoxybacillus flavithermus WK1 | Bacteria | Firmicutes | genome | 2832 | NCBI | https://www.ncbi.nlm.nih.gov/genbank/ |
| Aquifex aeolicus VF5 | Bacteria | Aquificales | genome | 1560 | NCBI | https://www.ncbi.nlm.nih.gov/genbank/ |
| Aquilegia coerulea 322 v31 | Eukaryota | Viridiplantae | genome | 43550 | Phytozome | https://phytozome-next.jgi.doe.gov/ |
| Arabidopsis lyrata 384 v21 | Eukaryota | Viridiplantae | genome | 33132 | Phytozome | https://phytozome-next.jgi.doe.gov/ |
| Arabidopsis thaliana 167 TAIR10 | Eukaryota | Viridiplantae | genome | 35386 | Phytozome | https://phytozome-next.jgi.doe.gov/ |
| Arachis hypogaea 530 v10 | Eukaryota | Viridiplantae | genome | 84714 | Phytozome | https://phytozome-next.jgi.doe.gov/ |
| Archaeoglobus fulgidus DSM_4304 | Archaea | Euryarchaeota | genome | 2420 | NCBI | https://www.ncbi.nlm.nih.gov/genbank/ |
| Archaeoglobus profundus DSM 5631 | Archaea | Euryarchaeota | genome | 1823 | NCBI | https://www.ncbi.nlm.nih.gov/genbank/ |
| Archaeoglobus veneficus SNP6 | Archaea | Euryarchaeota | genome | 2090 | NCBI | https://www.ncbi.nlm.nih.gov/genbank/ |
| Arcobacter butzleri RM4018 | Bacteria | Proteobacteria\|Deltaproteobacteria | genome | 2259 | NCBI | https://www.ncbi.nlm.nih.gov/genbank/ |
| Aromatoleum aromaticum EbN1 | Bacteria | Proteobacteria\|Betaproteobacteria | genome | 4590 | NCBI | https://www.ncbi.nlm.nih.gov/genbank/ |
| Arthrobacter aurescens TC1 | Bacteria | Actinobacteria | genome | 4590 | NCBI | https://www.ncbi.nlm.nih.gov/genbank/ |
| Arthrospira maxima CS-328 | Bacteria | Cyanobacteria\|Group A | genome | 5431 | NCBI | https://www.ncbi.nlm.nih.gov/genbank/ |
| Arthrospira platensis C1 | Bacteria | Cyanobacteria\|Group A | genome | 4853 | NCBI | https://www.ncbi.nlm.nih.gov/genbank/ |
| Arthrospira platensis NIES-39 | Bacteria | Cyanobacteria\|Group A | genome | 5967 | NCBI | https://www.ncbi.nlm.nih.gov/genbank/ |
| Arthrospira platensis str. Paraca | Bacteria | Cyanobacteria\|Group A | genome | 4630 | NCBI | https://www.ncbi.nlm.nih.gov/genbank/ |
| Arthrospira platensis YZ | Bacteria | Cyanobacteria\|Group A | genome | 4890 | NCBI | https://www.ncbi.nlm.nih.gov/genbank/ |
| Arthrospira sp. PCC 9108 | Bacteria | Cyanobacteria\|Group A | genome | 4965 | NCBI | https://www.ncbi.nlm.nih.gov/genbank/ |
| Asparagus officinalis 498 V11 | Eukaryota | Viridiplantae | genome | 27395 | Phytozome | https://phytozome-next.jgi.doe.gov/ |
| Aulosira laxa NIES-50 | Bacteria | Cyanobacteria\|Group B | genome | 7088 | NCBI | https://www.ncbi.nlm.nih.gov/genbank/ |
| Azoarcus sp. BH72 | Bacteria | Proteobacteria\|Betaproteobacteria | genome | 3989 | NCBI | https://www.ncbi.nlm.nih.gov/genbank/ |
| Azorhizobium caulinodans ORS_571 | Bacteria | Proteobacteria\|Alphaproteobacteria\|Hyphomicrobiales | genome | 4717 | NCBI | https://www.ncbi.nlm.nih.gov/genbank/ |
| Bacillus amyloliquefaciens FZB42 | Bacteria | Firmicutes | genome | 3693 | NCBI | https://www.ncbi.nlm.nih.gov/genbank/ |
| Bacteroides fragilis NCTC_9343 | Bacteria | CFB | genome | 4231 | NCBI | https://www.ncbi.nlm.nih.gov/genbank/ |
| Bacteroides fragilis YCH46 | Bacteria | CFB | genome | 4625 | NCBI | https://www.ncbi.nlm.nih.gov/genbank/ |
| Bartonella bacilliformis KC583 | Bacteria | Proteobacteria\|Alphaproteobacteria\|Hyphomicrobiales | genome | 1283 | NCBI | https://www.ncbi.nlm.nih.gov/genbank/ |
| Bathycoccus prasinos | Eukaryota | Viridiplantae | transcriptome | 5479 | 1KP | https://db.cngb.org/onekp/ |
| Baumannia cicadellinicola Hc | Bacteria | Proteobacteria | genome | 595 | NCBI | https://www.ncbi.nlm.nih.gov/genbank/ |
| Bdellovibrio bacteriovorus HD100 | Bacteria | Proteobacteria\|Deltaproteobacteria | genome | 3587 | NCBI | https://www.ncbi.nlm.nih.gov/genbank/ |
| Beijerinckia indica subsp. Indica ATCC 9039 | Bacteria | Proteobacteria\|Alphaproteobacteria\|Hyphomicrobiales | genome | 3784 | NCBI | https://www.ncbi.nlm.nih.gov/genbank/ |
| Beta vulgaris ssp vulgaris 782 EL10 22 | Eukaryota | Viridiplantae | genome | 25326 | Phytozome | https://phytozome-next.jgi.doe.gov/ |
| Bifidobacterium adolescentis ATCC_15703 | Bacteria | Actinobacteria | genome | 1631 | NCBI | https://www.ncbi.nlm.nih.gov/genbank/ |
| Blastochloris viridis sp005768725 | Bacteria | Proteobacteria\|Alphaproteobacteria\|MarineProteo1 | genome | 1785 | GTDB | https://gtdb.ecogenomic.org/ |
| Blochmannia floridanus | Bacteria | Proteobacteria | genome | 583 | NCBI | https://www.ncbi.nlm.nih.gov/genbank/ |
| Boechera stricta 278 v12 | Eukaryota | Viridiplantae | genome | 29812 | Phytozome | https://phytozome-next.jgi.doe.gov/ |
| Bordetella bronchiseptica RB50 | Bacteria | Proteobacteria\|Betaproteobacteria | genome | 4994 | NCBI | https://www.ncbi.nlm.nih.gov/genbank/ |
| Borrelia afzelii PKo | Bacteria | Spirochaeta | genome | 1214 | NCBI | https://www.ncbi.nlm.nih.gov/genbank/ |
| Brachypodium distachyon 314 v31 | Eukaryota | Viridiplantae | genome | 52972 | Phytozome | https://phytozome-next.jgi.doe.gov/ |
| Brachyspira hyodysenteriae WA1 | Bacteria | Spirochaeta | genome | 2642 | NCBI | https://www.ncbi.nlm.nih.gov/genbank/ |
| Bradyrhizobium sp. BTAi1 | Bacteria | Proteobacteria\|Alphaproteobacteria\|Hyphomicrobiales | genome | 7622 | NCBI | https://www.ncbi.nlm.nih.gov/genbank/ |
| Brasilonema sennae CENA114 | Bacteria | Cyanobacteria\|Group B | genome | 6311 | NCBI | https://www.ncbi.nlm.nih.gov/genbank/ |
| Brassica rapa Fpsc 277 v13 | Eukaryota | Viridiplantae | genome | 43370 | Phytozome | https://phytozome-next.jgi.doe.gov/ |
| Brucella abortus 9_941 | Bacteria | Proteobacteria\|Alphaproteobacteria\|Hyphomicrobiales | genome | 3085 | NCBI | https://www.ncbi.nlm.nih.gov/genbank/ |
| Bryopsis plumosa | Eukaryota | Viridiplantae | transcriptome | 7322 | 1KP | https://db.cngb.org/onekp/ |
| Buchnera aphidicola APS | Bacteria | Proteobacteria | genome | 555 | NCBI | https://www.ncbi.nlm.nih.gov/genbank/ |
| Burkholderia sp. 383 | Bacteria | Proteobacteria\|Betaproteobacteria | genome | 7717 | NCBI | https://www.ncbi.nlm.nih.gov/genbank/ |
| Caldiarchaeum subterraneum | Archaea | Thaumarchaeota | genome | 3235 | NCBI | https://www.ncbi.nlm.nih.gov/genbank/ |
| Caldicellulosiruptor saccharolyticus DSM_8903 | Bacteria | Firmicutes | genome | 2679 | NCBI | https://www.ncbi.nlm.nih.gov/genbank/ |
| Caldivirga maquilingensis IC_167 | Archaea | Crenarchaeota | genome | 1963 | NCBI | https://www.ncbi.nlm.nih.gov/genbank/ |
| Calothrix brevissima NIES-22 | Bacteria | Cyanobacteria\|Group B | genome | 6929 | NCBI | https://www.ncbi.nlm.nih.gov/genbank/ |
| Calothrix parasitica NIES-267 | Bacteria | Cyanobacteria\|Group B | genome | 6996 | NCBI | https://www.ncbi.nlm.nih.gov/genbank/ |

Supplementary Table S1: Composition of the genomes database used for similarity searches.

| Species | Domain | Taxonomy | Assembly type | N. proteins | Assembly source | Source link |
|---|---|---|---|---|---|---|
| Calothrix sp. NIES-2098 | Bacteria | Cyanobacteria\|Group B | genome | 7032 | NCBI | https://www.ncbi.nlm.nih.gov/genbank/ |
| Calothrix sp. NIES-2100 | Bacteria | Cyanobacteria\|Group B | genome | 7563 | NCBI | https://www.ncbi.nlm.nih.gov/genbank/ |
| Calothrix sp. NIES-3974 | Bacteria | Cyanobacteria\|Group B | genome | 4526 | NCBI | https://www.ncbi.nlm.nih.gov/genbank/ |
| Calothrix sp. NIES-4071 | Bacteria | Cyanobacteria\|Group B | genome | 9831 | NCBI | https://www.ncbi.nlm.nih.gov/genbank/ |
| Calothrix sp. NIES-4105 | Bacteria | Cyanobacteria\|Group B | genome | 9824 | NCBI | https://www.ncbi.nlm.nih.gov/genbank/ |
| Calothrix sp. PCC 6303 | Bacteria | Cyanobacteria\|Group B | genome | 5719 | JGI | https://genome.jgi.doe.gov/portal/ |
| Calothrix sp. PCC 7103 | Bacteria | Cyanobacteria\|Group B | genome | 10042 | JGI | https://genome.jgi.doe.gov/portal/ |
| Calothrix sp. PCC 7507 | Bacteria | Cyanobacteria\|Group B | genome | 6132 | JGI | https://genome.jgi.doe.gov/portal/ |
| Campylobacter concisus 13826 | Bacteria | Proteobacteria\|Deltaproteobacteria | genome | 1985 | NCBI | https://www.ncbi.nlm.nih.gov/genbank/ |
| Candidatus Atelocyanobacterium thalassa isolate ALOHA | Bacteria | Cyanobacteria\|Group C | genome | 1144 | NCBI | https://www.ncbi.nlm.nih.gov/genbank/ |
| Candidatus Blackallbacteria bacterium CG13 | Bacteria | Melainabacteria | genome | 4914 | GTDB | https://gtdb.ecogenomic.org/ |
| Candidatus Caenarcaniphilales bacterium DT_14 | Bacteria | Melainabacteria | genome | 2855 | GTDB | https://gtdb.ecogenomic.org/ |
| Candidatus Caenarcanum bioreactoricola UASB_169 | Bacteria | Melainabacteria | genome | 1860 | GTDB | https://gtdb.ecogenomic.org/ |
| Candidatus Gastranaerophilus phascolarctosicola Zag_221 | Bacteria | Melainabacteria | genome | 1799 | GTDB | https://gtdb.ecogenomic.org/ |
| Candidatus Magnetaquicoccus inordinatus str | Bacteria | Proteobacteria\|Alphaproteobacteria\|Magnetococcales | genome | 3742 | GTDB | https://gtdb.ecogenomic.org/ |
| Candidatus Melainabacteria bacterium AlinenLipids_bin-9350 | Bacteria | Melainabacteria | genome | 3466 | GTDB | https://gtdb.ecogenomic.org/ |
| Candidatus Melainabacteria bacterium GCA_013298875 | Bacteria | Melainabacteria | genome | 2321 | GTDB | https://gtdb.ecogenomic.org/ |
| Candidatus Melainabacteria bacterium LMEP_10873 | Bacteria | Melainabacteria | genome | 2120 | GTDB | https://gtdb.ecogenomic.org/ |
| Candidatus Melainabacteria bacterium UBA8530 | Bacteria | Melainabacteria | genome | 3506 | GTDB | https://gtdb.ecogenomic.org/ |
| Candidatus Methanoregula boonei 6A8 | Archaea | Euryarchaeota | genome | 2450 | NCBI | https://www.ncbi.nlm.nih.gov/genbank/ |
| Candidatus Micrarchaeum acidiphilum ARMAN-2 | Archaea | Euryarchaeota | genome | 1037 | NCBI | https://www.ncbi.nlm.nih.gov/genbank/ |
| Candidatus Midichloria mitochondrii.IricVA | Bacteria | Proteobacteria\|Alphaproteobacteria\|Rickettsiales\|Midichloriaceae | genome | 1372 | GTDB | https://gtdb.ecogenomic.org/ |
| Candidatus Nanosalina sp. J07AB43 | Archaea | Euryarchaeota | genome | 1677 | NCBI | https://www.ncbi.nlm.nih.gov/genbank/ |
| Candidatus Nanosalinarum sp. J07AB56 | Archaea | Euryarchaeota | genome | 1407 | NCBI | https://www.ncbi.nlm.nih.gov/genbank/ |
| Candidatus Neoehrlichia lotoris str.RAC413 | Bacteria | Proteobacteria\|Alphaproteobacteria\|Rickettsiales\|Anaplasmatacea | genome | 951 | GTDB | https://gtdb.ecogenomic.org/ |
| Candidatus Nitrosoarchaeum koreensis MY1 | Archaea | Euryarchaeota | genome | 1945 | NCBI | https://www.ncbi.nlm.nih.gov/genbank/ |
| Candidatus Nitrosoarchaeum limnia SFB1 | Archaea | Euryarchaeota | genome | 2038 | NCBI | https://www.ncbi.nlm.nih.gov/genbank/ |
| Candidatus Nitrososphaera gargensis Ga9.2 | Archaea | Thaumarchaeota | genome | 3565 | NCBI | https://www.ncbi.nlm.nih.gov/genbank/ |
| Candidatus Nitrosotenuis uzonensis | Archaea | Thaumarchaeota | genome | 1944 | NCBI | https://www.ncbi.nlm.nih.gov/genbank/ |
| Candidatus Obscuribacter phosphatis EBPR_351 | Bacteria | Melainabacteria | genome | 4612 | GTDB | https://gtdb.ecogenomic.org/ |
| Candidatus Parvarchaeum acidiphilum ARMAN-4 | Archaea | Euryarchaeota | genome | 911 | NCBI | https://www.ncbi.nlm.nih.gov/genbank/ |
| Candidatus Parvarchaeum acidophilus ARMAN-5 | Archaea | Euryarchaeota | genome | 1042 | NCBI | https://www.ncbi.nlm.nih.gov/genbank/ |
| Candidatus Phytoplasma asteris AYWB | Bacteria | Firmicutes | genome | 693 | NCBI | https://www.ncbi.nlm.nih.gov/genbank/ |
| Candidatus Sericytochromatia bacterium LF-bin-346 | Bacteria | Melainabacteria | genome | 5282 | GTDB | https://gtdb.ecogenomic.org/ |
| Candidatus Sericytochromatia bacterium LF-bin-423 | Bacteria | Melainabacteria | genome | 3982 | GTDB | https://gtdb.ecogenomic.org/ |
| Candidatus Sericytochromatia bacterium S15B-MN24_CBMW_12 | Bacteria | Melainabacteria | genome | 3603 | GTDB | https://gtdb.ecogenomic.org/ |
| Candidatus Similichlamydia laticola str | Bacteria | PVC\|Chlamydiae\|Parilichlamydiaceae | MAG | 720 | GTDB | https://gtdb.ecogenomic.org/ |
| Candidatus Synechococcus calcipolaris | Bacteria | Cyanobacteria\|Group E | genome | 3257 | NCBI | https://www.ncbi.nlm.nih.gov/genbank/ |
| Candidatus Vampirovibrio chlorellavorus NCIMB_11383 | Bacteria | Melainabacteria | genome | 2816 | GTDB | https://gtdb.ecogenomic.org/ |
| Candidatus Xenolissoclinum pacificiensis L6 | Bacteria | Proteobacteria\|Alphaproteobacteria\|Rickettsiales\|Anaplasmatacea | genome | 928 | GTDB | https://gtdb.ecogenomic.org/ |
| Capsella grandiflora 266 v11 | Eukaryota | Viridiplantae | genome | 26561 | Phytozome | https://phytozome-next.jgi.doe.gov/ |
| Carboxydothermus hydrogenoformans Z_2901 | Bacteria | Firmicutes | genome | 2620 | NCBI | https://www.ncbi.nlm.nih.gov/genbank/ |
| Carsonella ruddii PV | Bacteria | Proteobacteria | genome | 182 | NCBI | https://www.ncbi.nlm.nih.gov/genbank/ |
| Caulobacter crescentus CB15 | Bacteria | Proteobacteria\|Alphaproteobacteria\|Caulobacterales | genome | 3737 | NCBI | https://www.ncbi.nlm.nih.gov/genbank/ |
| Cellvibrio japonicus Ueda107 | Bacteria | Proteobacteria | genome | 3754 | NCBI | https://www.ncbi.nlm.nih.gov/genbank/ |
| Cenarchaeum symbiosum A | Archaea | Thaumarchaeota | genome | 2017 | NCBI | https://www.ncbi.nlm.nih.gov/genbank/ |
| Ceratodon purpureus GG1 539 v11 | Eukaryota | Viridiplantae | genome | 39339 | Phytozome | https://phytozome-next.jgi.doe.gov/ |
| Ceratopteris richardii 676 v21 | Eukaryota | Viridiplantae | genome | 75253 | Phytozome | https://phytozome-next.jgi.doe.gov/ |
| Chamaesiphon minutus PCC 6605 | Bacteria | Cyanobacteria\|Group A | genome | 5813 | NCBI | https://www.ncbi.nlm.nih.gov/genbank/ |
| Chara_braunii_S276 | Eukaryota | Viridiplantae | genome | 5665 | NCBI | https://www.ncbi.nlm.nih.gov/genbank/ |
| Chenopodium quinoa 392 v10 | Eukaryota | Viridiplantae | genome | 44776 | Phytozome | https://phytozome-next.jgi.doe.gov/ |
| Chlamydia muridarum Nigg | Bacteria | PVC\|Chlamydiae\|Chlamydiaceae | genome | 911 | NCBI | https://www.ncbi.nlm.nih.gov/genbank/ |
| Chlamydia trachomatis 70 | Bacteria | PVC\|Chlamydiae\|Chlamydiaceae | genome | 923 | NCBI | https://www.ncbi.nlm.nih.gov/genbank/ |
| Chlamydomonas reinhardtii CC 4532 707 v61 | Eukaryota | Viridiplantae | genome | 32672 | Phytozome | https://phytozome-next.jgi.doe.gov/ |
| Chlamydophila abortus S26_3 | Bacteria | PVC\|Chlamydiae\|Chlamydiaceae | genome | 932 | NCBI | https://www.ncbi.nlm.nih.gov/genbank/ |
| Chlamydophila pneumoniae CWL029 | Bacteria | PVC\|Chlamydiae\|Chlamydiaceae | genome | 1052 | NCBI | https://www.ncbi.nlm.nih.gov/genbank/ |
| Chlorella sp. NC64A | Eukaryota | Viridiplantae | genome | 9791 | JGI | https://mycocosm.jgi.doe.gov/ |
| Chlorella vulgaris C-169 | Eukaryota | Viridiplantae | genome | 9994 | JGI | https://mycocosm.jgi.doe.gov/ |
| Chlorobaculum parvum NCIB 8327 | Bacteria | Chlorobi | genome | 2043 | NCBI | https://www.ncbi.nlm.nih.gov/genbank/ |
| Chlorobium chlorochromatii CaD3 | Bacteria | Chlorobi | genome | 2002 | NCBI | https://www.ncbi.nlm.nih.gov/genbank/ |
| Chloroflexus aurantiacus J_10_fl | Bacteria | Chloroflexi | genome | 3853 | NCBI | https://www.ncbi.nlm.nih.gov/genbank/ |
| Chlorogloeopsis fritschii PCC 6912 | Bacteria | Cyanobacteria\|Group B | genome | 6491 | NCBI | https://www.ncbi.nlm.nih.gov/genbank/ |
| Chlorogloeopsis fritschii PCC 9212 | Bacteria | Cyanobacteria\|Group B | genome | 6351 | NCBI | https://www.ncbi.nlm.nih.gov/genbank/ |
| Chloroherpeton thalassium ATCC 35110 | Bacteria | Chlorobi | genome | 2710 | NCBI | https://www.ncbi.nlm.nih.gov/genbank/ |
| Chondrocystis sp. NIES-4102 | Bacteria | Cyanobacteria\|Group C | genome | 4150 | NCBI | https://www.ncbi.nlm.nih.gov/genbank/ |
| Chromobacterium violaceum ATCC_12472 | Bacteria | Proteobacteria\|Betaproteobacteria | genome | 4407 | NCBI | https://www.ncbi.nlm.nih.gov/genbank/ |
| Chromochloris zofingiensis 461 v5.2.3.2 | Eukaryota | Viridiplantae | genome | 15369 | Phytozome | https://phytozome-next.jgi.doe.gov/ |
| Chromohalobacter salexigens DSM 3043 | Bacteria | Proteobacteria | genome | 3298 | NCBI | https://www.ncbi.nlm.nih.gov/genbank/ |
| Chroococcidiopsis sp. PCC 6712 | Bacteria | Cyanobacteria\|Group C | genome | 5031 | JGI | https://genome.jgi.doe.gov/portal/ |
| Chroococcidiopsis thermalis PCC 7203 | Bacteria | Cyanobacteria\|Group B | genome | 5961 | JGI | https://genome.jgi.doe.gov/portal/ |
| Cicer arietinum 492 v10 | Eukaryota | Viridiplantae | genome | 28269 | Phytozome | https://phytozome-next.jgi.doe.gov/ |
| Cinnamomum kanehirae 531 v3 | Eukaryota | Viridiplantae | genome | 27899 | Phytozome | https://phytozome-next.jgi.doe.gov/ |
| Citrobacter koseri ATCC_BAA895 | Bacteria | Proteobacteria | genome | 5008 | NCBI | https://www.ncbi.nlm.nih.gov/genbank/ |
| Citrus sinensis 154 | Eukaryota | Viridiplantae | genome | 46147 | Phytozome | https://phytozome-next.jgi.doe.gov/ |
| Clavibacter michiganensis NCPPB_382 | Bacteria | Actinobacteria | genome | 3079 | NCBI | https://www.ncbi.nlm.nih.gov/genbank/ |
| Closterium_sp_NIES-67 | Eukaryota | Viridiplantae | genome | 29706 | NCBI | https://www.ncbi.nlm.nih.gov/genbank/ |
| Clostridium acetobutylicum ATCC_824 | Bacteria | Firmicutes | genome | 3848 | NCBI | https://www.ncbi.nlm.nih.gov/genbank/ |
| Codium fragile | Eukaryota | Viridiplantae | transcriptome | 6868 | 1KP | https://db.cngb.org/onekp/ |
| Coleofasciculus chthonoplastes PCC 7420 | Bacteria | Cyanobacteria\|Group A | genome | 8187 | NCBI | https://www.ncbi.nlm.nih.gov/genbank/ |
| Colwellia psychrerythraea 34H | Bacteria | Proteobacteria | genome | 4910 | NCBI | https://www.ncbi.nlm.nih.gov/genbank/ |
| Coprothermobacter proteolyticus DSM 5265 | Bacteria | Firmicutes | genome | 1482 | NCBI | https://www.ncbi.nlm.nih.gov/genbank/ |
| Coraliomargarita akajimensis DSM 45221 | Bacteria | PVC\|Verrucomicrobia | genome | 3146 | GTDB | https://gtdb.ecogenomic.org/ |
| Coraliomargarita sp. CAG_312 | Bacteria | PVC\|Verrucomicrobia | MAG | 2132 | GTDB | https://gtdb.ecogenomic.org/ |
| Corylus americana var rush 835 v11 | Eukaryota | Viridiplantae | genome | 30736 | Phytozome | https://phytozome-next.jgi.doe.gov/ |
| Corymbia citriodora 507 v21 | Eukaryota | Viridiplantae | genome | 45651 | Phytozome | https://phytozome-next.jgi.doe.gov/ |
| Corynebacterium diphtheriae NCTC_13129 | Bacteria | Actinobacteria | genome | 2272 | NCBI | https://www.ncbi.nlm.nih.gov/genbank/ |
| Coxiella burnetii RSA_493 | Bacteria | Proteobacteria | genome | 1848 | NCBI | https://www.ncbi.nlm.nih.gov/genbank/ |
| Crenarchaeota sp. | Archaea | Crenarchaeota | genome | 3165 | NCBI | https://www.ncbi.nlm.nih.gov/genbank/ |
| Criblamydia sequanensis CRIB-18 | Bacteria | PVC\|Chlamydiae\|Environmental_Chlamydia | genome | 2418 | NCBI | https://www.ncbi.nlm.nih.gov/genbank/ |
| Crinalium epipsammum PCC 9333 | Bacteria | Cyanobacteria\|Group A | genome | 4935 | JGI | https://genome.jgi.doe.gov/portal/ |
| Crocosphaera chwakensis CCY0110 | Bacteria | Cyanobacteria\|Group C | genome | 6475 | NCBI | https://www.ncbi.nlm.nih.gov/genbank/ |
| Crocosphaera subtropica ATCC 51142 | Bacteria | Cyanobacteria\|Group C | genome | 5304 | NCBI | https://www.ncbi.nlm.nih.gov/genbank/ |
| Crocosphaera subtropica ATCC 51472 | Bacteria | Cyanobacteria\|Group C | genome | 5048 | NCBI | https://www.ncbi.nlm.nih.gov/genbank/ |
| Crocosphaera watsonii WH 0003 | Bacteria | Cyanobacteria\|Group C | genome | 6120 | NCBI | https://www.ncbi.nlm.nih.gov/genbank/ |
| Crocosphaera watsonii WH 8501 | Bacteria | Cyanobacteria\|Group C | genome | 5958 | NCBI | https://www.ncbi.nlm.nih.gov/genbank/ |
| Cronobacter sakazakii ATCC BAA-894 | Bacteria | Proteobacteria | genome | 4420 | NCBI | https://www.ncbi.nlm.nih.gov/genbank/ |

### Supplementary Table S1: Composition of the genomes database used for similarity searches.

| Species | Domain | Taxonomy | Assembly type | N. proteins | Assembly source | Source link |
|---|---|---|---|---|---|---|
| Crustomastix stigmata CCMP3273 | Eukaryota | Viridiplantae | transcriptome | 17228 | MMETSP | https://www.imicrobe.us/#/projects/104 |
| Cucumis sativus 122 | Eukaryota | Viridiplantae | genome | 30364 | Phytozome | https://phytozome-next.jgi.doe.gov/ |
| Cupriavidus taiwanensis | Bacteria | Proteobacteria\|Betaproteobacteria | genome | 5899 | NCBI | https://www.ncbi.nlm.nih.gov/genbank/ |
| Cyanobacterium aponinum PCC 10605 | Bacteria | Cyanobacteria\|Group C | genome | 3540 | JGI | https://genome.jgi.doe.gov/portal/ |
| cyanobacterium sp. PCC 7702 | Bacteria | Cyanobacteria\|Group B | genome | 4375 | NCBI | https://www.ncbi.nlm.nih.gov/genbank/ |
| Cyanobacterium stanieri PCC 7202 | Bacteria | Cyanobacteria\|Group C | genome | 2880 | JGI | https://genome.jgi.doe.gov/portal/ |
| Cyanobium gracile PCC 6307 | Bacteria | Cyanobacteria\|Pro/Syn | genome | 3360 | JGI | https://genome.jgi.doe.gov/portal/ |
| Cyanobium sp. NIES-981 | Bacteria | Cyanobacteria\|Pro/Syn | genome | 2871 | NCBI | https://www.ncbi.nlm.nih.gov/genbank/ |
| Cyanobium sp. NS01 | Bacteria | Cyanobacteria\|Pro/Syn | genome | 2720 | NCBI | https://www.ncbi.nlm.nih.gov/genbank/ |
| Cyanothece sp. PCC 7424 | Bacteria | Cyanobacteria\|Group C | genome | 5653 | NCBI | https://www.ncbi.nlm.nih.gov/genbank/ |
| Cyanothece sp. PCC 7425 | Bacteria | Cyanobacteria\|Group E | genome | 5232 | NCBI | https://www.ncbi.nlm.nih.gov/genbank/ |
| Cylindrospermopsis curvispora GIHE-G1 | Bacteria | Cyanobacteria\|Group B | genome | 3370 | NCBI | https://www.ncbi.nlm.nih.gov/genbank/ |
| Cylindrospermopsis raciborskii CS-505 | Bacteria | Cyanobacteria\|Group B | genome | 3406 | NCBI | https://www.ncbi.nlm.nih.gov/genbank/ |
| Cylindrospermum stagnale PCC 7417 | Bacteria | Cyanobacteria\|Group B | genome | 6595 | JGI | https://genome.jgi.doe.gov/portal/ |
| Cytophaga hutchinsonii ATCC_33406 | Bacteria | CFB | genome | 3785 | NCBI | https://www.ncbi.nlm.nih.gov/genbank/ |
| Dactylococcopsis salina PCC 8305 | Bacteria | Cyanobacteria\|Group C | genome | 3557 | JGI | https://genome.jgi.doe.gov/portal/ |
| Dechloromonas aromatica RCB | Bacteria | Proteobacteria\|Betaproteobacteria | genome | 4171 | NCBI | https://www.ncbi.nlm.nih.gov/genbank/ |
| Dehalococcoides sp. BAV1 | Bacteria | Chloroflexi | genome | 1371 | NCBI | https://www.ncbi.nlm.nih.gov/genbank/ |
| Deinococcus geothermalis DSM_11300 | Bacteria | Deinococcales | genome | 3054 | NCBI | https://www.ncbi.nlm.nih.gov/genbank/ |
| Delftia acidovorans SPH_1 | Bacteria | Proteobacteria\|Betaproteobacteria | genome | 6040 | NCBI | https://www.ncbi.nlm.nih.gov/genbank/ |
| Desulfitobacterium hafniense Y51 | Bacteria | Firmicutes | genome | 5060 | NCBI | https://www.ncbi.nlm.nih.gov/genbank/ |
| Desulfococcus oleovorans Hxd3 | Bacteria | Proteobacteria\|Deltaproteobacteria | genome | 3265 | NCBI | https://www.ncbi.nlm.nih.gov/genbank/ |
| Desulfotalea psychrophila LSv54 | Bacteria | Proteobacteria\|Deltaproteobacteria | genome | 3234 | NCBI | https://www.ncbi.nlm.nih.gov/genbank/ |
| Desulfotomaculum reducens MI_1 | Bacteria | Firmicutes | genome | 3276 | NCBI | https://www.ncbi.nlm.nih.gov/genbank/ |
| Desulfovibrio desulfuricans G20 | Bacteria | Proteobacteria\|Deltaproteobacteria | genome | 3775 | NCBI | https://www.ncbi.nlm.nih.gov/genbank/ |
| Desulfovibrio vulgaris DP4 | Bacteria | Proteobacteria\|Deltaproteobacteria | genome | 3091 | NCBI | https://www.ncbi.nlm.nih.gov/genbank/ |
| Desulfovibrio vulgaris Hildenborough | Bacteria | Proteobacteria\|Deltaproteobacteria | genome | 3531 | NCBI | https://www.ncbi.nlm.nih.gov/genbank/ |
| Desulfurococcus kamchatkensis | Archaea | Crenarchaeota | genome | 1471 | NCBI | https://www.ncbi.nlm.nih.gov/genbank/ |
| Desulfurococcus mucosus DSM2162 | Archaea | Crenarchaeota | genome | 1345 | NCBI | https://www.ncbi.nlm.nih.gov/genbank/ |
| Diaphorobacter sp. TPSY | Bacteria | Proteobacteria\|Betaproteobacteria | genome | 3479 | NCBI | https://www.ncbi.nlm.nih.gov/genbank/ |
| Dichelobacter nodosus VCS1703A | Bacteria | Proteobacteria | genome | 1280 | NCBI | https://www.ncbi.nlm.nih.gov/genbank/ |
| Dictyoglomus thermophilum H-6-12 | Bacteria | Dictyoglomi | genome | 1912 | NCBI | https://www.ncbi.nlm.nih.gov/genbank/ |
| Dinoroseobacter shibae DFL_12 | Bacteria | Proteobacteria\|Alphaproteobacteria\|Rhodobacterales | genome | 4187 | NCBI | https://www.ncbi.nlm.nih.gov/genbank/ |
| Dioscorea alata 550 v21 | Eukaryota | Viridiplantae | genome | 38603 | Phytozome | https://phytozome-next.jgi.doe.gov/ |
| Dolichomastix tenuilepis CCMP3274 | Eukaryota | Viridiplantae | transcriptome | 17000 | MMETSP | https://www.imicrobe.us/#/projects/104 |
| Dolichospermum compactum NIES-806 | Bacteria | Cyanobacteria\|Group B | genome | 4346 | NCBI | https://www.ncbi.nlm.nih.gov/genbank/ |
| Dolichospermum sp. UHCC 0315A | Bacteria | Cyanobacteria\|Group B | genome | 4563 | NCBI | https://www.ncbi.nlm.nih.gov/genbank/ |
| Dunaliella salina 325 v10 | Eukaryota | Viridiplantae | genome | 18801 | Phytozome | https://phytozome-next.jgi.doe.gov/ |
| Ehrlichia canis Jake | Bacteria | Proteobacteria\|Alphaproteobacteria\|Rickettsiales\|Anaplasmatacea | genome | 925 | NCBI | https://www.ncbi.nlm.nih.gov/genbank/ |
| Eleusine coracana 560 v11 | Eukaryota | Viridiplantae | genome | 48836 | Phytozome | https://phytozome-next.jgi.doe.gov/ |
| Elusimicrobium minutum Pei191 | Bacteria | Other bacteria | genome | 1529 | NCBI | https://www.ncbi.nlm.nih.gov/genbank/ |
| Enterobacter sp. 638 | Bacteria | Proteobacteria | genome | 4240 | NCBI | https://www.ncbi.nlm.nih.gov/genbank/ |
| Enterococcus faecalis V583 | Bacteria | Firmicutes | genome | 3265 | NCBI | https://www.ncbi.nlm.nih.gov/genbank/ |
| Erythrobacter litoralis HTCC2594 | Bacteria | Proteobacteria\|Alphaproteobacteria\|Sphingomonadales | genome | 3011 | NCBI | https://www.ncbi.nlm.nih.gov/genbank/ |
| Escherichia coli 536 | Bacteria | Proteobacteria | genome | 4620 | NCBI | https://www.ncbi.nlm.nih.gov/genbank/ |
| Eucalyptus grandis 297 v20 | Eukaryota | Viridiplantae | genome | 46280 | Phytozome | https://phytozome-next.jgi.doe.gov/ |
| Euhalothece natronophila Z-M001 | Bacteria | Cyanobacteria\|Group C | genome | 3259 | NCBI | https://www.ncbi.nlm.nih.gov/genbank/ |
| Exiguobacterium sibiricum 255-15 | Bacteria | Firmicutes | genome | 3015 | NCBI | https://www.ncbi.nlm.nih.gov/genbank/ |
| Ferroglobus placidus DSM 10642 | Archaea | Euryarchaeota | genome | 2480 | NCBI | https://www.ncbi.nlm.nih.gov/genbank/ |
| Ferroplasma acidarmanus fer1 | Archaea | Euryarchaeota | genome | 1986 | NCBI | https://www.ncbi.nlm.nih.gov/genbank/ |
| Fervidobacterium nodosum Rt17_B1 | Bacteria | Thermotogales | genome | 1750 | NCBI | https://www.ncbi.nlm.nih.gov/genbank/ |
| Finegoldia magna ATCC_29328 | Bacteria | Firmicutes | genome | 1813 | NCBI | https://www.ncbi.nlm.nih.gov/genbank/ |
| Fischerella muscicola PCC 7414 | Bacteria | Cyanobacteria\|Group B | genome | 5462 | NCBI | https://www.ncbi.nlm.nih.gov/genbank/ |
| Fischerella muscicola SAG 1427-1 | Bacteria | Cyanobacteria\|Group B | genome | 5798 | NCBI | https://www.ncbi.nlm.nih.gov/genbank/ |
| Fischerella sp. JSC-11 | Bacteria | Cyanobacteria\|Group B | genome | 4553 | NCBI | https://www.ncbi.nlm.nih.gov/genbank/ |
| Fischerella sp. NIES-3754 | Bacteria | Cyanobacteria\|Group B | genome | 4640 | NCBI | https://www.ncbi.nlm.nih.gov/genbank/ |
| Fischerella sp. NIES-4106 | Bacteria | Cyanobacteria\|Group B | genome | 5645 | NCBI | https://www.ncbi.nlm.nih.gov/genbank/ |
| Fischerella sp. PCC 9339 | Bacteria | Cyanobacteria\|Group B | genome | 6563 | NCBI | https://www.ncbi.nlm.nih.gov/genbank/ |
| Fischerella sp. PCC 9431 | Bacteria | Cyanobacteria\|Group B | genome | 6071 | JGI | https://genome.jgi.doe.gov/portal/ |
| Fischerella sp. PCC 9605 | Bacteria | Cyanobacteria\|Group B | genome | 6994 | JGI | https://genome.jgi.doe.gov/portal/ |
| Fischerella thermalis PCC 7521 | Bacteria | Cyanobacteria\|Group B | genome | 4342 | NCBI | https://www.ncbi.nlm.nih.gov/genbank/ |
| Flavobacterium johnsoniae UW101 | Bacteria | CFB | genome | 5017 | NCBI | https://www.ncbi.nlm.nih.gov/genbank/ |
| Flavobacterium psychrophilum JIP02_86 | Bacteria | CFB | genome | 2412 | NCBI | https://www.ncbi.nlm.nih.gov/genbank/ |
| Fortiea contorta PCC 7126 | Bacteria | Cyanobacteria\|Group B | genome | 5130 | JGI | https://genome.jgi.doe.gov/portal/ |
| Fragaria vesca 501 v20a2 | Eukaryota | Viridiplantae | genome | 50732 | Phytozome | https://phytozome-next.jgi.doe.gov/ |
| Francisella novicida U112 | Bacteria | Proteobacteria | genome | 1719 | NCBI | https://www.ncbi.nlm.nih.gov/genbank/ |
| Frankia alni ACN14a | Bacteria | Actinobacteria | genome | 6711 | NCBI | https://www.ncbi.nlm.nih.gov/genbank/ |
| Fremyella diplosiphon NIES-3275 | Bacteria | Cyanobacteria\|Group B | genome | 7571 | NCBI | https://www.ncbi.nlm.nih.gov/genbank/ |
| Fusobacterium nucleatum ATCC_25586 | Bacteria | Firmicutes | genome | 2067 | NCBI | https://www.ncbi.nlm.nih.gov/genbank/ |
| Geitlerinema sp. PCC 7407 | Bacteria | Cyanobacteria\|Group H | genome | 3812 | NCBI | https://www.ncbi.nlm.nih.gov/genbank/ |
| Geminocystis herdmanii PCC 6308 | Bacteria | Cyanobacteria\|Group C | genome | 4114 | JGI | https://genome.jgi.doe.gov/portal/ |
| Geminocystis sp. NIES-3708 | Bacteria | Cyanobacteria\|Group C | genome | 3429 | NCBI | https://www.ncbi.nlm.nih.gov/genbank/ |
| Geminocystis sp. NIES-3709 | Bacteria | Cyanobacteria\|Group C | genome | 3584 | NCBI | https://www.ncbi.nlm.nih.gov/genbank/ |
| Geobacillus kaustophilus HTA426 | Bacteria | Firmicutes | genome | 3540 | NCBI | https://www.ncbi.nlm.nih.gov/genbank/ |
| Geobacter metallireducens GS_15 | Bacteria | Proteobacteria\|Deltaproteobacteria | genome | 3532 | NCBI | https://www.ncbi.nlm.nih.gov/genbank/ |
| Geobacter sulfurreducens PCA | Bacteria | Proteobacteria\|Deltaproteobacteria | genome | 3445 | NCBI | https://www.ncbi.nlm.nih.gov/genbank/ |
| Geobacter uraniumreducens Rf4 | Bacteria | Proteobacteria\|Deltaproteobacteria | genome | 4357 | NCBI | https://www.ncbi.nlm.nih.gov/genbank/ |
| Gloeobacter kilaueensis JS1 | Bacteria | Cyanobacteria\|Gloeobacterales | genome | 4462 | NCBI | https://www.ncbi.nlm.nih.gov/genbank/ |
| Gloeobacter morelensis MG652769 | Bacteria | Cyanobacteria\|Gloeobacterales | genome | 4751 | GEMS | https://genome.jgi.doe.gov/portal/GEMs/GEMs.home.html |
| Gloeobacter violaceus PCC 7421 | Bacteria | Cyanobacteria\|Gloeobacterales | genome | 4430 | NCBI | https://www.ncbi.nlm.nih.gov/genbank/ |
| Gloeobacter violaceus SpSt-379 | Bacteria | Cyanobacteria\|Gloeobacterales | genome | 4157 | GEMS | https://genome.jgi.doe.gov/portal/GEMs/GEMs.home.html |
| Gloeocapsa sp. PCC 73106 | Bacteria | Cyanobacteria\|Group C | genome | 4031 | JGI | https://genome.jgi.doe.gov/portal/ |
| Gloeocapsa sp. PCC 7428 | Bacteria | Cyanobacteria\|Group B | genome | 5241 | JGI | https://genome.jgi.doe.gov/portal/ |
| Gloeomargarita lithophora | Bacteria | Cyanobacteria\|Gloeomargaritales | genome | 2990 | NCBI | https://www.ncbi.nlm.nih.gov/genbank/ |
| Gloeothece verrucosa PCC 7822 | Bacteria | Cyanobacteria\|Group C | genome | 6557 | NCBI | https://www.ncbi.nlm.nih.gov/genbank/ |
| Gluconacetobacter diazotrophicus PAI_5 | Bacteria | Proteobacteria\|Alphaproteobacteria\|Rhodospirillales | genome | 7353 | NCBI | https://www.ncbi.nlm.nih.gov/genbank/ |
| Glycine max 275 Wm82a2v1 | Eukaryota | Viridiplantae | genome | 88647 | Phytozome | https://phytozome-next.jgi.doe.gov/ |
| Gonium pectorale | Eukaryota | Viridiplantae | transcriptome | 9614 | 1KP | https://db.cngb.org/onekp/ |
| Gossypium hirsutum 527 v21 | Eukaryota | Viridiplantae | genome | 107216 | Phytozome | https://phytozome-next.jgi.doe.gov/ |
| Gramella forsetii KT0803 | Bacteria | CFB | genome | 3584 | NCBI | https://www.ncbi.nlm.nih.gov/genbank/ |
| Granulibacter bethesdensis CGDNIH1 | Bacteria | Proteobacteria\|Alphaproteobacteria\|Rhodospirillales | genome | 2437 | NCBI | https://www.ncbi.nlm.nih.gov/genbank/ |
| Haemophilus ducreyi 35000HP | Bacteria | Proteobacteria | genome | 1717 | NCBI | https://www.ncbi.nlm.nih.gov/genbank/ |
| Hahella chejuensis KCTC_2396 | Bacteria | Proteobacteria | genome | 6778 | NCBI | https://www.ncbi.nlm.nih.gov/genbank/ |
| Halalkalicoccus jeotgali B3 | Archaea | Euryarchaeota | genome | 3873 | NCBI | https://www.ncbi.nlm.nih.gov/genbank/ |
| Haliangium ochraceum DSM 14365 | Bacteria | Myxobacteria | genome | 6719 | NCBI | https://www.ncbi.nlm.nih.gov/genbank/ |
| Haloarcula hispanica ATCC 33960 | Archaea | Euryarchaeota | genome | 3859 | NCBI | https://www.ncbi.nlm.nih.gov/genbank/ |

### Supplementary Table S1: Composition of the genomes database used for similarity searches.

| Species | Domain | Taxonomy | Assembly type | N. proteins | Assembly source | Source link |
|---|---|---|---|---|---|---|
| Haloarcula marismortui ATCC_43049 | Archaea | Euryarchaeota | genome | 4240 | NCBI | https://www.ncbi.nlm.nih.gov/genbank/ |
| Halobacterium salinarum R1 | Archaea | Euryarchaeota | genome | 2749 | NCBI | https://www.ncbi.nlm.nih.gov/genbank/ |
| Haloferax volcanii DS2 | Archaea | Euryarchaeota | genome | 4015 | NCBI | https://www.ncbi.nlm.nih.gov/genbank/ |
| Halogeometricum borinquense DSM 11551 | Archaea | Euryarchaeota | genome | 4005 | NCBI | https://www.ncbi.nlm.nih.gov/genbank/ |
| Halomicrobium mukohataei DSM 12286 | Archaea | Euryarchaeota | genome | 3349 | NCBI | https://www.ncbi.nlm.nih.gov/genbank/ |
| Halomicronema hongdechloris C2206 | Bacteria | Cyanobacteria\|Group H | genome | 4810 | NCBI | https://www.ncbi.nlm.nih.gov/genbank/ |
| halophilic archaeon DL31 | Archaea | Euryarchaeota | genome | 3476 | NCBI | https://www.ncbi.nlm.nih.gov/genbank/ |
| Halopiger xanaduensis SH-6 | Archaea | Euryarchaeota | genome | 4221 | NCBI | https://www.ncbi.nlm.nih.gov/genbank/ |
| Haloquadratum walsbyi DSM_16790 | Archaea | Euryarchaeota | genome | 2646 | NCBI | https://www.ncbi.nlm.nih.gov/genbank/ |
| Halorhabdus utahensis DSM 12940 | Archaea | Euryarchaeota | genome | 2998 | NCBI | https://www.ncbi.nlm.nih.gov/genbank/ |
| Halorhodospira halophila SL1 | Bacteria | Proteobacteria | genome | 2407 | NCBI | https://www.ncbi.nlm.nih.gov/genbank/ |
| Halorubrum lacusprofundi ATCC 49239 | Archaea | Euryarchaeota | genome | 3560 | NCBI | https://www.ncbi.nlm.nih.gov/genbank/ |
| Haloterrigena turkmenica DSM 5511 | Archaea | Euryarchaeota | genome | 5113 | NCBI | https://www.ncbi.nlm.nih.gov/genbank/ |
| Halothece sp. PCC 7418 | Bacteria | Cyanobacteria\|Group C | genome | 3849 | JGI | https://genome.jgi.doe.gov/portal/ |
| Halothermothrix orenii H 168 | Bacteria | Firmicutes | genome | 2342 | NCBI | https://www.ncbi.nlm.nih.gov/genbank/ |
| Helianthus annuus 494 r12 | Eukaryota | Viridiplantae | genome | 52243 | Phytozome | https://phytozome-next.jgi.doe.gov/ |
| Helicobacter acinonychis Sheeba | Bacteria | Proteobacteria\|Deltaproteobacteria | genome | 1618 | NCBI | https://www.ncbi.nlm.nih.gov/genbank/ |
| Heliobacterium modesticaldum Ice1 | Bacteria | Firmicutes | genome | 3000 | NCBI | https://www.ncbi.nlm.nih.gov/genbank/ |
| Herminiimonas arsenicoxydans | Bacteria | Proteobacteria\|Betaproteobacteria | genome | 3295 | NCBI | https://www.ncbi.nlm.nih.gov/genbank/ |
| Herpetosiphon aurantiacus ATCC 23779 | Bacteria | Chloroflexi | genome | 5278 | NCBI | https://www.ncbi.nlm.nih.gov/genbank/ |
| Heterochlamydomonas inaequalis | Eukaryota | Viridiplantae | transcriptome | 9117 | 1KP | https://db.cngb.org/onekp/ |
| Hordeum vulgare 462 r1 | Eukaryota | Viridiplantae | genome | 248180 | Phytozome | https://phytozome-next.jgi.doe.gov/ |
| Hydrogenobaculum sp. Y04AAS1 | Bacteria | Aquificales | genome | 1629 | NCBI | https://www.ncbi.nlm.nih.gov/genbank/ |
| Hyperthermus butylicus DSM_5456 | Archaea | Crenarchaeota | genome | 1602 | NCBI | https://www.ncbi.nlm.nih.gov/genbank/ |
| Hyphomonas neptunium ATCC_15444 | Bacteria | Proteobacteria\|Alphaproteobacteria\|Hyphomonadales | genome | 3505 | NCBI | https://www.ncbi.nlm.nih.gov/genbank/ |
| Idiomarina loihiensis L2TR | Bacteria | Proteobacteria | genome | 2628 | NCBI | https://www.ncbi.nlm.nih.gov/genbank/ |
| Ignatius tetrasporus | Eukaryota | Viridiplantae | transcriptome | 7647 | 1KP | https://db.cngb.org/onekp/ |
| Ignicoccus hospitalis KIN4_I | Archaea | Crenarchaeota | genome | 1434 | NCBI | https://www.ncbi.nlm.nih.gov/genbank/ |
| Ignisphaera aggregans DSM17230 | Archaea | Crenarchaeota | genome | 1930 | NCBI | https://www.ncbi.nlm.nih.gov/genbank/ |
| Isosphaera pallida ATCC 43644 | Bacteria | PVC\|Planctomycetes | genome | 3792 | GTDB | https://gtdb.ecogenomic.org/ |
| Jannaschia sp. CCS1 | Bacteria | Proteobacteria\|Alphaproteobacteria\|Rhodobacterales | genome | 4283 | NCBI | https://www.ncbi.nlm.nih.gov/genbank/ |
| Janthinobacterium sp. Marseille | Bacteria | Proteobacteria\|Betaproteobacteria | genome | 3697 | NCBI | https://www.ncbi.nlm.nih.gov/genbank/ |
| Kamptonema formosum PCC 6407 | Bacteria | Cyanobacteria\|Group A | genome | 5642 | JGI | https://genome.jgi.doe.gov/portal/ |
| Kamptonema sp. PCC 6506 | Bacteria | Cyanobacteria\|Group A | genome | 5826 | NCBI | https://www.ncbi.nlm.nih.gov/genbank/ |
| Kineococcus radiotolerans SRS30216 | Bacteria | Actinobacteria | genome | 4681 | NCBI | https://www.ncbi.nlm.nih.gov/genbank/ |
| Kiritimatiella glycovorans str | Bacteria | PVC\|Verrucomicrobia | genome | 2429 | GTDB | https://gtdb.ecogenomic.org/ |
| Klebsiella pneumoniae MGH_78578 | Bacteria | Proteobacteria | genome | 5185 | NCBI | https://www.ncbi.nlm.nih.gov/genbank/ |
| Klebsormidium_nitens_NIES-2285 | Eukaryota | Viridiplantae | genome | 16283 | NCBI | https://www.ncbi.nlm.nih.gov/genbank/ |
| Kocuria rhizophila DC2201 | Bacteria | Actinobacteria | genome | 2357 | NCBI | https://www.ncbi.nlm.nih.gov/genbank/ |
| Korarchaeum cryptofilum OPF8 | Archaea | Korarchaeota | genome | 1602 | NCBI | https://www.ncbi.nlm.nih.gov/genbank/ |
| Lactobacillus acidophilus NCFM | Bacteria | Firmicutes | genome | 1862 | NCBI | https://www.ncbi.nlm.nih.gov/genbank/ |
| Lactococcus lactis Il1403 | Bacteria | Firmicutes | genome | 2321 | NCBI | https://www.ncbi.nlm.nih.gov/genbank/ |
| Lawsonia intracellularis PHE_MN1_00 | Bacteria | Proteobacteria\|Deltaproteobacteria | genome | 1337 | NCBI | https://www.ncbi.nlm.nih.gov/genbank/ |
| Legionella pneumophila Corby | Bacteria | Proteobacteria | genome | 3206 | NCBI | https://www.ncbi.nlm.nih.gov/genbank/ |
| Leifsonia xyli CTCB07 | Bacteria | Actinobacteria | genome | 2030 | NCBI | https://www.ncbi.nlm.nih.gov/genbank/ |
| Lens culinaris 718 v1 | Eukaryota | Viridiplantae | genome | 39141 | Phytozome | https://phytozome-next.jgi.doe.gov/ |
| Lentisphaera araneosa HTCC2155 | Bacteria | PVC\|Lentispherae | MAG | 4965 | GTDB | https://gtdb.ecogenomic.org/ |
| Leptolyngbya boryana dg5 | Bacteria | Cyanobacteria\|Group H | genome | 6176 | NCBI | https://www.ncbi.nlm.nih.gov/genbank/ |
| Leptolyngbya boryana IAM M-101 | Bacteria | Cyanobacteria\|Group H | genome | 6177 | NCBI | https://www.ncbi.nlm.nih.gov/genbank/ |
| Leptolyngbya boryana NIES-2135 | Bacteria | Cyanobacteria\|Group H | genome | 6457 | NCBI | https://www.ncbi.nlm.nih.gov/genbank/ |
| Leptolyngbya boryana PCC 6306 | Bacteria | Cyanobacteria\|Group H | genome | 6778 | JGI | https://genome.jgi.doe.gov/portal/ |
| Leptolyngbya sp. 15MV | Bacteria | Cyanobacteria\|Group H | genome | 5314 | NCBI | https://www.ncbi.nlm.nih.gov/genbank/ |
| Leptolyngbya sp. 7M | Bacteria | Cyanobacteria\|Group H | genome | 7020 | NCBI | https://www.ncbi.nlm.nih.gov/genbank/ |
| Leptolyngbya sp. BL0902 | Bacteria | Cyanobacteria\|Group H | genome | 4037 | NCBI | https://www.ncbi.nlm.nih.gov/genbank/ |
| Leptolyngbya sp. NIES-3755 | Bacteria | Cyanobacteria\|Group H | genome | 6301 | NCBI | https://www.ncbi.nlm.nih.gov/genbank/ |
| Leptolyngbya sp. O-77 | Bacteria | Cyanobacteria\|Group H | genome | 4473 | NCBI | https://www.ncbi.nlm.nih.gov/genbank/ |
| Leptolyngbya sp. PCC 6406 | Bacteria | Cyanobacteria\|Group H | genome | 5124 | JGI | https://genome.jgi.doe.gov/portal/ |
| Leptolyngbya sp. PCC 7375 | Bacteria | Cyanobacteria\|Group H | genome | 8065 | JGI | https://genome.jgi.doe.gov/portal/ |
| Leptolyngbya sp. PCC 7376 | Bacteria | Cyanobacteria\|Group C | genome | 4510 | JGI | https://genome.jgi.doe.gov/portal/ |
| Leptolyngbyaceae cyanobacterium JSC-12 | Bacteria | Cyanobacteria\|Group H | genome | 4680 | NCBI | https://www.ncbi.nlm.nih.gov/genbank/ |
| Leptosira obovata | Eukaryota | Viridiplantae | transcriptome | 7570 | 1KP | https://db.cngb.org/onekp/ |
| Leptospira borgpetersenii JB197 | Bacteria | Spirochaeta | genome | 2880 | NCBI | https://www.ncbi.nlm.nih.gov/genbank/ |
| Leptothrix cholodnii SP-6 | Bacteria | Proteobacteria\|Betaproteobacteria | genome | 4363 | NCBI | https://www.ncbi.nlm.nih.gov/genbank/ |
| Leuconostoc citreum KM20 | Bacteria | Firmicutes | genome | 1820 | NCBI | https://www.ncbi.nlm.nih.gov/genbank/ |
| Limnospira indica PCC 8005 | Bacteria | Cyanobacteria\|Group A | genome | 5653 | NCBI | https://www.ncbi.nlm.nih.gov/genbank/ |
| Listeria innocua Clip11262 | Bacteria | Firmicutes | genome | 3043 | NCBI | https://www.ncbi.nlm.nih.gov/genbank/ |
| Lobochlamys segnis | Eukaryota | Viridiplantae | transcriptome | 7468 | 1KP | https://db.cngb.org/onekp/ |
| Lobomonas rostrata | Eukaryota | Viridiplantae | transcriptome | 8230 | 1KP | https://db.cngb.org/onekp/ |
| Lotus japonicus 571 Lj10v1 | Eukaryota | Viridiplantae | genome | 28251 | Phytozome | https://phytozome-next.jgi.doe.gov/ |
| Lupinus albus 567 v1 | Eukaryota | Viridiplantae | genome | 38258 | Phytozome | https://phytozome-next.jgi.doe.gov/ |
| Lyngbya sp. PCC 8106 | Bacteria | Cyanobacteria\|Group A | genome | 6142 | NCBI | https://www.ncbi.nlm.nih.gov/genbank/ |
| Lysinibacillus sphaericus C3_41 | Bacteria | Firmicutes | genome | 4771 | NCBI | https://www.ncbi.nlm.nih.gov/genbank/ |
| Macrococcus caseolyticus JCSC5402 | Bacteria | Firmicutes | genome | 2052 | NCBI | https://www.ncbi.nlm.nih.gov/genbank/ |
| Magnetococcales bacterium DC0425bin3 | Bacteria | Proteobacteria\|Alphaproteobacteria\|Magnetococcales | genome | 3353 | GTDB | https://gtdb.ecogenomic.org/ |
| Magnetococcales bacterium DCbin2 | Bacteria | Proteobacteria\|Alphaproteobacteria\|Magnetococcales | genome | 3069 | GTDB | https://gtdb.ecogenomic.org/ |
| Magnetococcales bacterium DCbin4 | Bacteria | Proteobacteria\|Alphaproteobacteria\|Magnetococcales | genome | 3844 | GTDB | https://gtdb.ecogenomic.org/ |
| Magnetococcales bacterium GCA_002689455 | Bacteria | Proteobacteria\|Alphaproteobacteria\|MarineProteo1 | genome | 1668 | GTDB | https://gtdb.ecogenomic.org/ |
| Magnetococcales bacterium HA3dbin3 | Bacteria | Proteobacteria\|Alphaproteobacteria\|Magnetococcales | genome | 2665 | GTDB | https://gtdb.ecogenomic.org/ |
| Magnetococcales bacterium HCHbin5 | Bacteria | Proteobacteria\|Alphaproteobacteria\|Magnetococcales | genome | 3550 | GTDB | https://gtdb.ecogenomic.org/ |
| Magnetococcales bacterium sp002753095 | Bacteria | Proteobacteria\|Alphaproteobacteria\|Magnetococcales | genome | 3197 | GTDB | https://gtdb.ecogenomic.org/ |
| Magnetococcales bacterium sp002753515 | Bacteria | Proteobacteria\|Alphaproteobacteria\|Magnetococcales | genome | 3675 | GTDB | https://gtdb.ecogenomic.org/ |
| Magnetococcales bacterium sp002753565 | Bacteria | Proteobacteria\|Alphaproteobacteria\|Magnetococcales | genome | 3422 | GTDB | https://gtdb.ecogenomic.org/ |
| Magnetococcales bacterium WMHbin3 | Bacteria | Proteobacteria\|Alphaproteobacteria\|Magnetococcales | genome | 4067 | GTDB | https://gtdb.ecogenomic.org/ |
| Magnetococcales bacterium WMHbinv6 | Bacteria | Proteobacteria\|Alphaproteobacteria\|Magnetococcales | genome | 3223 | GTDB | https://gtdb.ecogenomic.org/ |
| Magnetococcus sp. MC_1 | Bacteria | Proteobacteria\|Alphaproteobacteria\|Magnetococcales | genome | 3716 | NCBI | https://www.ncbi.nlm.nih.gov/genbank/ |
| Magnetofaba australis IT-1 | Bacteria | Proteobacteria\|Alphaproteobacteria\|Magnetococcales | genome | 4295 | GTDB | https://gtdb.ecogenomic.org/ |
| Magnetospirillum magneticum AMB_1 | Bacteria | Proteobacteria\|Alphaproteobacteria\|Rhodospirillales | genome | 4559 | NCBI | https://www.ncbi.nlm.nih.gov/genbank/ |
| Manihot esculenta 671 v81 | Eukaryota | Viridiplantae | genome | 59151 | Phytozome | https://phytozome-next.jgi.doe.gov/ |
| Mannheimia succiniciproducens MBEL55E | Bacteria | Proteobacteria | genome | 2369 | NCBI | https://www.ncbi.nlm.nih.gov/genbank/ |
| Marchantia polymorpha 320 v31 | Eukaryota | Viridiplantae | genome | 24674 | Phytozome | https://phytozome-next.jgi.doe.gov/ |
| Maricaulis maris MCS10 | Bacteria | Proteobacteria\|Alphaproteobacteria\|Maricaulales | genome | 3063 | NCBI | https://www.ncbi.nlm.nih.gov/genbank/ |
| Marine group II euryarchaeote SCGC AB-629-J06 | Archaea | Euryarchaeota | genome | 550 | NCBI | https://www.ncbi.nlm.nih.gov/genbank/ |
| Marinobacter aquaeolei VT8 | Bacteria | Proteobacteria | genome | 4272 | NCBI | https://www.ncbi.nlm.nih.gov/genbank/ |
| Marinomonas sp. MWYL1 | Bacteria | Proteobacteria | genome | 4439 | NCBI | https://www.ncbi.nlm.nih.gov/genbank/ |
| Mastigocladopsis repens PCC 10914 | Bacteria | Cyanobacteria\|Group B | genome | 5761 | JGI | https://genome.jgi.doe.gov/portal/ |
| Medicago truncatula 285 Mt40v1 | Eukaryota | Viridiplantae | genome | 62319 | Phytozome | https://phytozome-next.jgi.doe.gov/ |

### Supplementary Table S1: Composition of the genomes database used for similarity searches.

| Species | Domain | Taxonomy | Assembly type | N. proteins | Assembly source | Source link |
|---|---|---|---|---|---|---|
| Mesoplasma florum L1 | Bacteria | Firmicutes | genome | 682 | NCBI | https://www.ncbi.nlm.nih.gov/genbank/ |
| Mesorhizobium sp. BNC1 | Bacteria | Proteobacteria\|Alphaproteobacteria\|Hyphomicrobiales | genome | 4543 | NCBI | https://www.ncbi.nlm.nih.gov/genbank/ |
| Metallosphaera cuprina Ar-4 | Archaea | Crenarchaeota | genome | 2029 | NCBI | https://www.ncbi.nlm.nih.gov/genbank/ |
| Metallosphaera sedula DSM_5348 | Archaea | Crenarchaeota | genome | 2256 | NCBI | https://www.ncbi.nlm.nih.gov/genbank/ |
| Methanobacterium sp. AL-21 | Archaea | Euryarchaeota | genome | 2493 | NCBI | https://www.ncbi.nlm.nih.gov/genbank/ |
| Methanobrevibacter ruminantium M1 | Archaea | Euryarchaeota | genome | 2217 | NCBI | https://www.ncbi.nlm.nih.gov/genbank/ |
| Methanobrevibacter smithii ATCC_35061 | Archaea | Crenarchaeota | genome | 1793 | NCBI | https://www.ncbi.nlm.nih.gov/genbank/ |
| Methanocaldococcus fervens AG86 | Archaea | Euryarchaeota | genome | 1581 | NCBI | https://www.ncbi.nlm.nih.gov/genbank/ |
| Methanocaldococcus infernus ME | Archaea | Euryarchaeota | genome | 1441 | NCBI | https://www.ncbi.nlm.nih.gov/genbank/ |
| Methanocaldococcus jannaschii DSM 2661 | Archaea | Euryarchaeota | genome | 1786 | NCBI | https://www.ncbi.nlm.nih.gov/genbank/ |
| Methanocaldococcus sp. FS406-22 | Archaea | Euryarchaeota | genome | 1816 | NCBI | https://www.ncbi.nlm.nih.gov/genbank/ |
| Methanocaldococcus vulcanius M7 | Archaea | Euryarchaeota | genome | 1742 | NCBI | https://www.ncbi.nlm.nih.gov/genbank/ |
| Methanocella paludicola SANAE | Archaea | Euryarchaeota | genome | 3004 | NCBI | https://www.ncbi.nlm.nih.gov/genbank/ |
| Methanococcoides burtonii DSM_6242 | Archaea | Euryarchaeota | genome | 2273 | NCBI | https://www.ncbi.nlm.nih.gov/genbank/ |
| Methanococcus aeolicus Nankai_3 | Archaea | Euryarchaeota | genome | 1490 | NCBI | https://www.ncbi.nlm.nih.gov/genbank/ |
| Methanococcus maripaludis C5 | Archaea | Euryarchaeota | genome | 1840 | NCBI | https://www.ncbi.nlm.nih.gov/genbank/ |
| Methanococcus vannielii | Archaea | Euryarchaeota | genome | 1678 | NCBI | https://www.ncbi.nlm.nih.gov/genbank/ |
| Methanococcus voltae A3 | Archaea | Euryarchaeota | genome | 1690 | NCBI | https://www.ncbi.nlm.nih.gov/genbank/ |
| Methanocorpusculum labreanum Z | Archaea | Euryarchaeota | genome | 1739 | NCBI | https://www.ncbi.nlm.nih.gov/genbank/ |
| Methanoculleus marisnigri JR1 | Archaea | Euryarchaeota | genome | 2489 | NCBI | https://www.ncbi.nlm.nih.gov/genbank/ |
| Methanohalobium evestigatum Z-7303 | Archaea | Euryarchaeota | genome | 2254 | NCBI | https://www.ncbi.nlm.nih.gov/genbank/ |
| Methanohalophilus mahii DSM 5219 | Archaea | Euryarchaeota | genome | 1987 | NCBI | https://www.ncbi.nlm.nih.gov/genbank/ |
| Methanoplanus petrolearius DSM11571 | Archaea | Euryarchaeota | genome | 2785 | NCBI | https://www.ncbi.nlm.nih.gov/genbank/ |
| Methanopyrus kandleri AV19 | Archaea | Euryarchaeota | genome | 1687 | NCBI | https://www.ncbi.nlm.nih.gov/genbank/ |
| Methanosaeta concilii GP6 | Archaea | Euryarchaeota | genome | 2850 | NCBI | https://www.ncbi.nlm.nih.gov/genbank/ |
| Methanosaeta harundinacea 6Ac | Archaea | Euryarchaeota | genome | 2462 | NCBI | https://www.ncbi.nlm.nih.gov/genbank/ |
| Methanosaeta thermophila PT | Archaea | Euryarchaeota | genome | 1696 | NCBI | https://www.ncbi.nlm.nih.gov/genbank/ |
| Methanosalsum zhilinae DSM 4017 | Archaea | Euryarchaeota | genome | 1976 | NCBI | https://www.ncbi.nlm.nih.gov/genbank/ |
| Methanosarcina acetivorans C2A | Archaea | Euryarchaeota | genome | 4540 | NCBI | https://www.ncbi.nlm.nih.gov/genbank/ |
| Methanosarcina barkeri str. Fusaro | Archaea | Euryarchaeota | genome | 3624 | NCBI | https://www.ncbi.nlm.nih.gov/genbank/ |
| Methanosarcina mazei Go1 | Archaea | Euryarchaeota | genome | 3368 | NCBI | https://www.ncbi.nlm.nih.gov/genbank/ |
| Methanosphaera stadtmanae DSM_3091 | Archaea | Euryarchaeota | genome | 1534 | NCBI | https://www.ncbi.nlm.nih.gov/genbank/ |
| Methanosphaerula palustris E1-9c | Archaea | Euryarchaeota | genome | 2655 | NCBI | https://www.ncbi.nlm.nih.gov/genbank/ |
| Methanospirillum hungatei JF_1 | Archaea | Euryarchaeota | genome | 3139 | NCBI | https://www.ncbi.nlm.nih.gov/genbank/ |
| Methanothermobacter marburgensis str. Marburg | Archaea | Euryarchaeota | genome | 1757 | NCBI | https://www.ncbi.nlm.nih.gov/genbank/ |
| Methanothermobacter thermautotrophicus | Archaea | Euryarchaeota | genome | 1907 | NCBI | https://www.ncbi.nlm.nih.gov/genbank/ |
| Methanothermococcus okinawensis IH1 | Archaea | Euryarchaeota | genome | 1729 | NCBI | https://www.ncbi.nlm.nih.gov/genbank/ |
| Methanothermus fervidus | Archaea | Euryarchaeota | genome | 1283 | NCBI | https://www.ncbi.nlm.nih.gov/genbank/ |
| Methanotorris igneus Kol 5 | Archaea | Euryarchaeota | genome | 1772 | NCBI | https://www.ncbi.nlm.nih.gov/genbank/ |
| Methylacidiphilum infernorum V4 | Bacteria | PVC\|Verrucomicrobia | genome | 2109 | GTDB | https://gtdb.ecogenomic.org/ |
| Methylibium petroleiphilum PM1 | Bacteria | Proteobacteria\|Betaproteobacteria | genome | 4449 | NCBI | https://www.ncbi.nlm.nih.gov/genbank/ |
| Methylobacillus flagellatus KT | Bacteria | Proteobacteria\|Betaproteobacteria | genome | 2753 | NCBI | https://www.ncbi.nlm.nih.gov/genbank/ |
| Methylobacterium extorquens PA1 | Bacteria | Proteobacteria\|Alphaproteobacteria\|Hyphomicrobiales | genome | 4829 | NCBI | https://www.ncbi.nlm.nih.gov/genbank/ |
| Methylocella silvestris BL2 | Bacteria | Proteobacteria\|Alphaproteobacteria\|Hyphomicrobiales | genome | 3818 | NCBI | https://www.ncbi.nlm.nih.gov/genbank/ |
| Methylococcus capsulatus Bath | Bacteria | Proteobacteria | genome | 2956 | NCBI | https://www.ncbi.nlm.nih.gov/genbank/ |
| Microcoleus sp. PCC 7113 | Bacteria | Cyanobacteria\|Group A | genome | 6658 | JGI | https://genome.jgi.doe.gov/portal/ |
| Microcoleus vaginatus FGP-2 | Bacteria | Cyanobacteria\|Group A | genome | 5102 | NCBI | https://www.ncbi.nlm.nih.gov/genbank/ |
| Microcystis aeruginosa FD4 | Bacteria | Cyanobacteria\|Group C | genome | 4708 | NCBI | https://www.ncbi.nlm.nih.gov/genbank/ |
| Microcystis aeruginosa NIES-2481 | Bacteria | Cyanobacteria\|Group C | genome | 3880 | NCBI | https://www.ncbi.nlm.nih.gov/genbank/ |
| Microcystis aeruginosa NIES-2549 | Bacteria | Cyanobacteria\|Group C | genome | 3755 | NCBI | https://www.ncbi.nlm.nih.gov/genbank/ |
| Microcystis aeruginosa NIES-298 | Bacteria | Cyanobacteria\|Group C | genome | 4344 | NCBI | https://www.ncbi.nlm.nih.gov/genbank/ |
| Microcystis aeruginosa NIES-843 | Bacteria | Cyanobacteria\|Group C | genome | 6312 | NCBI | https://www.ncbi.nlm.nih.gov/genbank/ |
| Microcystis aeruginosa PCC 7806 | Bacteria | Cyanobacteria\|Group C | genome | 4891 | NCBI | https://www.ncbi.nlm.nih.gov/genbank/ |
| Microcystis panniformis FACHB-1757 | Bacteria | Cyanobacteria\|Group C | genome | 4462 | NCBI | https://www.ncbi.nlm.nih.gov/genbank/ |
| Microcystis sp. MC19 | Bacteria | Cyanobacteria\|Group C | genome | 4302 | NCBI | https://www.ncbi.nlm.nih.gov/genbank/ |
| Microcystis viridis NIES-102 | Bacteria | Cyanobacteria\|Group C | genome | 4655 | NCBI | https://www.ncbi.nlm.nih.gov/genbank/ |
| Micromonas pusilla CCMP1545 228 v30 | Eukaryota | Viridiplantae | genome | 10660 | Phytozome | https://phytozome-next.jgi.doe.gov/ |
| Micromonas sp. RCC299 | Eukaryota | Viridiplantae | genome | 10109 | JGI | https://mycocosm.jgi.doe.gov/ |
| Midichloria bacterium endos.Acanthamoeba | Bacteria | Proteobacteria\|Alphaproteobacteria\|Rickettsiales\|Midichloriaceae | genome | 1549 | GTDB | https://gtdb.ecogenomic.org/ |
| Mimulus guttatus 256 v20 | Eukaryota | Viridiplantae | genome | 33573 | Phytozome | https://phytozome-next.jgi.doe.gov/ |
| Miscanthus sinensis 497 v71 | Eukaryota | Viridiplantae | genome | 89486 | Phytozome | https://phytozome-next.jgi.doe.gov/ |
| Moorella thermoacetica ATCC 39073 | Bacteria | Firmicutes | genome | 2463 | NCBI | https://www.ncbi.nlm.nih.gov/genbank/ |
| Moorena producens 3L | Bacteria | Cyanobacteria\|Group A | genome | 7355 | NCBI | https://www.ncbi.nlm.nih.gov/genbank/ |
| Moorena producens JHB | Bacteria | Cyanobacteria\|Group A | genome | 7590 | NCBI | https://www.ncbi.nlm.nih.gov/genbank/ |
| Moorena producens PAL-8-15-08-1 | Bacteria | Cyanobacteria\|Group A | genome | 7469 | NCBI | https://www.ncbi.nlm.nih.gov/genbank/ |
| Musa acuminata 304 v1 | Eukaryota | Viridiplantae | genome | 36528 | Phytozome | https://phytozome-next.jgi.doe.gov/ |
| Mycobacterium abscessus | Bacteria | Actinobacteria | genome | 4941 | NCBI | https://www.ncbi.nlm.nih.gov/genbank/ |
| Mycoplasma agalactiae PG2 | Bacteria | Firmicutes | genome | 742 | NCBI | https://www.ncbi.nlm.nih.gov/genbank/ |
| Myxococcus xanthus DK_1622 | Bacteria | Myxobacteria | genome | 7331 | NCBI | https://www.ncbi.nlm.nih.gov/genbank/ |
| Myxosarcina sp. GI1 | Bacteria | Cyanobacteria\|Group C | genome | 6110 | NCBI | https://www.ncbi.nlm.nih.gov/genbank/ |
| Nanoarchaeum equitans Kin4_M | Archaea | Euryarchaeota | genome | 536 | NCBI | https://www.ncbi.nlm.nih.gov/genbank/ |
| Natranaerobius thermophilus JWNM-WN-LF | Bacteria | Firmicutes | genome | 2906 | NCBI | https://www.ncbi.nlm.nih.gov/genbank/ |
| Natrialba magadii ATCC 43099 | Archaea | Euryarchaeota | genome | 4212 | NCBI | https://www.ncbi.nlm.nih.gov/genbank/ |
| Natronomonas pharaonis DSM_2160 | Archaea | Euryarchaeota | genome | 2822 | NCBI | https://www.ncbi.nlm.nih.gov/genbank/ |
| Nautilia profundicola AmH | Bacteria | Proteobacteria\|Deltaproteobacteria | genome | 1730 | NCBI | https://www.ncbi.nlm.nih.gov/genbank/ |
| Neisseria gonorrhoeae FA_1090 | Bacteria | Proteobacteria\|Betaproteobacteria | genome | 2002 | NCBI | https://www.ncbi.nlm.nih.gov/genbank/ |
| Neorickettsia sennetsu Miyayama | Bacteria | Proteobacteria\|Alphaproteobacteria\|Rickettsiales\|Anaplasmatacea | genome | 932 | NCBI | https://www.ncbi.nlm.nih.gov/genbank/ |
| Nephroselmis olivacea | Eukaryota | Viridiplantae | transcriptome | 10728 | 1KP | https://db.cngb.org/onekp/ |
| Nitratiruptor sp. SB155_2 | Bacteria | Proteobacteria\|Deltaproteobacteria | genome | 1843 | NCBI | https://www.ncbi.nlm.nih.gov/genbank/ |
| Nitrobacter hamburgensis X14 | Bacteria | Proteobacteria\|Alphaproteobacteria\|Hyphomonadales | genome | 4326 | NCBI | https://www.ncbi.nlm.nih.gov/genbank/ |
| Nitrosococcus oceani ATCC_19707 | Bacteria | Proteobacteria | genome | 3017 | NCBI | https://www.ncbi.nlm.nih.gov/genbank/ |
| Nitrosomonas europaea ATCC_19718 | Bacteria | Proteobacteria\|Betaproteobacteria | genome | 2461 | NCBI | https://www.ncbi.nlm.nih.gov/genbank/ |
| Nitrosopumilus maritimus SCM1 | Archaea | Thaumarchaeota | genome | 1795 | NCBI | https://www.ncbi.nlm.nih.gov/genbank/ |
| Nitrosospira multiformis ATCC_25196 | Bacteria | Proteobacteria\|Betaproteobacteria | genome | 2805 | NCBI | https://www.ncbi.nlm.nih.gov/genbank/ |
| Nocardia farcinica IFM_10152 | Bacteria | Actinobacteria | genome | 5936 | NCBI | https://www.ncbi.nlm.nih.gov/genbank/ |
| Nocardioides sp. JS614 | Bacteria | Actinobacteria | genome | 4909 | NCBI | https://www.ncbi.nlm.nih.gov/genbank/ |
| Nodosilinea nodulosa PCC 7104 | Bacteria | Cyanobacteria\|Group H | genome | 6278 | JGI | https://genome.jgi.doe.gov/portal/ |
| Nodularia spumigena CCY9414 | Bacteria | Cyanobacteria\|Group B | genome | 4860 | NCBI | https://www.ncbi.nlm.nih.gov/genbank/ |
| Nodularia spumigena UHCC 0039 | Bacteria | Cyanobacteria\|Group B | genome | 4403 | NCBI | https://www.ncbi.nlm.nih.gov/genbank/ |
| Nostoc carneum NIES-2107 | Bacteria | Cyanobacteria\|Group B | genome | 7034 | NCBI | https://www.ncbi.nlm.nih.gov/genbank/ |
| Nostoc edaphicum CCNP1411 | Bacteria | Cyanobacteria\|Group B | genome | 6466 | NCBI | https://www.ncbi.nlm.nih.gov/genbank/ |
| Nostoc flagelliforme CCNUN1 | Bacteria | Cyanobacteria\|Group B | genome | 8045 | NCBI | https://www.ncbi.nlm.nih.gov/genbank/ |
| Nostoc linckia NIES-25 | Bacteria | Cyanobacteria\|Group B | genome | 6616 | NCBI | https://www.ncbi.nlm.nih.gov/genbank/ |
| Nostoc piscinale CENA21 | Bacteria | Cyanobacteria\|Group B | genome | 5310 | NCBI | https://www.ncbi.nlm.nih.gov/genbank/ |
| Nostoc punctiforme 11 | Bacteria | Cyanobacteria\|Group B | genome | 6690 | NCBI | https://www.ncbi.nlm.nih.gov/genbank/ |
| Nostoc punctiforme PCC 73102 | Bacteria | Cyanobacteria\|Group B | genome | 6982 | NCBI | https://www.ncbi.nlm.nih.gov/genbank/ |

### Supplementary Table S1: Composition of the genomes database used for similarity searches.

| Species | Domain | Taxonomy | Assembly type | N. proteins | Assembly source | Source link |
|---|---|---|---|---|---|---|
| Nostoc sp. ATCC 53789 | Bacteria | Cyanobacteria\|Group B | genome | 6724 | NCBI | https://www.ncbi.nlm.nih.gov/genbank/ |
| Nostoc sp. C052 | Bacteria | Cyanobacteria\|Group B | genome | 7769 | NCBI | https://www.ncbi.nlm.nih.gov/genbank/ |
| Nostoc sp. C057 | Bacteria | Cyanobacteria\|Group B | genome | 7343 | NCBI | https://www.ncbi.nlm.nih.gov/genbank/ |
| Nostoc sp. CENA543 | Bacteria | Cyanobacteria\|Group B | genome | 5823 | NCBI | https://www.ncbi.nlm.nih.gov/genbank/ |
| Nostoc sp. Lobaria pulmonaria cyanobiont | Bacteria | Cyanobacteria\|Group B | genome | 6074 | NCBI | https://www.ncbi.nlm.nih.gov/genbank/ |
| Nostoc sp. NIES-2111 | Bacteria | Cyanobacteria\|Group B | genome | 6329 | NCBI | https://www.ncbi.nlm.nih.gov/genbank/ |
| Nostoc sp. NIES-3756 | Bacteria | Cyanobacteria\|Group B | genome | 5771 | NCBI | https://www.ncbi.nlm.nih.gov/genbank/ |
| Nostoc sp. NIES-4103 | Bacteria | Cyanobacteria\|Group B | genome | 6562 | NCBI | https://www.ncbi.nlm.nih.gov/genbank/ |
| Nostoc sp. PCC 7107 | Bacteria | Cyanobacteria\|Group B | genome | 5372 | JGI | https://genome.jgi.doe.gov/portal/ |
| Nostoc sp. PCC 7524 | Bacteria | Cyanobacteria\|Group B | genome | 5534 | JGI | https://genome.jgi.doe.gov/portal/ |
| Nostoc sp. PCC7120 | Bacteria | Cyanobacteria\|Group B | genome | 6130 | NCBI | https://www.ncbi.nlm.nih.gov/genbank/ |
| Nostoc sp. Peltigera membranacea cyanobiont | Bacteria | Cyanobacteria\|Group B | genome | 7042 | NCBI | https://www.ncbi.nlm.nih.gov/genbank/ |
| Nostoc sp. TCL240-02 | Bacteria | Cyanobacteria\|Group B | genome | 6094 | NCBI | https://www.ncbi.nlm.nih.gov/genbank/ |
| Nostoc sp. TCL26-01 | Bacteria | Cyanobacteria\|Group B | genome | 6025 | NCBI | https://www.ncbi.nlm.nih.gov/genbank/ |
| Nostoc sp. UHCC 0702 | Bacteria | Cyanobacteria\|Group B | genome | 7090 | NCBI | https://www.ncbi.nlm.nih.gov/genbank/ |
| Nostoc sphaeroides | Bacteria | Cyanobacteria\|Group B | genome | 5504 | NCBI | https://www.ncbi.nlm.nih.gov/genbank/ |
| Nostoc sphaeroides CCNUC1 | Bacteria | Cyanobacteria\|Group B | genome | 7630 | NCBI | https://www.ncbi.nlm.nih.gov/genbank/ |
| Nostocales cyanobacterium HT-58-2 | Bacteria | Cyanobacteria\|Group B | genome | 6131 | NCBI | https://www.ncbi.nlm.nih.gov/genbank/ |
| Novosphingobium aromaticivorans DSM_12444 | Bacteria | Proteobacteria\|Alphaproteobacteria\|Sphingomonadales | genome | 3937 | NCBI | https://www.ncbi.nlm.nih.gov/genbank/ |
| Nymphaea colorata 566 v12 | Eukaryota | Viridiplantae | genome | 43975 | Phytozome | https://phytozome-next.jgi.doe.gov/ |
| Oceanobacillus iheyensis HTE831 | Bacteria | Firmicutes | genome | 3500 | NCBI | https://www.ncbi.nlm.nih.gov/genbank/ |
| Ochrobactrum anthropi ATCC_49188 | Bacteria | Proteobacteria\|Alphaproteobacteria\|Hyphomicrobiales | genome | 4799 | NCBI | https://www.ncbi.nlm.nih.gov/genbank/ |
| Oedogonium cardiacum | Eukaryota | Viridiplantae | transcriptome | 7246 | 1KP | https://db.cngb.org/onekp/ |
| Oenococcus oeni PSU_1 | Bacteria | Firmicutes | genome | 1691 | NCBI | https://www.ncbi.nlm.nih.gov/genbank/ |
| Olea europaea 451 v10 | Eukaryota | Viridiplantae | genome | 50684 | Phytozome | https://phytozome-next.jgi.doe.gov/ |
| Oligotropha carboxidovorans OM5 | Bacteria | Proteobacteria\|Alphaproteobacteria\|Hyphomicrobiales | genome | 3847 | NCBI | https://www.ncbi.nlm.nih.gov/genbank/ |
| Onion yellows Phytoplasma sp. OY-M | Bacteria | Firmicutes | genome | 754 | NCBI | https://www.ncbi.nlm.nih.gov/genbank/ |
| Opitutaceae bacterium TAV5 | Bacteria | PVC\|Verrucomicrobia | genome | 6010 | GTDB | https://gtdb.ecogenomic.org/ |
| Opitutus terrae PB90-1 | Bacteria | PVC\|Verrucomicrobia | genome | 4612 | NCBI | https://www.ncbi.nlm.nih.gov/genbank/ |
| Oropetium thomaeum 386 v10 | Eukaryota | Viridiplantae | genome | 28437 | Phytozome | https://phytozome-next.jgi.doe.gov/ |
| Oryza sativa 323 v70 | Eukaryota | Viridiplantae | genome | 52424 | Phytozome | https://phytozome-next.jgi.doe.gov/ |
| Oscillatoria acuminata PCC 6304 | Bacteria | Cyanobacteria\|Group A | genome | 5953 | JGI | https://genome.jgi.doe.gov/portal/ |
| Oscillatoria nigro-viridis PCC 7112 | Bacteria | Cyanobacteria\|Group A | genome | 6805 | JGI | https://genome.jgi.doe.gov/portal/ |
| Oscillatoria sp. PCC 10802 | Bacteria | Cyanobacteria\|Group A | genome | 6950 | JGI | https://genome.jgi.doe.gov/portal/ |
| Ostreococcus lucimarinus 231 | Eukaryota | Viridiplantae | genome | 7796 | Phytozome | https://phytozome-next.jgi.doe.gov/ |
| Ostreococcus tauri | Eukaryota | Viridiplantae | genome | 7725 | JGI | https://mycocosm.jgi.doe.gov/ |
| Oxynema sp. AP17 | Bacteria | Cyanobacteria\|Group A | genome | 4832 | NCBI | https://www.ncbi.nlm.nih.gov/genbank/ |
| Panicum virgatum 516 v51 | Eukaryota | Viridiplantae | genome | 129942 | Phytozome | https://phytozome-next.jgi.doe.gov/ |
| Parabacteroides distasonis ATCC_8503 | Bacteria | CFB | genome | 3850 | NCBI | https://www.ncbi.nlm.nih.gov/genbank/ |
| Parachlamydia acanthamoebae | Bacteria | PVC\|Chlamydiae\|Environmental_Chlamydia | genome | 2809 | NCBI | https://www.ncbi.nlm.nih.gov/genbank/ |
| Parachlorella kessleri | Eukaryota | Viridiplantae | transcriptome | 7510 | 1KP | https://db.cngb.org/onekp/ |
| Paracoccus denitrificans PD1222 | Bacteria | Proteobacteria\|Alphaproteobacteria\|Rhodobacterales | genome | 5077 | NCBI | https://www.ncbi.nlm.nih.gov/genbank/ |
| Parvibaculum lavamentivorans DS1 | Bacteria | Proteobacteria\|Alphaproteobacteria\|Hyphomicrobiales | genome | 3636 | NCBI | https://www.ncbi.nlm.nih.gov/genbank/ |
| Pasteurella multocida Pm70 | Bacteria | Proteobacteria | genome | 2015 | NCBI | https://www.ncbi.nlm.nih.gov/genbank/ |
| Pectobacterium atrosepticum SCRI1043 | Bacteria | Proteobacteria | genome | 4472 | NCBI | https://www.ncbi.nlm.nih.gov/genbank/ |
| Pediococcus pentosaceus ATCC_25745 | Bacteria | Firmicutes | genome | 1755 | NCBI | https://www.ncbi.nlm.nih.gov/genbank/ |
| Pelagibacter ubique HTCC1062 | Bacteria | Proteobacteria\|Alphaproteobacteria\|Pelagibacterales | genome | 1354 | NCBI | https://www.ncbi.nlm.nih.gov/genbank/ |
| Pelobacter carbinolicus DSM_2380 | Bacteria | Proteobacteria\|Deltaproteobacteria | genome | 3352 | NCBI | https://www.ncbi.nlm.nih.gov/genbank/ |
| Pelobacter propionicus DSM_2379 | Bacteria | Proteobacteria\|Deltaproteobacteria | genome | 3804 | NCBI | https://www.ncbi.nlm.nih.gov/genbank/ |
| Pelodictyon luteolum DSM_273 | Bacteria | Chlorobi | genome | 2083 | NCBI | https://www.ncbi.nlm.nih.gov/genbank/ |
| Pelotomaculum thermopropionicum SI | Bacteria | Firmicutes | genome | 2918 | NCBI | https://www.ncbi.nlm.nih.gov/genbank/ |
| Petrotoga mobilis SJ95 | Bacteria | Thermotogales | genome | 1898 | NCBI | https://www.ncbi.nlm.nih.gov/genbank/ |
| Phacotus lenticularis | Eukaryota | Viridiplantae | transcriptome | 8348 | 1KP | https://db.cngb.org/onekp/ |
| Phaseolus acutifolius 580 v10 | Eukaryota | Viridiplantae | genome | 50635 | Phytozome | https://phytozome-next.jgi.doe.gov/ |
| Phenylobacterium zucineum HLK1 | Bacteria | Proteobacteria\|Alphaproteobacteria\|Caulobacterales | genome | 3854 | NCBI | https://www.ncbi.nlm.nih.gov/genbank/ |
| Phormidium sp. ETS-05 | Bacteria | Cyanobacteria\|Group A | genome | 5058 | NCBI | https://www.ncbi.nlm.nih.gov/genbank/ |
| Photobacterium profundum SS9 | Bacteria | Proteobacteria | genome | 5489 | NCBI | https://www.ncbi.nlm.nih.gov/genbank/ |
| Photorhabdus luminescens TTO1 | Bacteria | Proteobacteria | genome | 4683 | NCBI | https://www.ncbi.nlm.nih.gov/genbank/ |
| Phycisphaera mikurensis NBRC 102666 | Bacteria | PVC\|Planctomycetes | genome | 3127 | GTDB | https://gtdb.ecogenomic.org/ |
| Physcomitrium patens 318 v33 | Eukaryota | Viridiplantae | genome | 87533 | Phytozome | https://phytozome-next.jgi.doe.gov/ |
| Phytoplasma australiense | Bacteria | Firmicutes | genome | 696 | NCBI | https://www.ncbi.nlm.nih.gov/genbank/ |
| Picrophilus torridus DSM_9790 | Archaea | Euryarchaeota | genome | 1535 | NCBI | https://www.ncbi.nlm.nih.gov/genbank/ |
| Pirellula staleyi DSM 6068 | Bacteria | PVC\|Planctomycetes | genome | 4719 | GTDB | https://gtdb.ecogenomic.org/ |
| Pirula salina | Eukaryota | Viridiplantae | transcriptome | 5817 | 1KP | https://db.cngb.org/onekp/ |
| Planctopirus limnophila DSM 3776 | Bacteria | PVC\|Planctomycetes | genome | 4288 | GTDB | https://gtdb.ecogenomic.org/ |
| Planktothrix rubescens NIVA-CYA 18 | Bacteria | Cyanobacteria\|Group A | genome | 4823 | NCBI | https://www.ncbi.nlm.nih.gov/genbank/ |
| Planophila laetevirens | Eukaryota | Viridiplantae | transcriptome | 6158 | 1KP | https://db.cngb.org/onekp/ |
| Plesiocystis pacifica SIR-1 | Bacteria | Myxobacteria | genome | 8450 | NCBI | https://www.ncbi.nlm.nih.gov/genbank/ |
| Pleurocapsa sp. PCC 7319 | Bacteria | Cyanobacteria\|Group C | genome | 6566 | JGI | https://genome.jgi.doe.gov/portal/ |
| Pleurocapsa sp. PCC 7327 | Bacteria | Cyanobacteria\|Group C | genome | 4573 | JGI | https://genome.jgi.doe.gov/portal/ |
| Polaromonas sp. JS666 | Bacteria | Proteobacteria\|Betaproteobacteria | genome | 5453 | NCBI | https://www.ncbi.nlm.nih.gov/genbank/ |
| Polynucleobacter sp. QLW P1DMWA 1 | Bacteria | Proteobacteria\|Betaproteobacteria | genome | 2077 | NCBI | https://www.ncbi.nlm.nih.gov/genbank/ |
| Populus trichocarpa | Eukaryota | Viridiplantae | genome | 45555 | JGI | https://mycocosm.jgi.doe.gov/ |
| Populus trichocarpa 533 v41 | Eukaryota | Viridiplantae | genome | 52400 | Phytozome | https://phytozome-next.jgi.doe.gov/ |
| Porphyromonas gingivalis W83 | Bacteria | CFB | genome | 1909 | NCBI | https://www.ncbi.nlm.nih.gov/genbank/ |
| Portulaca amilis 692 v10 | Eukaryota | Viridiplantae | genome | 58732 | Phytozome | https://phytozome-next.jgi.doe.gov/ |
| Prasinococcus capsulatus CCMP1194 | Eukaryota | Viridiplantae | transcriptome | 8610 | MMETSP | https://www.imicrobe.us/#/projects/104 |
| Prasinoderma coloniale CCMP1413 | Eukaryota | Viridiplantae | transcriptome | 10355 | MMETSP | https://www.imicrobe.us/#/projects/104 |
| Prochlorococcus marinus str. AS9601 | Bacteria | Cyanobacteria\|Pro/Syn | genome | 1921 | NCBI | https://www.ncbi.nlm.nih.gov/genbank/ |
| Prochlorococcus marinus str. CCMP1375 | Bacteria | Cyanobacteria\|Pro/Syn | genome | 1883 | NCBI | https://www.ncbi.nlm.nih.gov/genbank/ |
| Prochlorococcus marinus str. MIT 9215 | Bacteria | Cyanobacteria\|Pro/Syn | genome | 1983 | NCBI | https://www.ncbi.nlm.nih.gov/genbank/ |
| Prochlorococcus marinus str. MIT 9301 | Bacteria | Cyanobacteria\|Pro/Syn | genome | 1907 | NCBI | https://www.ncbi.nlm.nih.gov/genbank/ |
| Prochlorococcus marinus str. MIT 9303 | Bacteria | Cyanobacteria\|Pro/Syn | genome | 2997 | NCBI | https://www.ncbi.nlm.nih.gov/genbank/ |
| Prochlorococcus marinus str. MIT 9312 | Bacteria | Cyanobacteria\|Pro/Syn | genome | 1810 | NCBI | https://www.ncbi.nlm.nih.gov/genbank/ |
| Prochlorococcus marinus str. MIT 9313 | Bacteria | Cyanobacteria\|Pro/Syn | genome | 2269 | NCBI | https://www.ncbi.nlm.nih.gov/genbank/ |
| Prochlorococcus marinus str. MIT 9515 | Bacteria | Cyanobacteria\|Pro/Syn | genome | 1906 | NCBI | https://www.ncbi.nlm.nih.gov/genbank/ |
| Prochlorococcus marinus str. NATL1A | Bacteria | Cyanobacteria\|Pro/Syn | genome | 2193 | NCBI | https://www.ncbi.nlm.nih.gov/genbank/ |
| Prochlorococcus marinus str. NATL2A | Bacteria | Cyanobacteria\|Pro/Syn | genome | 2163 | NCBI | https://www.ncbi.nlm.nih.gov/genbank/ |
| Prochlorococcus marinus subsp. pastoris str. CCMP1986 | Bacteria | Cyanobacteria\|Pro/Syn | genome | 1717 | NCBI | https://www.ncbi.nlm.nih.gov/genbank/ |
| Propionibacterium acnes KPA171202 | Bacteria | Actinobacteria | genome | 2297 | NCBI | https://www.ncbi.nlm.nih.gov/genbank/ |
| Prosthecochloris vibrioformis DSM_265 | Bacteria | Chlorobi | genome | 1753 | NCBI | https://www.ncbi.nlm.nih.gov/genbank/ |
| Proteobacteria bacterium sp002937495 | Bacteria | Proteobacteria\|Alphaproteobacteria\|MarineProteo1 | genome | 1171 | GTDB | https://gtdb.ecogenomic.org/ |
| Proteobacteria bacterium UBA2136 | Bacteria | Proteobacteria\|Alphaproteobacteria\|MarineProteo1 | genome | 1732 | GTDB | https://gtdb.ecogenomic.org/ |
| Proteobacteria bacterium UBA6156 | Bacteria | Proteobacteria\|Alphaproteobacteria\|MarineProteo1 | genome | 1320 | GTDB | https://gtdb.ecogenomic.org/ |
| Proteobacteria bacterium UBA6503 | Bacteria | Proteobacteria\|Alphaproteobacteria\|MarineProteo1 | genome | 1763 | GTDB | https://gtdb.ecogenomic.org/ |
| Proteus mirabilis HI4320 | Bacteria | Proteobacteria | genome | 3662 | NCBI | https://www.ncbi.nlm.nih.gov/genbank/ |

### Supplementary Table S1: Composition of the genomes database used for similarity searches.

| Species | Domain | Taxonomy | Assembly type | N. proteins | Assembly source | Source link |
|---|---|---|---|---|---|---|
| Protochlamydia amoebophila UWE25 | Bacteria | PVC\|Chlamydiae\|Environmental_Chlamydia | genome | 2031 | NCBI | https://www.ncbi.nlm.nih.gov/genbank/ |
| Prunus persica 298 v21 | Eukaryota | Viridiplantae | genome | 47089 | Phytozome | https://phytozome-next.jgi.doe.gov/ |
| Pseudanabaena biceps PCC 7429 | Bacteria | Cyanobacteria\|Pseudanabaenales | genome | 4727 | JGI | https://genome.jgi.doe.gov/portal/ |
| Pseudanabaena sp. ABRG5-3 | Bacteria | Cyanobacteria\|Pseudanabaenales | genome | 4924 | NCBI | https://www.ncbi.nlm.nih.gov/genbank/ |
| Pseudanabaena sp. PCC 6802 | Bacteria | Cyanobacteria\|Pseudanabaenales | genome | 5212 | JGI | https://genome.jgi.doe.gov/portal/ |
| Pseudanabaena sp. PCC 7367 | Bacteria | Cyanobacteria\|Pseudanabaenales | genome | 3908 | JGI | https://genome.jgi.doe.gov/portal/ |
| Pseudoalteromonas atlantica T6c | Bacteria | Proteobacteria | genome | 4281 | NCBI | https://www.ncbi.nlm.nih.gov/genbank/ |
| Pseudomonas aeruginosa PAO1 | Bacteria | Proteobacteria | genome | 5566 | NCBI | https://www.ncbi.nlm.nih.gov/genbank/ |
| Pseudomonas fluorescens sp003241595 | Bacteria | Proteobacteria\|Alphaproteobacteria\|MarineProteo1 | genome | 1481 | GTDB | https://gtdb.ecogenomic.org/ |
| Pseudoneochloris marina | Eukaryota | Viridiplantae | transcriptome | 6231 | 1KP | https://db.cngb.org/onekp/ |
| Psychrobacter arcticus 273_4 | Bacteria | Proteobacteria | genome | 2120 | NCBI | https://www.ncbi.nlm.nih.gov/genbank/ |
| Psychromonas ingrahamii 37 | Bacteria | Proteobacteria | genome | 3545 | NCBI | https://www.ncbi.nlm.nih.gov/genbank/ |
| Pyramimonas parkeae CCMP726 | Eukaryota | Viridiplantae | transcriptome | 23234 | MMETSP | https://www.imicrobe.us/#/projects/104 |
| Pyrobaculum aerophilum IM2 | Archaea | Crenarchaeota | genome | 2605 | NCBI | https://www.ncbi.nlm.nih.gov/genbank/ |
| Pyrobaculum arsenaticum DSM 13514 | Archaea | Crenarchaeota | genome | 2299 | NCBI | https://www.ncbi.nlm.nih.gov/genbank/ |
| Pyrobaculum calidifontis JCM 11548 | Archaea | Crenarchaeota | genome | 2149 | NCBI | https://www.ncbi.nlm.nih.gov/genbank/ |
| Pyrobaculum islandicum DSM 4184 | Archaea | Crenarchaeota | genome | 1978 | NCBI | https://www.ncbi.nlm.nih.gov/genbank/ |
| Pyrobaculum sp. 1860 | Archaea | Crenarchaeota | genome | 2827 | NCBI | https://www.ncbi.nlm.nih.gov/genbank/ |
| Pyrococcus abyssi GE5 | Archaea | Euryarchaeota | genome | 1782 | NCBI | https://www.ncbi.nlm.nih.gov/genbank/ |
| Pyrococcus furiosus DSM 3638 | Archaea | Euryarchaeota | genome | 2125 | NCBI | https://www.ncbi.nlm.nih.gov/genbank/ |
| Pyrococcus horikoshii OT3 | Archaea | Euryarchaeota | genome | 1955 | NCBI | https://www.ncbi.nlm.nih.gov/genbank/ |
| Pyrococcus sp. NA2 | Archaea | Euryarchaeota | genome | 1980 | NCBI | https://www.ncbi.nlm.nih.gov/genbank/ |
| Pyrococcus yayanosii CH1 | Archaea | Euryarchaeota | genome | 1865 | NCBI | https://www.ncbi.nlm.nih.gov/genbank/ |
| Pyrolobus fumarii 1A | Archaea | Crenarchaeota | genome | 1967 | NCBI | https://www.ncbi.nlm.nih.gov/genbank/ |
| Quercus rubra 687 v21 | Eukaryota | Viridiplantae | genome | 47780 | Phytozome | https://phytozome-next.jgi.doe.gov/ |
| Ralstonia eutropha H16 | Bacteria | Proteobacteria\|Betaproteobacteria | genome | 6626 | NCBI | https://www.ncbi.nlm.nih.gov/genbank/ |
| Raphidiopsis brookii D9 | Bacteria | Cyanobacteria\|Group B | genome | 3007 | NCBI | https://www.ncbi.nlm.nih.gov/genbank/ |
| Raphidiopsis curvata NIES-932 | Bacteria | Cyanobacteria\|Group B | genome | 2924 | NCBI | https://www.ncbi.nlm.nih.gov/genbank/ |
| Renibacterium salmoninarum ATCC_33209 | Bacteria | Actinobacteria | genome | 3507 | NCBI | https://www.ncbi.nlm.nih.gov/genbank/ |
| Rhizobium etli CFN_42 | Bacteria | Proteobacteria\|Alphaproteobacteria\|Hyphomicrobiales | genome | 5963 | NCBI | https://www.ncbi.nlm.nih.gov/genbank/ |
| Rhodobacter sphaeroides 2_4_1 | Bacteria | Proteobacteria\|Alphaproteobacteria\|Rhodobacterales | genome | 4242 | NCBI | https://www.ncbi.nlm.nih.gov/genbank/ |
| Rhodococcus sp. RHA1 | Bacteria | Actinobacteria | genome | 9145 | NCBI | https://www.ncbi.nlm.nih.gov/genbank/ |
| Rhodoferax ferrireducens T118 | Bacteria | Proteobacteria\|Betaproteobacteria | genome | 4418 | NCBI | https://www.ncbi.nlm.nih.gov/genbank/ |
| Rhodopirellula baltica SH_1 | Bacteria | PVC\|Planctomycetes | genome | 7325 | NCBI | https://www.ncbi.nlm.nih.gov/genbank/ |
| Rhodopseudomonas palustris BisA53 | Bacteria | Proteobacteria\|Alphaproteobacteria\|Hyphomicrobiales | genome | 4878 | NCBI | https://www.ncbi.nlm.nih.gov/genbank/ |
| Rhodospirillum rubrum ATCC 11170 | Bacteria | Proteobacteria\|Alphaproteobacteria\|Rhodospirillales | genome | 3841 | NCBI | https://www.ncbi.nlm.nih.gov/genbank/ |
| Richelia intracellularis HH01 | Bacteria | Cyanobacteria\|Group B | genome | 3841 | NCBI | https://www.ncbi.nlm.nih.gov/genbank/ |
| Ricinus communis 119 | Eukaryota | Viridiplantae | genome | 31221 | Phytozome | https://phytozome-next.jgi.doe.gov/ |
| Rickettsia akari Hartford | Bacteria | Proteobacteria\|Alphaproteobacteria\|Rickettsiales\|Rickettsiaceae | genome | 1259 | NCBI | https://www.ncbi.nlm.nih.gov/genbank/ |
| Rickettsia rhipicephali str.3-7-female6-CWPP | Bacteria | Proteobacteria\|Alphaproteobacteria\|Rickettsiales\|Rickettsiaceae | genome | 1511 | GTDB | https://gtdb.ecogenomic.org/ |
| Rickettsia rickettsii str | Bacteria | Proteobacteria\|Alphaproteobacteria\|Rickettsiales\|Rickettsiaceae | genome | 1433 | GTDB | https://gtdb.ecogenomic.org/ |
| Rickettsia typhi str.Wilmington | Bacteria | Proteobacteria\|Alphaproteobacteria\|Rickettsiales\|Rickettsiaceae | genome | 868 | GTDB | https://gtdb.ecogenomic.org/ |
| Rickettsiales bacterium Ac37b | Bacteria | Proteobacteria\|Alphaproteobacteria\|Rickettsiales\|Arcanobacteriac | genome | 1854 | GTDB | https://gtdb.ecogenomic.org/ |
| Rickettsiales bacterium sp002725445 | Bacteria | Proteobacteria\|Alphaproteobacteria\|Rickettsiales | genome | 1767 | GTDB | https://gtdb.ecogenomic.org/ |
| Rickettsiales bacterium sp005787525 | Bacteria | Proteobacteria\|Alphaproteobacteria\|Rickettsiales\|Anaplasmatacea | genome | 1377 | GTDB | https://gtdb.ecogenomic.org/ |
| Rickettsiales bacterium UBA2645 | Bacteria | Proteobacteria\|Alphaproteobacteria\|Rickettsiales\|Anaplasmatacea | genome | 1747 | GTDB | https://gtdb.ecogenomic.org/ |
| Rippkaea orientalis PCC 8801 | Bacteria | Cyanobacteria\|Group C | genome | 4349 | NCBI | https://www.ncbi.nlm.nih.gov/genbank/ |
| Rippkaea orientalis PCC 8802 | Bacteria | Cyanobacteria\|Group C | genome | 4403 | NCBI | https://www.ncbi.nlm.nih.gov/genbank/ |
| Rivularia sp. PCC 7116 | Bacteria | Cyanobacteria\|Group B | genome | 6848 | JGI | https://genome.jgi.doe.gov/portal/ |
| Roseiflexus castenholzii DSM_13941 | Bacteria | Chloroflexi | genome | 4330 | NCBI | https://www.ncbi.nlm.nih.gov/genbank/ |
| Roseobacter denitrificans OCh_114 | Bacteria | Proteobacteria\|Alphaproteobacteria\|Rhodobacterales | genome | 4129 | NCBI | https://www.ncbi.nlm.nih.gov/genbank/ |
| Rubidibacter lacunae KORDI 51-2 | Bacteria | Cyanobacteria\|Group C | genome | 3450 | NCBI | https://www.ncbi.nlm.nih.gov/genbank/ |
| Rubrobacter xylanophilus DSM_9941 | Bacteria | Actinobacteria | genome | 3140 | NCBI | https://www.ncbi.nlm.nih.gov/genbank/ |
| Ruegeria sp. TM1040 | Bacteria | Proteobacteria\|Alphaproteobacteria\|Rhodobacterales | genome | 3864 | NCBI | https://www.ncbi.nlm.nih.gov/genbank/ |
| Saccharophagus degradans 2_40 | Bacteria | Proteobacteria | genome | 4007 | NCBI | https://www.ncbi.nlm.nih.gov/genbank/ |
| Saccharopolyspora erythraea NRRL_2338 | Bacteria | Actinobacteria | genome | 7197 | NCBI | https://www.ncbi.nlm.nih.gov/genbank/ |
| Salinibacter ruber DSM_13855 | Bacteria | CFB | genome | 2833 | NCBI | https://www.ncbi.nlm.nih.gov/genbank/ |
| Salinispora arenicola CNS_205 | Bacteria | Actinobacteria | genome | 4917 | NCBI | https://www.ncbi.nlm.nih.gov/genbank/ |
| Salix purpurea 519 v51 | Eukaryota | Viridiplantae | genome | 57462 | Phytozome | https://phytozome-next.jgi.doe.gov/ |
| Salmonella enterica arizonae | Bacteria | Proteobacteria | genome | 4498 | NCBI | https://www.ncbi.nlm.nih.gov/genbank/ |
| Schrenkiella parvula 574 v22 | Eukaryota | Viridiplantae | genome | 26847 | Phytozome | https://phytozome-next.jgi.doe.gov/ |
| Scytonema hofmanni PCC 7110 | Bacteria | Cyanobacteria\|Group B | genome | 9591 | NCBI | https://www.ncbi.nlm.nih.gov/genbank/ |
| Scytonema sp. HK-05 | Bacteria | Cyanobacteria\|Group B | genome | 7394 | NCBI | https://www.ncbi.nlm.nih.gov/genbank/ |
| Scytonema sp. NIES-4073 | Bacteria | Cyanobacteria\|Group B | genome | 7400 | NCBI | https://www.ncbi.nlm.nih.gov/genbank/ |
| Selaginella moellendorffii 91 | Eukaryota | Viridiplantae | genome | 22285 | Phytozome | https://phytozome-next.jgi.doe.gov/ |
| Serratia proteamaculans 568 | Bacteria | Proteobacteria | genome | 4942 | NCBI | https://www.ncbi.nlm.nih.gov/genbank/ |
| Setaria viridis 500 v21 | Eukaryota | Viridiplantae | genome | 52459 | Phytozome | https://phytozome-next.jgi.doe.gov/ |
| Shewanella amazonensis SB2B | Bacteria | Proteobacteria | genome | 3645 | NCBI | https://www.ncbi.nlm.nih.gov/genbank/ |
| Shigella boydii Sb227 | Bacteria | Proteobacteria | genome | 4282 | NCBI | https://www.ncbi.nlm.nih.gov/genbank/ |
| Silicibacter pomeroyi DSS_3 | Bacteria | Proteobacteria\|Alphaproteobacteria\|Rhodobacterales | genome | 4252 | NCBI | https://www.ncbi.nlm.nih.gov/genbank/ |
| Simkania negevensis Z | Bacteria | PVC\|Chlamydiae\|Simkaniaceae | genome | 2512 | NCBI | https://www.ncbi.nlm.nih.gov/genbank/ |
| Singulisphaera acidiphila DSM 18658 | Bacteria | PVC\|Planctomycetes | genome | 7588 | GTDB | https://gtdb.ecogenomic.org/ |
| Sinorhizobium medicae WSM419 | Bacteria | Proteobacteria\|Alphaproteobacteria\|Hyphomicrobiales | genome | 6213 | NCBI | https://www.ncbi.nlm.nih.gov/genbank/ |
| Sodalis glossinidius | Bacteria | Proteobacteria | genome | 2715 | NCBI | https://www.ncbi.nlm.nih.gov/genbank/ |
| Solanum lycopersicum 390 ITAG24 | Eukaryota | Viridiplantae | genome | 34725 | Phytozome | https://phytozome-next.jgi.doe.gov/ |
| Solanum tuberosum 686 v61 | Eukaryota | Viridiplantae | genome | 44851 | Phytozome | https://phytozome-next.jgi.doe.gov/ |
| Solibacter usitatus Ellin6076 | Bacteria | Acidobacteria | genome | 7826 | NCBI | https://www.ncbi.nlm.nih.gov/genbank/ |
| Sorangium cellulosum | Bacteria | Myxobacteria | genome | 9381 | NCBI | https://www.ncbi.nlm.nih.gov/genbank/ |
| Sorghum bicolor 454 v311 | Eukaryota | Viridiplantae | genome | 47121 | Phytozome | https://phytozome-next.jgi.doe.gov/ |
| Spermatozopsis exsultans | Eukaryota | Viridiplantae | transcriptome | 6524 | 1KP | https://db.cngb.org/onekp/ |
| Sphaerospermopsis kisseleviana NIES-73 | Bacteria | Cyanobacteria\|Group B | genome | 4730 | NCBI | https://www.ncbi.nlm.nih.gov/genbank/ |
| Sphingomonas wittichii RW1 | Bacteria | Proteobacteria\|Alphaproteobacteria\|Sphingomonadales | genome | 5345 | NCBI | https://www.ncbi.nlm.nih.gov/genbank/ |
| Sphingopyxis alaskensis RB2256 | Bacteria | Proteobacteria\|Alphaproteobacteria\|Sphingomonadales | genome | 3195 | NCBI | https://www.ncbi.nlm.nih.gov/genbank/ |
| Spinacia oleracea 575 Spov3 | Eukaryota | Viridiplantae | genome | 34875 | Phytozome | https://phytozome-next.jgi.doe.gov/ |
| Spirodela polyrhiza 290 v2 | Eukaryota | Viridiplantae | genome | 19623 | Phytozome | https://phytozome-next.jgi.doe.gov/ |
| Spirulina major PCC 6313 | Bacteria | Cyanobacteria\|Group C | genome | 4356 | JGI | https://genome.jgi.doe.gov/portal/ |
| Spirulina subsalsa PCC 9445 | Bacteria | Cyanobacteria\|Group C | genome | 4565 | JGI | https://genome.jgi.doe.gov/portal/ |
| Stanieria cyanosphaera PCC 7437 | Bacteria | Cyanobacteria\|Group C | genome | 4955 | JGI | https://genome.jgi.doe.gov/portal/ |
| Stanieria sp. NIES-3757 | Bacteria | Cyanobacteria\|Group C | genome | 4603 | NCBI | https://www.ncbi.nlm.nih.gov/genbank/ |
| Staphylococcus aureus MRSA252 | Bacteria | Firmicutes | genome | 2656 | NCBI | https://www.ncbi.nlm.nih.gov/genbank/ |
| Staphylothermus hellenicus DSM 12710 | Archaea | Crenarchaeota | genome | 1672 | JGI | https://genome.jgi.doe.gov/portal/ |
| Staphylothermus marinus F1 | Archaea | Crenarchaeota | genome | 1570 | NCBI | https://www.ncbi.nlm.nih.gov/genbank/ |
| Stenotrophomonas maltophilia K279a | Bacteria | Proteobacteria | genome | 4386 | NCBI | https://www.ncbi.nlm.nih.gov/genbank/ |
| Streptococcus agalactiae 2603V_R | Bacteria | Actinobacteria | genome | 2124 | NCBI | https://www.ncbi.nlm.nih.gov/genbank/ |
| Streptomyces avermitilis MA_4680 | Bacteria | Actinobacteria | genome | 7676 | NCBI | https://www.ncbi.nlm.nih.gov/genbank/ |
| Sulcia muelleri GWSS | Bacteria | Chlorobi | genome | 227 | NCBI | https://www.ncbi.nlm.nih.gov/genbank/ |

### Supplementary Table S1: Composition of the genomes database used for similarity searches.

| Species | Domain | Taxonomy | Assembly type | N. proteins | Assembly source | Source link |
|---|---|---|---|---|---|---|
| Sulfolobus acidocaldarius DSM_639 | Archaea | Crenarchaeota | genome | 2223 | NCBI | https://www.ncbi.nlm.nih.gov/genbank/ |
| Sulfolobus islandicus L.S.2.15 | Archaea | Crenarchaeota | genome | 2737 | NCBI | https://www.ncbi.nlm.nih.gov/genbank/ |
| Sulfolobus solfataricus P2 | Archaea | Crenarchaeota | genome | 2977 | NCBI | https://www.ncbi.nlm.nih.gov/genbank/ |
| Sulfolobus tokodaii str 7 | Archaea | Crenarchaeota | genome | 2825 | NCBI | https://www.ncbi.nlm.nih.gov/genbank/ |
| Sulfurihydrogenibium sp. YO3AOP1 | Bacteria | Aquificales | genome | 1721 | NCBI | https://www.ncbi.nlm.nih.gov/genbank/ |
| Sulfurimonas denitrificans DSM 1251 | Bacteria | Proteobacteria\|Deltaproteobacteria | genome | 2096 | NCBI | https://www.ncbi.nlm.nih.gov/genbank/ |
| Sulfurovum sp. NBC37_1 | Bacteria | Proteobacteria\|Deltaproteobacteria | genome | 2438 | NCBI | https://www.ncbi.nlm.nih.gov/genbank/ |
| Symbiobacterium thermophilum IAM_14863 | Bacteria | Firmicutes | genome | 3338 | NCBI | https://www.ncbi.nlm.nih.gov/genbank/ |
| Synechococcus elongatus PCC 11801 | Bacteria | Cyanobacteria\|Group H | genome | 2478 | NCBI | https://www.ncbi.nlm.nih.gov/genbank/ |
| Synechococcus elongatus PCC 11802 | Bacteria | Cyanobacteria\|Group H | genome | 2476 | NCBI | https://www.ncbi.nlm.nih.gov/genbank/ |
| Synechococcus elongatus PCC 6301 | Bacteria | Cyanobacteria\|Group H | genome | 2527 | NCBI | https://www.ncbi.nlm.nih.gov/genbank/ |
| Synechococcus elongatus PCC 7942 | Bacteria | Cyanobacteria\|Group H | genome | 2728 | NCBI | https://www.ncbi.nlm.nih.gov/genbank/ |
| Synechococcus elongatus UTEX 3055 | Bacteria | Cyanobacteria\|Group H | genome | 2851 | NCBI | https://www.ncbi.nlm.nih.gov/genbank/ |
| Synechococcus sp. BDU 130192 | Bacteria | Cyanobacteria\|Group C | genome | 3014 | NCBI | https://www.ncbi.nlm.nih.gov/genbank/ |
| Synechococcus sp. CC9311 | Bacteria | Cyanobacteria\|Pro/Syn | genome | 2892 | NCBI | https://www.ncbi.nlm.nih.gov/genbank/ |
| Synechococcus sp. CC9605 | Bacteria | Cyanobacteria\|Pro/Syn | genome | 2645 | NCBI | https://www.ncbi.nlm.nih.gov/genbank/ |
| Synechococcus sp. CC9902 | Bacteria | Cyanobacteria\|Pro/Syn | genome | 2307 | NCBI | https://www.ncbi.nlm.nih.gov/genbank/ |
| Synechococcus sp. JA-2-3Ba | Bacteria | Cyanobacteria\|Thermostichales | genome | 2862 | NCBI | https://www.ncbi.nlm.nih.gov/genbank/ |
| Synechococcus sp. JA-3-3Ab | Bacteria | Cyanobacteria\|Thermostichales | genome | 2760 | NCBI | https://www.ncbi.nlm.nih.gov/genbank/ |
| Synechococcus sp. M44_DOE_062 | Bacteria | Cyanobacteria\|Thermostichales | genome | 2575 | NCBI | https://www.ncbi.nlm.nih.gov/genbank/ |
| Synechococcus sp. NIES-970 | Bacteria | Cyanobacteria\|Group C | genome | 2844 | NCBI | https://www.ncbi.nlm.nih.gov/genbank/ |
| Synechococcus sp. NKBG042902 | Bacteria | Cyanobacteria\|Group C | genome | 3066 | NCBI | https://www.ncbi.nlm.nih.gov/genbank/ |
| Synechococcus sp. NKBG15041c | Bacteria | Cyanobacteria\|Group C | genome | 2865 | NCBI | https://www.ncbi.nlm.nih.gov/genbank/ |
| Synechococcus sp. PCC 11901 | Bacteria | Cyanobacteria\|Group C | genome | 3162 | NCBI | https://www.ncbi.nlm.nih.gov/genbank/ |
| Synechococcus sp. PCC 6312 | Bacteria | Cyanobacteria\|Group E | genome | 3740 | JGI | https://genome.jgi.doe.gov/portal/ |
| Synechococcus sp. PCC 7002 | Bacteria | Cyanobacteria\|Group C | genome | 3252 | NCBI | https://www.ncbi.nlm.nih.gov/genbank/ |
| Synechococcus sp. PCC 7003 | Bacteria | Cyanobacteria\|Group C | genome | 3085 | NCBI | https://www.ncbi.nlm.nih.gov/genbank/ |
| Synechococcus sp. PCC 7117 | Bacteria | Cyanobacteria\|Group C | genome | 3152 | NCBI | https://www.ncbi.nlm.nih.gov/genbank/ |
| Synechococcus sp. PCC 73109 | Bacteria | Cyanobacteria\|Group C | genome | 3012 | NCBI | https://www.ncbi.nlm.nih.gov/genbank/ |
| Synechococcus sp. PCC 7335 | Bacteria | Cyanobacteria\|Group H | genome | 5469 | NCBI | https://www.ncbi.nlm.nih.gov/genbank/ |
| Synechococcus sp. PCC 7336 | Bacteria | Cyanobacteria\|Thermostichales | genome | 4682 | JGI | https://genome.jgi.doe.gov/portal/ |
| Synechococcus sp. PCC 7502 | Bacteria | Cyanobacteria\|Pseudanabaenales | genome | 3574 | JGI | https://genome.jgi.doe.gov/portal/ |
| Synechococcus sp. PCC 8807 | Bacteria | Cyanobacteria\|Group C | genome | 3072 | NCBI | https://www.ncbi.nlm.nih.gov/genbank/ |
| Synechococcus sp. RCC307 | Bacteria | Cyanobacteria\|Pro/Syn | genome | 2535 | NCBI | https://www.ncbi.nlm.nih.gov/genbank/ |
| Synechococcus sp. UTEX 2973 | Bacteria | Cyanobacteria\|Group H | genome | 2693 | NCBI | https://www.ncbi.nlm.nih.gov/genbank/ |
| Synechococcus sp. WH 5701 | Bacteria | Cyanobacteria\|Pro/Syn | genome | 3346 | NCBI | https://www.ncbi.nlm.nih.gov/genbank/ |
| Synechococcus sp. WH 7803 | Bacteria | Cyanobacteria\|Pro/Syn | genome | 2533 | NCBI | https://www.ncbi.nlm.nih.gov/genbank/ |
| Synechococcus sp. WH 8102 | Bacteria | Cyanobacteria\|Pro/Syn | genome | 2519 | NCBI | https://www.ncbi.nlm.nih.gov/genbank/ |
| Synechocystis sp. CACIAM 05 | Bacteria | Cyanobacteria\|Group C | genome | 3189 | NCBI | https://www.ncbi.nlm.nih.gov/genbank/ |
| Synechocystis sp. IPPAS B-1465 | Bacteria | Cyanobacteria\|Group C | genome | 3526 | NCBI | https://www.ncbi.nlm.nih.gov/genbank/ |
| Synechocystis sp. PCC 6714 | Bacteria | Cyanobacteria\|Group C | genome | 3310 | NCBI | https://www.ncbi.nlm.nih.gov/genbank/ |
| Synechocystis sp. PCC 6803 | Bacteria | Cyanobacteria\|Group C | genome | 3572 | NCBI | https://www.ncbi.nlm.nih.gov/genbank/ |
| Synechocystis sp. PCC 6803 substr. GT-I | Bacteria | Cyanobacteria\|Group C | genome | 3181 | NCBI | https://www.ncbi.nlm.nih.gov/genbank/ |
| Synechocystis sp. PCC 6803 substr. PCC-N | Bacteria | Cyanobacteria\|Group C | genome | 3180 | NCBI | https://www.ncbi.nlm.nih.gov/genbank/ |
| Synechocystis sp. PCC 6803 substr. PCC-P | Bacteria | Cyanobacteria\|Group C | genome | 3179 | NCBI | https://www.ncbi.nlm.nih.gov/genbank/ |
| Synechocystis sp. PCC 7338 | Bacteria | Cyanobacteria\|Group C | genome | 3373 | NCBI | https://www.ncbi.nlm.nih.gov/genbank/ |
| Synechocystis sp. PCC 7509 | Bacteria | Cyanobacteria\|Group B | genome | 4787 | JGI | https://genome.jgi.doe.gov/portal/ |
| Syntrophobacter fumaroxidans MPOB | Bacteria | Proteobacteria\|Deltaproteobacteria | genome | 4064 | NCBI | https://www.ncbi.nlm.nih.gov/genbank/ |
| Syntrophomonas wolfei subsp. Wolfei str. Goettingen | Bacteria | Firmicutes | genome | 2504 | NCBI | https://www.ncbi.nlm.nih.gov/genbank/ |
| Syntrophus aciditrophicus SB | Bacteria | Proteobacteria\|Deltaproteobacteria | genome | 3168 | NCBI | https://www.ncbi.nlm.nih.gov/genbank/ |
| Tetraselmis chuii PLY429 | Eukaryota | Viridiplantae | transcriptome | 23036 | MMETSP | https://www.imicrobe.us/#/projects/104 |
| Theobroma cacao 233 | Eukaryota | Viridiplantae | genome | 44404 | Phytozome | https://phytozome-next.jgi.doe.gov/ |
| Thermoanaerobacter pseudethanolicus ATCC 33223 | Bacteria | Firmicutes | genome | 2243 | NCBI | https://www.ncbi.nlm.nih.gov/genbank/ |
| Thermobifida fusca YX | Bacteria | Actinobacteria | genome | 3110 | NCBI | https://www.ncbi.nlm.nih.gov/genbank/ |
| Thermococcus barophilus MP | Archaea | Euryarchaeota | genome | 2207 | NCBI | https://www.ncbi.nlm.nih.gov/genbank/ |
| Thermococcus gammatolerans EJ3 | Archaea | Euryarchaeota | genome | 2156 | NCBI | https://www.ncbi.nlm.nih.gov/genbank/ |
| Thermococcus kodakarensis KOD1 | Archaea | Euryarchaeota | genome | 2306 | NCBI | https://www.ncbi.nlm.nih.gov/genbank/ |
| Thermococcus onnurineus NA1 | Archaea | Euryarchaeota | genome | 1976 | NCBI | https://www.ncbi.nlm.nih.gov/genbank/ |
| Thermococcus sibiricus MM 739 | Archaea | Euryarchaeota | genome | 2035 | NCBI | https://www.ncbi.nlm.nih.gov/genbank/ |
| Thermococcus sp. AM4 | Archaea | Euryarchaeota | genome | 2168 | NCBI | https://www.ncbi.nlm.nih.gov/genbank/ |
| Thermodesulfovibrio yellowstonii DSM 11347 | Bacteria | Nitrospirae | genome | 2033 | NCBI | https://www.ncbi.nlm.nih.gov/genbank/ |
| Thermofilum pendens Hrk_5 | Archaea | Crenarchaeota | genome | 1876 | NCBI | https://www.ncbi.nlm.nih.gov/genbank/ |
| Thermoleptolyngbya sp. PKUAC-SCTA183 | Bacteria | Cyanobacteria\|Group H | genome | 4273 | NCBI | https://www.ncbi.nlm.nih.gov/genbank/ |
| Thermoleptolyngbya sp. PKUAC-SCTB121 | Bacteria | Cyanobacteria\|Group H | genome | 4296 | NCBI | https://www.ncbi.nlm.nih.gov/genbank/ |
| Thermomicrobium roseum DSM 5159 | Bacteria | Chloroflexi | genome | 2854 | NCBI | https://www.ncbi.nlm.nih.gov/genbank/ |
| Thermoplasma acidophilum DSM_1728 | Archaea | Euryarchaeota | genome | 1482 | NCBI | https://www.ncbi.nlm.nih.gov/genbank/ |
| Thermoplasma volcanium GSS1 | Archaea | Euryarchaeota | genome | 1499 | NCBI | https://www.ncbi.nlm.nih.gov/genbank/ |
| Thermoproteus neutrophilus V24Sta | Archaea | Crenarchaeota | genome | 1966 | NCBI | https://www.ncbi.nlm.nih.gov/genbank/ |
| Thermoproteus tenax Kra 1 | Archaea | Crenarchaeota | genome | 2049 | NCBI | https://www.ncbi.nlm.nih.gov/genbank/ |
| Thermoproteus uzoniensis 768-20 | Archaea | Crenarchaeota | genome | 2186 | NCBI | https://www.ncbi.nlm.nih.gov/genbank/ |
| Thermosipho melanesiensis BI429 | Bacteria | Thermotogales | genome | 1879 | NCBI | https://www.ncbi.nlm.nih.gov/genbank/ |
| Thermosphaera aggregans DSM 11486 | Archaea | Crenarchaeota | genome | 1387 | NCBI | https://www.ncbi.nlm.nih.gov/genbank/ |
| Thermosynechococcus elongatus BP-1 | Bacteria | Cyanobacteria\|Group E | genome | 2476 | NCBI | https://www.ncbi.nlm.nih.gov/genbank/ |
| Thermosynechococcus elongatus PKUAC-SCTE542 | Bacteria | Cyanobacteria\|Group E | genome | 2505 | NCBI | https://www.ncbi.nlm.nih.gov/genbank/ |
| Thermosynechococcus sp. CL-1 | Bacteria | Cyanobacteria\|Group E | genome | 2512 | NCBI | https://www.ncbi.nlm.nih.gov/genbank/ |
| Thermosynechococcus sp. NK55a | Bacteria | Cyanobacteria\|Group E | genome | 2330 | NCBI | https://www.ncbi.nlm.nih.gov/genbank/ |
| Thermosynechococcus sp. TA-1 | Bacteria | Cyanobacteria\|Group E | genome | 2523 | NCBI | https://www.ncbi.nlm.nih.gov/genbank/ |
| Thermosynechococcus vulcanus NIES-2134 | Bacteria | Cyanobacteria\|Group E | genome | 2413 | NCBI | https://www.ncbi.nlm.nih.gov/genbank/ |
| Thermotoga lettingae TMO | Bacteria | Thermotogales | genome | 2040 | NCBI | https://www.ncbi.nlm.nih.gov/genbank/ |
| Thermus thermophilus HB8 | Bacteria | Deinococcales | genome | 2238 | NCBI | https://www.ncbi.nlm.nih.gov/genbank/ |
| Thioalkalivibrio sp. HL-EbGR7 | Bacteria | Proteobacteria | genome | 3283 | NCBI | https://www.ncbi.nlm.nih.gov/genbank/ |
| Thiobacillus denitrificans ATCC_25259 | Bacteria | Proteobacteria\|Betaproteobacteria | genome | 2827 | NCBI | https://www.ncbi.nlm.nih.gov/genbank/ |
| Thiomicrospira crunogena XCL_2 | Bacteria | Proteobacteria | genome | 2196 | NCBI | https://www.ncbi.nlm.nih.gov/genbank/ |
| Thuja plicata 572 v31 | Eukaryota | Viridiplantae | genome | 65809 | Phytozome | https://phytozome-next.jgi.doe.gov/ |
| Tolypothrix sp. PCC 7910 | Bacteria | Cyanobacteria\|Group B | genome | 6654 | NCBI | https://www.ncbi.nlm.nih.gov/genbank/ |
| Tolypothrix sp. PCC 9009 | Bacteria | Cyanobacteria\|Group B | genome | 7225 | JGI | https://genome.jgi.doe.gov/portal/ |
| Tolypothrix tenuis PCC 7101 | Bacteria | Cyanobacteria\|Group B | genome | 7079 | NCBI | https://www.ncbi.nlm.nih.gov/genbank/ |
| Trebouxia arboricola | Eukaryota | Viridiplantae | transcriptome | 7032 | 1KP | https://db.cngb.org/onekp/ |
| Trentepohlia annulata | Eukaryota | Viridiplantae | transcriptome | 4705 | 1KP | https://db.cngb.org/onekp/ |
| Treponema denticola ATCC_35405 | Bacteria | Spirochaeta | genome | 2767 | NCBI | https://www.ncbi.nlm.nih.gov/genbank/ |
| Trichodesmium erythraeum | Bacteria | Cyanobacteria\|Group A | genome | 4451 | NCBI | https://www.ncbi.nlm.nih.gov/genbank/ |
| Trichormus variabilis | Bacteria | Cyanobacteria\|Group B | genome | 5661 | NCBI | https://www.ncbi.nlm.nih.gov/genbank/ |
| Trichormus variabilis ATCC 29413 | Bacteria | Cyanobacteria\|Group B | genome | 5687 | NCBI | https://www.ncbi.nlm.nih.gov/genbank/ |
| Trifolium pratense 385 v2 | Eukaryota | Viridiplantae | genome | 41297 | Phytozome | https://phytozome-next.jgi.doe.gov/ |
| Triticum aestivum 296 v22 | Eukaryota | Viridiplantae | genome | 293053 | Phytozome | https://phytozome-next.jgi.doe.gov/ |
| Tropheryma whipplei TW08_27 | Bacteria | Actinobacteria | genome | 783 | NCBI | https://www.ncbi.nlm.nih.gov/genbank/ |

### Supplementary Table S1: Composition of the genomes database used for similarity searches.

| Species | Domain | Taxonomy | Assembly type | N. proteins | Assembly source | Source link |
|---|---|---|---|---|---|---|
| Uncultured Anoxichlamydiales bacterium B4_bin.14 | Bacteria | PVC\|Chlamydiae\|Anoxichlamydiales | MAG | 1825 | GTDB | https://gtdb.ecogenomic.org/ |
| Uncultured Anoxichlamydiales bacterium K1060_chlam_5 | Bacteria | PVC\|Chlamydiae\|Anoxichlamydiales | MAG | 1346 | GTDB | https://gtdb.ecogenomic.org/ |
| Uncultured Anoxichlamydiales bacterium REEB52 | Bacteria | PVC\|Chlamydiae\|Anoxichlamydiales | MAG | 1666 | GTDB | https://gtdb.ecogenomic.org/ |
| Uncultured Anoxichlamydiales bacterium SZAS-17 | Bacteria | PVC\|Chlamydiae\|Anoxichlamydiales | MAG | 2208 | GTDB | https://gtdb.ecogenomic.org/ |
| Uncultured Anoxichlamydiales bacterium Zod_Metabat.1285 | Bacteria | PVC\|Chlamydiae\|Anoxichlamydiales | MAG | 1113 | GTDB | https://gtdb.ecogenomic.org/ |
| Uncultured Chlamydiales bacterium AG-470-G05 | Bacteria | PVC\|Chlamydiae\|Basal_Chlamydia | MAG | 1031 | GTDB | https://gtdb.ecogenomic.org/ |
| Uncultured Chlamydiales bacterium K_DeepCast_35m_m1_012 | Bacteria | PVC\|Chlamydiae\|Basal_Chlamydia | MAG | 1575 | GTDB | https://gtdb.ecogenomic.org/ |
| Uncultured Chlamydiales bacterium K_DeepCast_35m_m2_096 | Bacteria | PVC\|Chlamydiae\|Basal_Chlamydia | MAG | 1308 | GTDB | https://gtdb.ecogenomic.org/ |
| Uncultured Chlamydiales bacterium K_DeepCast_35m_m2_204 | Bacteria | PVC\|Chlamydiae\|Basal_Chlamydia | MAG | 1286 | GTDB | https://gtdb.ecogenomic.org/ |
| Uncultured Chlamydiales bacterium K940_chlam_8 | Bacteria | PVC\|Chlamydiae\|Basal_Chlamydia | MAG | 1319 | GTDB | https://gtdb.ecogenomic.org/ |
| Uncultured Gloeobacterales MAG ES-bin-141 | Bacteria | Cyanobacteria\|Gloeobacterales | genome | 3886 | GEMS | https://genome.jgi.doe.gov/portal/GEMs/GEMs.home.html |
| Uncultured Gloeobacterales MAG ES-bin-313 | Bacteria | Cyanobacteria\|Gloeobacterales | genome | 4133 | GEMS | https://genome.jgi.doe.gov/portal/GEMs/GEMs.home.html |
| Uncultured Marine group II euryarchaeote | Archaea | Euryarchaeota | genome | 1781 | NCBI | https://www.ncbi.nlm.nih.gov/genbank/ |
| Uncultured Methanogenic archaeon RC-I | Archaea | Euryarchaeota | genome | 3085 | NCBI | https://www.ncbi.nlm.nih.gov/genbank/ |
| Uncultured Rhabdochlamydia bacterium 15C | Bacteria | PVC\|Chlamydiae\|Rhabdochlamydia | MAG | 1380 | GTDB | https://gtdb.ecogenomic.org/ |
| Uncultured Rhabdochlamydia bacterium Ga0074140 | Bacteria | PVC\|Chlamydiae\|Rhabdochlamydia | MAG | 1648 | GTDB | https://gtdb.ecogenomic.org/ |
| Uncultured Rhabdochlamydia bacterium palsa_1444 | Bacteria | PVC\|Chlamydiae\|Rhabdochlamydia | MAG | 2102 | GTDB | https://gtdb.ecogenomic.org/ |
| Uncultured Rhabdochlamydia bacterium SZAS-12 | Bacteria | PVC\|Chlamydiae\|Rhabdochlamydia | MAG | 1925 | GTDB | https://gtdb.ecogenomic.org/ |
| Uncultured Rhabdochlamydia bacterium SZAS-15 | Bacteria | PVC\|Chlamydiae\|Rhabdochlamydia | MAG | 1622 | GTDB | https://gtdb.ecogenomic.org/ |
| Uncultured Simckaniaceae bacterium M30B1 | Bacteria | PVC\|Chlamydiae\|Simkaniaceae | genome | 1769 | GTDB | https://gtdb.ecogenomic.org/ |
| Uncultured Simckaniaceae bacterium M30B2 | Bacteria | PVC\|Chlamydiae\|Simkaniaceae | genome | 1989 | GTDB | https://gtdb.ecogenomic.org/ |
| Uncultured Thermostichales MAG AL19W-Fe_13 | Bacteria | Cyanobacteria\|Thermostichales | genome | 3205 | GEMS | https://genome.jgi.doe.gov/portal/GEMs/GEMs.home.html |
| Uncultured Thermostichales MAG ATX19-S_21 | Bacteria | Cyanobacteria\|Thermostichales | genome | 4250 | GEMS | https://genome.jgi.doe.gov/portal/GEMs/GEMs.home.html |
| Uncultured Thermostichales MAG ATX19-S_50 | Bacteria | Cyanobacteria\|Thermostichales | genome | 2922 | GEMS | https://genome.jgi.doe.gov/portal/GEMs/GEMs.home.html |
| Ureaplasma parvum ATCC 700970 | Bacteria | Firmicutes | genome | 614 | NCBI | https://www.ncbi.nlm.nih.gov/genbank/ |
| Uronema belkae | Eukaryota | Viridiplantae | transcriptome | 7496 | 1KP | https://db.cngb.org/onekp/ |
| Vaccinium darrowii 700 v12 | Eukaryota | Viridiplantae | genome | 36717 | Phytozome | https://phytozome-next.jgi.doe.gov/ |
| Verminephrobacter eiseniae EF01_2 | Bacteria | Proteobacteria\|Betaproteobacteria | genome | 4947 | NCBI | https://www.ncbi.nlm.nih.gov/genbank/ |
| Vibrio cholerae N16961 | Bacteria | Proteobacteria | genome | 3835 | NCBI | https://www.ncbi.nlm.nih.gov/genbank/ |
| Vigna unguiculata 469 v11 | Eukaryota | Viridiplantae | genome | 42287 | Phytozome | https://phytozome-next.jgi.doe.gov/ |
| Vitis vinifera 457 V21.fa | Eukaryota | Viridiplantae | genome | 55564 | Phytozome | https://phytozome-next.jgi.doe.gov/ |
| Volvox carteri 317 v21 | Eukaryota | Viridiplantae | genome | 16075 | Phytozome | https://phytozome-next.jgi.doe.gov/ |
| Vulcanisaeta distributa DSM14429 | Archaea | Crenarchaeota | genome | 2493 | NCBI | https://www.ncbi.nlm.nih.gov/genbank/ |
| Vulcanisaeta moutnovskia 768-28 | Archaea | Crenarchaeota | genome | 2320 | NCBI | https://www.ncbi.nlm.nih.gov/genbank/ |
| Waddlia chondrophila WSU 86-1044 | Bacteria | PVC\|Chlamydiae\|Chlamydiaceae | genome | 1956 | NCBI | https://www.ncbi.nlm.nih.gov/genbank/ |
| Wigglesworthia glossinidia | Bacteria | Proteobacteria | genome | 617 | NCBI | https://www.ncbi.nlm.nih.gov/genbank/ |
| Wolbachia endosymbiont D.melanogaster | Bacteria | Proteobacteria\|Alphaproteobacteria\|Rickettsiales\|Anaplasmatacea | genome | 1276 | GTDB | https://gtdb.ecogenomic.org/ |
| Wolbachia endosymbiont O.ochengi | Bacteria | Proteobacteria\|Alphaproteobacteria\|Rickettsiales\|Anaplasmatacea | genome | 800 | GTDB | https://gtdb.ecogenomic.org/ |
| Wolbachia endosymbiont TRS | Bacteria | Proteobacteria\|Alphaproteobacteria\|Rickettsiales\|Anaplasmatacea | genome | 805 | NCBI | https://www.ncbi.nlm.nih.gov/genbank/ |
| Wolinella succinogenes DSM_1740 | Bacteria | Proteobacteria\|Deltaproteobacteria | genome | 2042 | NCBI | https://www.ncbi.nlm.nih.gov/genbank/ |
| Xanthobacter autotrophicus Py2 | Bacteria | Proteobacteria\|Alphaproteobacteria\|Hyphomicrobiales | genome | 5035 | NCBI | https://www.ncbi.nlm.nih.gov/genbank/ |
| Xanthomonas campestris ATCC_33913 | Bacteria | Proteobacteria | genome | 4181 | NCBI | https://www.ncbi.nlm.nih.gov/genbank/ |
| Xenococcus sp. PCC 7305 | Bacteria | Cyanobacteria\|Group C | genome | 5329 | JGI | https://genome.jgi.doe.gov/portal/ |
| Xylella fastidiosa 9a5c | Bacteria | Proteobacteria | genome | 2832 | NCBI | https://www.ncbi.nlm.nih.gov/genbank/ |
| Yersinia enterocolitica 8081 | Bacteria | Proteobacteria | genome | 4051 | NCBI | https://www.ncbi.nlm.nih.gov/genbank/ |
| Zea mays V4 | Eukaryota | Viridiplantae | genome | 131484 | Phytozome | https://phytozome-next.jgi.doe.gov/ |
| Zostera marina 668 V31 | Eukaryota | Viridiplantae | genome | 21483 | Phytozome | https://phytozome-next.jgi.doe.gov/ |
| Zymomonas mobilis ZM4 | Bacteria | Proteobacteria\|Alphaproteobacteria\|Sphingomonadales | genome | 2041 | NCBI | https://www.ncbi.nlm.nih.gov/genbank/ |
