## Supplemental Figures for "Starch granule initiation doesn’t require a starch synthase 4 isoform in *Chlamydomonas reinhardtii*"

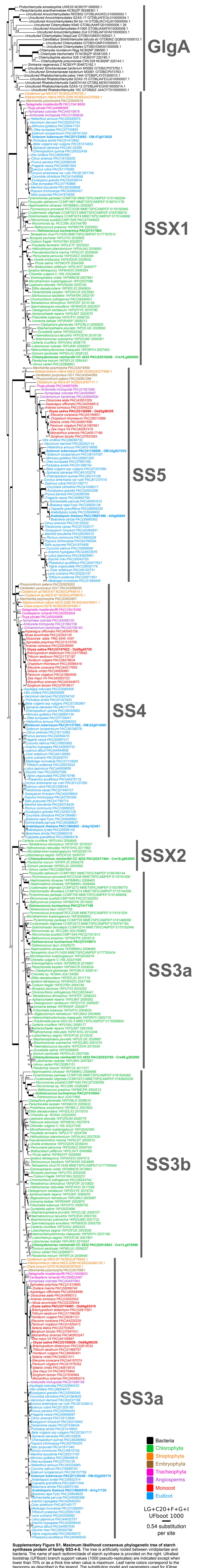


**Supplementary Figure S1. Maximum likelihood consensus phylogenetic tree of starch synthases protein of family SS3-4-5.** The tree is artificially rooted between viridiplantae and bacteria. The name of each isoform/clade of starch synthase is provided on the right. Ultrafast bootstrap (UFBoot) branch support values (1000 pseudo-replicates) are indicated except when lower than 70% or as a thick line when value is maximum. Leaf name colors correspond to the taxonomy as described in the legend. The substitution model and parameters used for IQtree reconstruction as well as the branch length scale are provided at the bottom right. Protein identifiers specify the source and the reference separated by a pipe symbol. Species used as genomic reference are in bold and the identifiers of the locus encoding the corresponding protein are indicated.

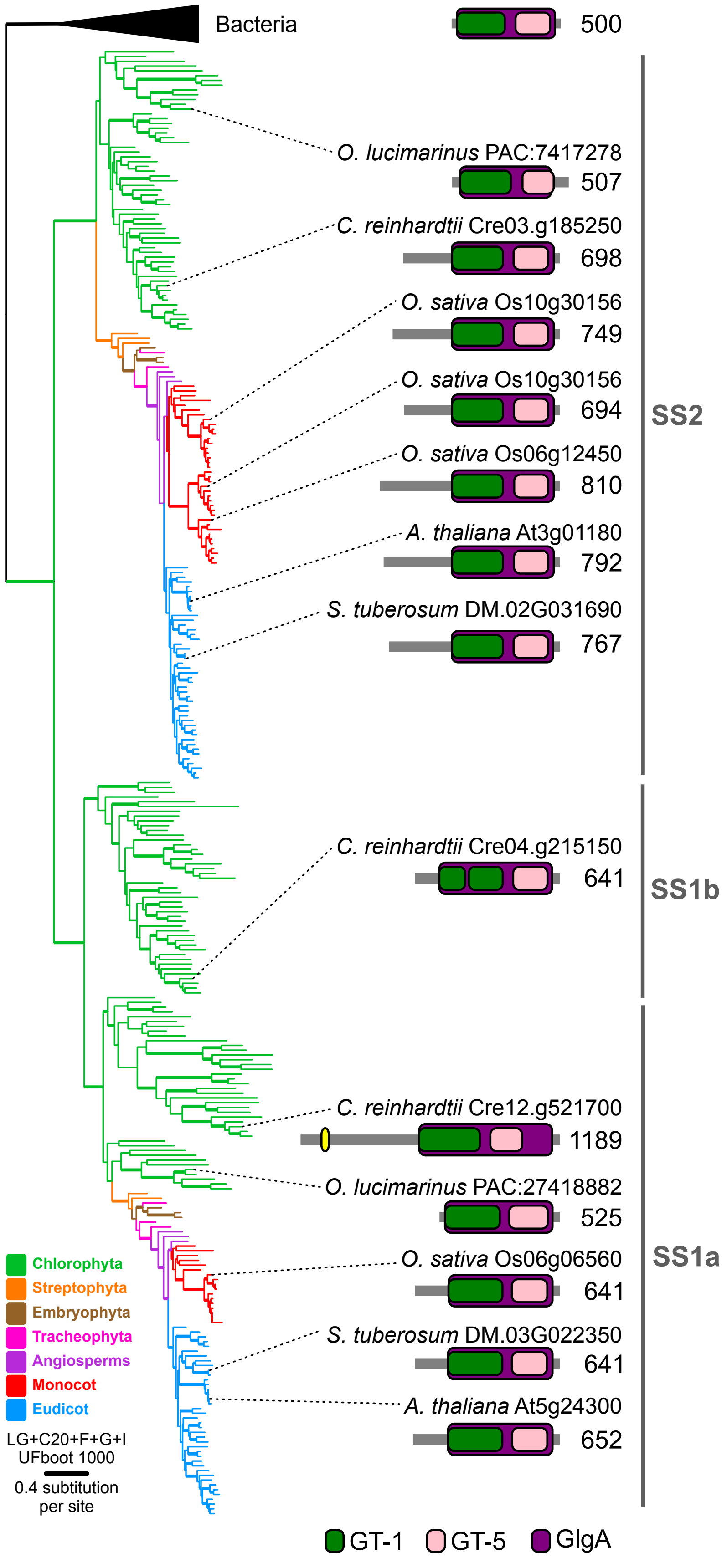


**Supplementary Figure S2. Phylogenetic and conserved domain analysis of starch synthases** isoforms belonging to family SS1-2. The maximum-likelihood tree on the left was reconstructed using IQ-tree and is artificially rooted between bacteria and viridiplantae. The name of each isoform/clade of starch synthase is provided on the right. Branch colors correspond to the taxonomy as described in the legend. Branch supports, estimated using 1000 UFBoot bootstrap pseudo-replicates, are depicted by a thick line if over or equal to 99%. Evolutionary model and branch length scale are also provided. The corresponding detailed tree with all species names, protein references and support values is available in supplementary figure S3, in the present file. Sketches of conserved domain organization are provided for a selection of proteins in each isoform family; conserved domain names and colors are detailed in the legend; numbers correspond to the length of each protein.

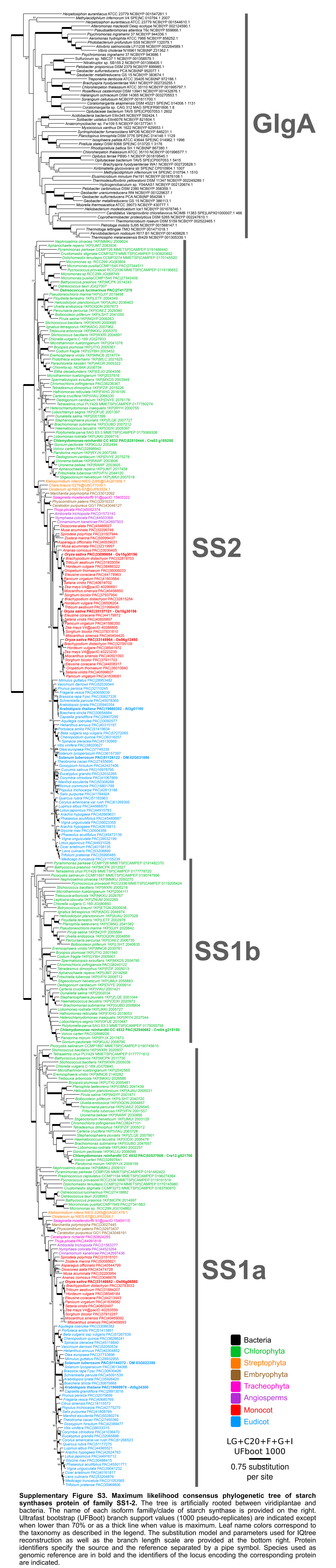


**Supplementary Figure S3. Maximum likelihood consensus phylogenetic tree of starch synthases protein of family SS1-2.** The tree is artificially rooted between viridiplantae and bacteria. The name of each isoform familly/clade of starch synthase is provided on the right. Ultrafast bootstrap (UFBoot) branch support values (1000 pseudo-replicates) are indicated except when lower than 70% or as a thick line when value is maximum. Leaf name colors correspond to the taxonomy as described in the legend. The substitution model and parameters used for IQtree reconstruction as well as the branch length scale are provided at the bottom right. Protein identifiers specify the source and the reference separated by a pipe symbol. Species used as genomic reference are in bold and the identifiers of the locus encoding the corresponding protein are indicated.

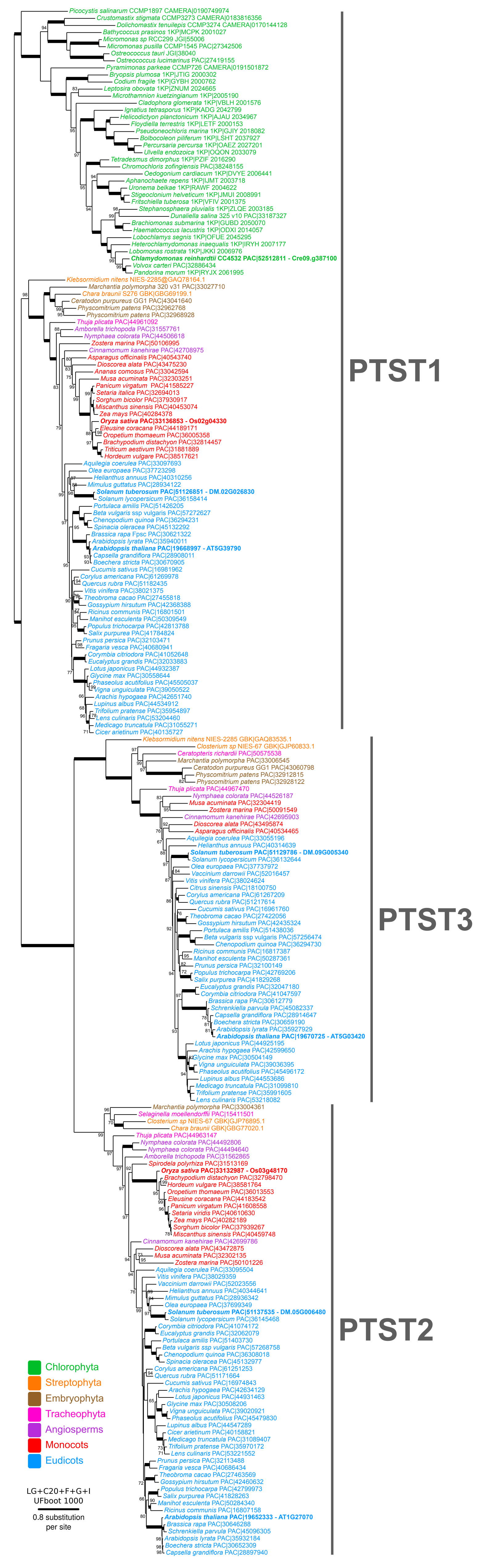


**Supplementary Figure S4. Maximum likelihood consensus phylogenetic tree of PTST-like proteins.** The tree is artificially rooted between chlorophyta and streptophyta. The name of each isoform/clade is provided on the right. Ultrafast bootstrap (UFBoot) branch support values (1000 pseudo-replicates) are indicated except when lower than 70% or as a thick line when value is maximum. Leaf name colors correspond to the taxonomy as described in the legend. The substitution model and parameters used for IQtree reconstruction as well as the branch length scale are provided at the bottom right. Protein identifiers specify the source and the reference separated by a pipe symbol. Species used as genomic reference are in bold and the identifiers of the locus encoding the corresponding protein are indicated.

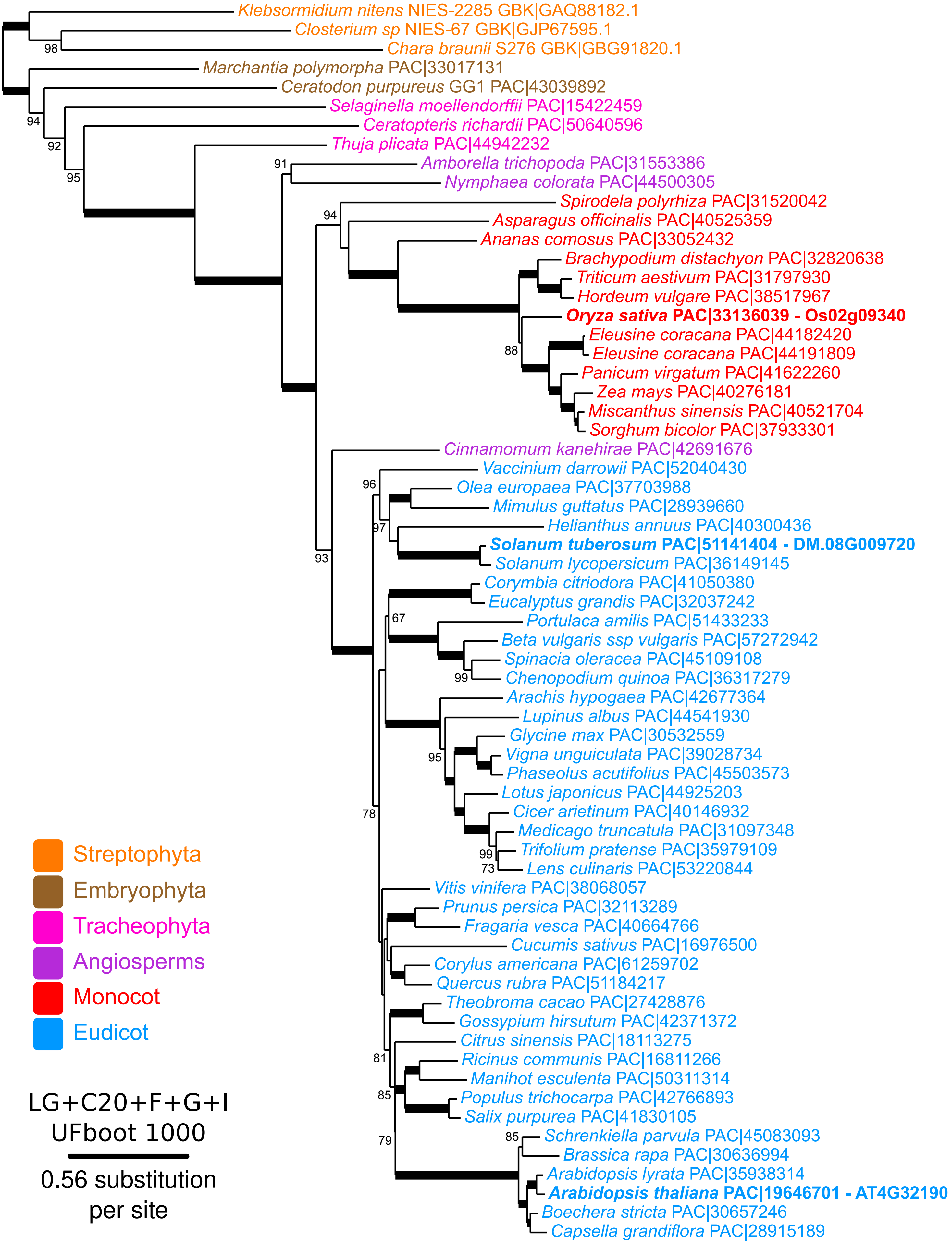


**Supplementary Figure S5. Maximum likelihood consensus phylogenetic tree of PII1 proteins.** The tree is artificially rooted between charophyceae and other streptophyta. Ultrafast bootstrap (UFBoot) branch support values (1000 pseudo-replicates) are indicated except when lower than 70% or as a thick line when value is maximum. Leaf name colors correspond to the taxonomy as described in the legend. The substitution model and parameters used for IQtree reconstruction as well as the branch length scale are provided at the bottom right. Protein identifiers specify the source and the reference separated by a pipe symbol. Species used as genomic reference are in bold and the identifiers of the locus encoding the corresponding protein are indicated.

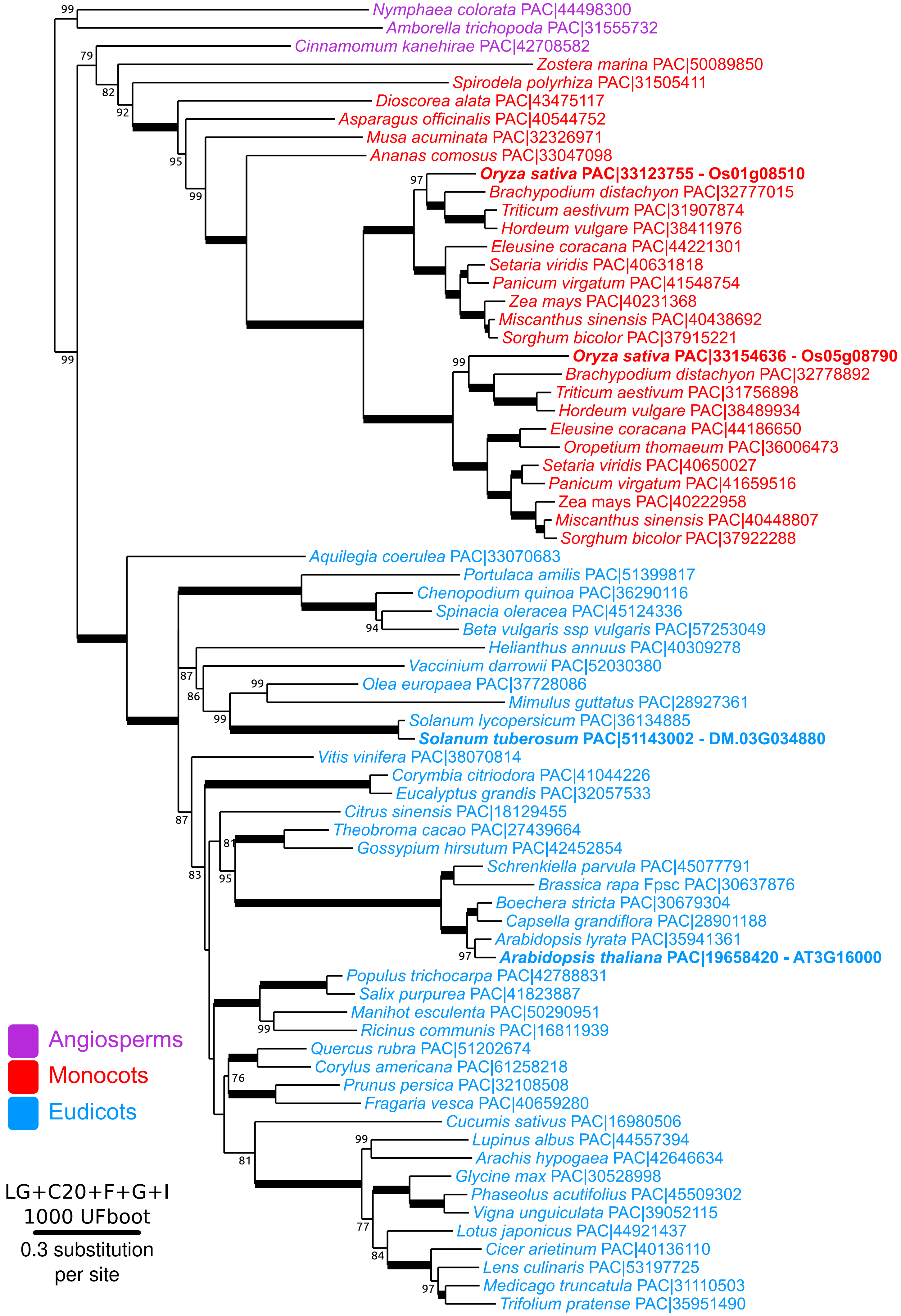


**Supplementary Figure S6. Maximum likelihood consensus phylogenetic tree of MFP1 proteins.** The tree is artificially rooted. Ultrafast bootstrap (UFBoot) branch support values (1000 pseudo-replicates) are indicated except when lower than 70% or as a thick line when value is maximum. Leaf name colors correspond to the taxonomy as described in the legend. The substitution model and parameters used for IQtree reconstruction as well as the branch length scale are provided at the bottom right. Protein identifiers specify the source and the reference separated by a pipe symbol. Species used as genomic reference are in bold and the identifiers of the locus encoding the corresponding protein are indicated.

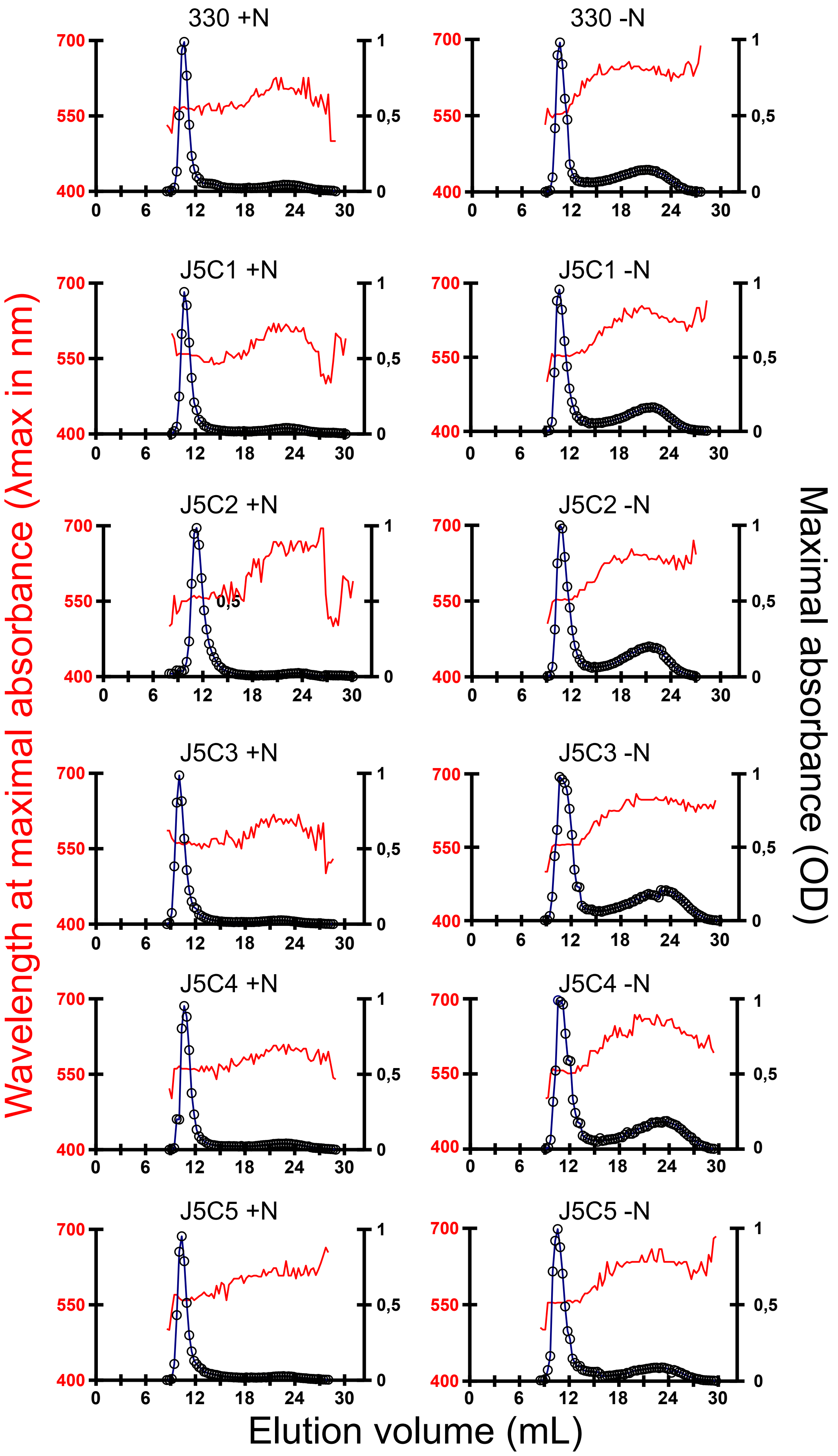


**Supplementary Figure S7: Separation of amylopectin and amylose by CL2B-sepharose chromatography.** The optical density (open circles) was measured for each 300 µL fraction at lmax (red thin line). The sample was loaded on the same column setup described by Delrue et al. (1992). Starches from the wild-type strain 330 and five complemented strains (J5C) were extracted from both mixotrophic (+N; left panel) and nitrogen-deprived cultures (-N; right panel). Quantification of amylose and amylopectin ratios were obtained by pooling amylopectin and amylose fractions separately and measuring the amount of glucose through the standard amyloglucosidase assay.
